# Dynamics of sleep, feeding, and metabolic homeostasis in *Drosophila* ensheathing glia, astrocytes, and neurons

**DOI:** 10.1101/2022.07.07.499175

**Authors:** Andres Flores-Valle, Ivan Vishniakou, Johannes D. Seelig

## Abstract

Sleep is critical for homeostatic processes in the brain, including metabolism and waste removal. Here, we identify brain-wide, locally acting sleep homeostats for the short, naturally occurring sleep bouts of *Drosophila* in the two major classes of glia that arborize inside the brain, astrocytes and ensheathing glia. We show that glia surround respiratory tracheal tubes, that the metabolic gas carbon dioxide, changes in pH, or behavioral activity, all induce long lasting calcium responses, and that astrocytes and glia show circadian calcium modulations. Glia describe sleep homeostasis in behaving flies more faithfully than previously identified sleep circuits in the central complex, but a subset of neurons in the fan-shaped body is important for feeding homeostasis. Local optogenetic activation of astrocytes or ensheathing glia is sufficient to induce sleep. Together, glia calcium levels can be modeled as homeostatic controllers of metabolic activity, thus establishing a link between metabolism and sleep.

## Introduction

Sleep is important for multiple homeostatic processes in the brain, including the reconfiguration of networks for learning and memory^1,2^, metabolism^3–5^, for the removal of waste products^4,5^, regulation of ion concentrations^6^, or as a response to disease^7,8^. In particular glia are important for many of these processes^7,9,10^. Of the different classes of *Drosophila* glia^11,12^, astrocytes and ensheathing glia arborize inside the neuropil, where axons and dendrites form synapses, numbering around 4000 and 5000 cells^11^, respectively, compared to about 130’000 neurons^13^. Both astrocytes and glia are important for circadian as well as homeostatic sleep regulation^7,10,14,15^.

Sleep bouts in Drosophila^16–20^ are distributed throughout the day and night with a mean duration of only around 20 minutes^20–22^. How commonly used approaches for investigating sleep homeostasis in the fly, which typically rely on many hours of sleep deprivation, are related to these short sleep bouts, is not known. Unlike in mice, where calcium imaging links glia dynamics to sleep-wake cycles^23–30^, in flies the dynamics of glia and also of most sleep-related neurons have yet to be described in behaving animals.

Here, we use two-photon^31^ calcium imaging in flies navigating in virtual reality over multiple days to investigate the dynamics of ensheathing glia (EG), astrocyte-like glia (AL, in the following also referred to as astrocytes), and sleep-related neuron in the central complex^14,15,32–36^. We find that astrocytes and ensheathing glia show both calcium-dependent circadian modulation and homeostatic dynamics. Circadian activity is stronger in astrocytes, and homeostatic activity is stronger in ensheathing glia. Glia monitor sleep and wakefulness more reliably than previously identified sleep-related neurons in the central complex, a subset of which, dFB neurons^32^, we find to represent feeding homeostasis. Calcium dynamics does not only encode sleep need, but local activation of ensheathing glia or astrocytes in central brain compartments using optogenetics is sufficient to induce sleep.

To understand the mechanism through which glia sense sleep need, we take advantage of the connectome^37^, and show that trachea are embedded inside ensheathing glia, suggesting a role of glia in gas metabolism. Consistent with this idea, we show that exposure to CO_2_, which changes pH^38–42^, or direct optogenetic perturbations of pH^43,44^, lead to homeostatic calcium responses in glia and astrocytes, but not in neurons. Additionally, both CO_2_ exposure and optogenetics induce sleep-like behavior. We finally model glia calcium activity as a controller that counteracts local pH changes to maintain a physiological setpoint. These results link sleep homeostasis to metabolism in behaving flies.

### Time scale of sleep homeostasis

Walking flies show a distribution of bouts of rest or sleep of different lengths during the day and night^16,17,21,22^ with a mean bout duration of around 20 minutes^21,22^. We confirmed that this short sleep duration is not the result of minor fly movements that could only briefly interrupt overall longer sleep bouts. We quantified the distribution of sleep bouts in flies walking in a rectangular chamber by tracking them with a camera (Fig. 1a, b, Supplementary Fig. S1a, b, see Methods). We removed shorter periods of movement of varying lengths between consecutive bouts of immobility (Fig. 1c) and computed the resulting distribution of sleep bouts. Even when filtering out movement periods of 2 minutes, 90% of sleep bouts were still shorter than 50 minutes during the day and night (Fig. 1 d), with even shorter sleep bouts in constant darkness (Supplementary Fig. S1c), consistent with previous results^21,22^.

**Figure 1.**
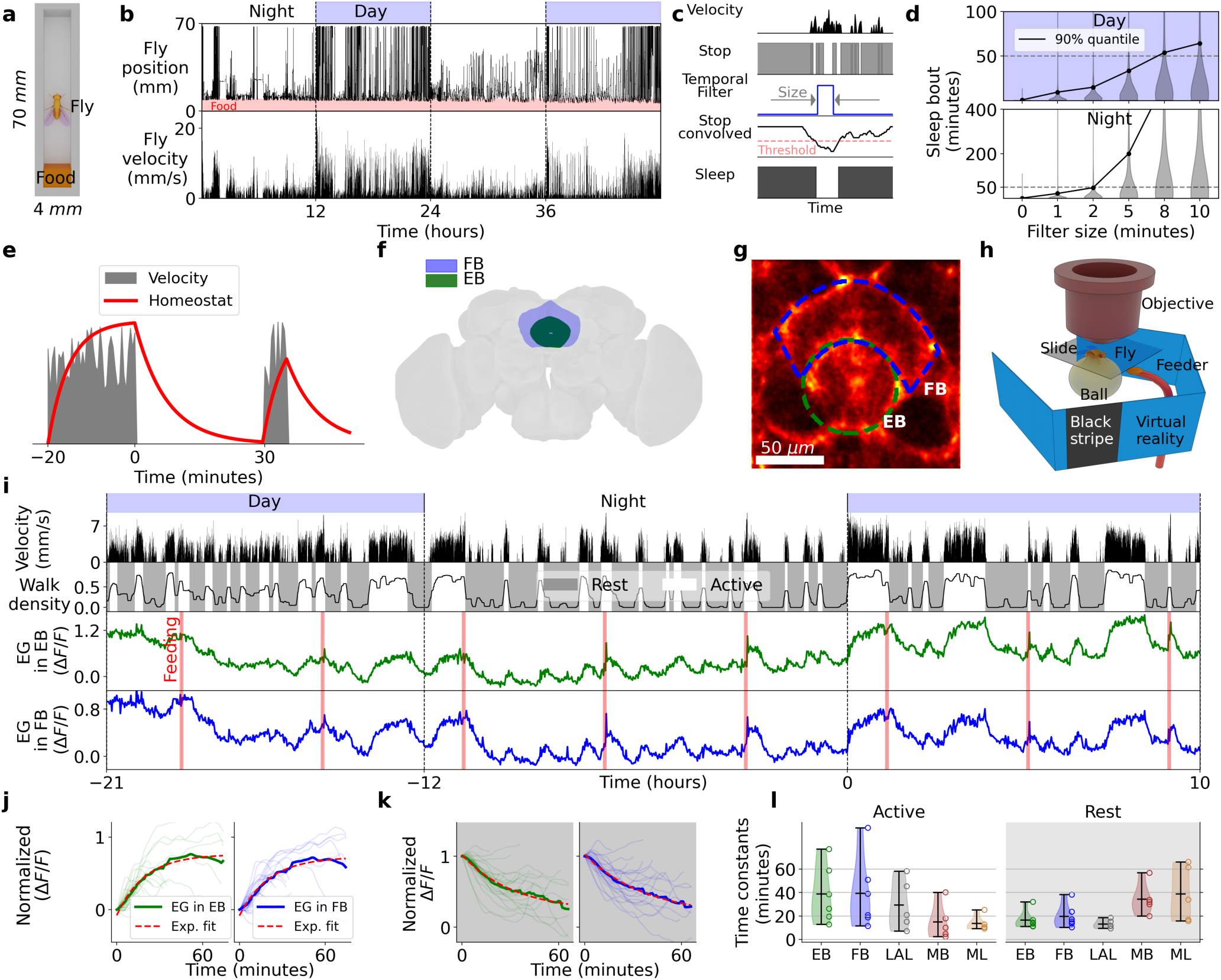
Behavior in freely moving flies and long-term calcium imaging in ensheathing glia. **a** The position of flies walking in a rectangular chamber with food is tracked from the top with a camera. **b** Position along the rectangular chamber and velocity over two days for a single fly. **c** Velocity (first row) is thresholded to find bouts of immobility (’stop’, second row). A temporal filter (third row) is convolved with the stop state of the fly (fourth row) and a threshold is used to characterize sleep (fifth row). **d** Distribution of sleep bouts for a total of 15 flies during the day and night as a function of filter size. **e** Schematic of expected dependence between sleep homeostat signal and behavior, integrating time awake (rising, with non-zero velocity higher, shown in grey) and time sleeping (decreasing, zero velocity). **f** EB and FB in the *Drosophila* brain. **g** Ensheathing glia express jGCaMP8m and regions of interest (ROIs) for analyzing fluorescence in EB (green) and FB (blue). **h** Setup for long-term imaging: a fly is glued to a cover slide and navigates on an air supported ball in VR under the microscope objective. **I** Long-term recording of a single fly over 31 hours. Top row: day and night cycle in VR. Second row from top: absolute ball velocity. Third row: walk density obtained by low-pass filtering (cut-off period of 6 minutes) of ’walk’ (walk: 1 if the fly has non-zero velocity in 1 second bins, 0 otherwise). Active and rest states were defined when the walk density was above or below a threshold (see Methods). Fourth and fifth row: calcium activity in EB and FB ensheathing glia over time, respectively. Red lines indicate time of feeding. **j** Single traces of increasing normalized calcium activity (see Methods) during active states (thin lines), average (thick line), and exponential fit (red) for EB (left) and FB (right). **k** Same as j but during rest state, fitted with exponentials (red). **l** Time constants for exponential fitting during active (white region) and rest states (grey region) for EB, FB, LAL, MB, and midline (ML). Violin plots represent the distribution of time constants for 6 flies in the EB and FB, 5 flies in the LAL, and 5 flies in the MB and ML (circles). Short horizontal black lines show maximum, mean, and minimum values in descending order.

A sleep homeostat needs to keep track of time spent awake and sleeping^45–47^. This is typically modeled with two single exponential functions approaching an upper threshold for tracking time spent awake, and a lower threshold for measuring time spent sleeping (Fig. 1e). The time constants of the two exponentials determine average wake and sleep bout durations^45–47^. Thus, based on behavior the dynamics, or time constants, of a brain signal that keeps track of naturally occurring sleep and wake bouts in the fly are expected to lie in the range of several minutes or at most tens of minutes.

### Homeostatic activity of ensheathing glia

Both astrocytes and ensheathing glia are important for sleep^10,12,14,48–52^, but whether and how these different glia types contribute to the short sleep bouts of the fly is not know. Ensheathing glia form diffusion barriers around different brain compartments^11,12^. Such a structural arrangement would be fitting for monitoring the accumulation of metabolites and therefore for integrating neural activity in brain compartments, as expected of a sleep homeostat^47^. Astrocytes, which tile the brain more homogeneously^11^ could sense neural activity more locally^53,54^.

To investigate calcium dynamics of glia and astrocytes in behaving animals we used two-photon calcium imaging. In these experiments, flies walked on an air supported ball in a virtual reality setup in closed-loop with a single dark stripe on a bright background during the day and in darkness during the night (12 hours light/dark cycle)^55^. We first expressed the calcium indicator jGCaMP8m^56^ in ensheathing glia using the GAL4 line R56F03^11,49,57^ and imaged different compartments of the central brain, for example the ellipsoid body (EB) and fan-shaped body (FB, Fig. 1f, g) of the central complex. We fed flies in the imaging setup using an automated feeder (Fig. 1h) every four hours^55,58^, which induced epochs of continuous walking activity (often before feeding) or rest and sleep (often after feeding, Fig. 1i). Under these conditions, ensheathing glia in different brain areas including the EB, FB, the lateral accessory lobe (LAL, Supplementary Fig. S2a, b, mushroom bodies (MB), and midline (ML), Supplementary Fig. S2c), showed pronounced calcium fluctuations (Fig. 1i, Supplementary Figs. S3-S5, and Supplementary Videos S1-S6 for EB and FB).

In a first analysis we distinguished two behavioral states based on the fly’s walking activity (see sections below for a more detailed, model-based analysis): ’walk’ (fly is walking) and ’stop’ (fly is standing still as assessed based on ball velocity, second row in Fig. 1i, see Methods for details). We then selected epochs of at least 10 minutes during which the fly was walking most of the time (active, third row in Fig. 1i) or immobile most of the time (rest, third row in Fig. 1i) by thresholding the ’walk density’, obtained by low-pass filtering of ’walk’ (see Methods).

Fluorescence traces were averaged over the selected active and rest epochs (Figs. 1j, k, Supplementary Figs. S6-S8). The averaged fluorescence could be fitted with exponentials (see Methods), the expected time course of sleep homeostasis^59,60^ (red line in Figs. 1j, k, Supplementary Figs. S6-S8). We verified that saturation was not due to indicator saturation (see Methods and Supplementary Fig. S10). The resulting rise and decay time constants of the exponential fits are shown in Fig. 1l for 6 flies for EB and FB, 5 flies for LAL, and 5 flies for MB and ML, respectively, and lie in the range expected based on behavior (Fig. 1d, e). Dynamics in the EB, FB, and LAL were similar, whereas the MB and ML showed faster dynamics during active, and slower dynamics during rest epochs, respectively (see also Supplementary Fig. S2-S7, Supplementary Information, and Supplementary Figs. S19, S20 for MB, and ML). Ensheathing glia also showed fluctuations at circadian timescales, with overall higher activity during day than night, with however smaller changes in amplitude than those resulting from homeostatic activity (Fig. 1i, see below).

**Table S1.** Percentage of time sleeping per hour for each fly according to the different definitions used in the paper.

| Fly | Sleep<br>using ball<br>velocity (stop)<br>(% per hour) | Sleep<br>using walk density<br>(rest)<br>(% per hour) | Sleep<br>with 5 minutes<br>threshold<br>(% per hour) |
| --- | --- | --- | --- |
| 1 | 50.0 | 49.1 | 18.7 |
| 2 | 56.8 | 51.5 | 28.8 |
| 3 | 68.1 | 61.0 | 33.6 |
| 4 | 70.8 | 51.6 | 30.4 |
| 5 | 57.6 | 46.0 | 19.1 |
| 6 | 76.4 | 61.2 | 31.0 |

### Astrocytes show circadian and homeostatic modulation

To investigate whether astrocytes showed similar dynamics as ensheathing glia during behavior, we expressed the calcium indicator jGCaMP8m in 86E01-GAL4^50^. Astrocytes showed calcium signals with stronger circadian modulation than ensheathing glia (Fig. 2a) and also showed homeostatic fluctuations superimposed on the circadian modulations (Fig. 2a). Time constants for these modulations were first computed as before (see below for a more detailed, model-base analysis) by separating epochs where flies were mostly walking or sleeping (Fig. 2b, see Methods). The time constants for astrocyte homeostatic signals resulting from this analysis were slightly faster than those of ensheathing glia (Fig. 2c).

**Figure 2.**
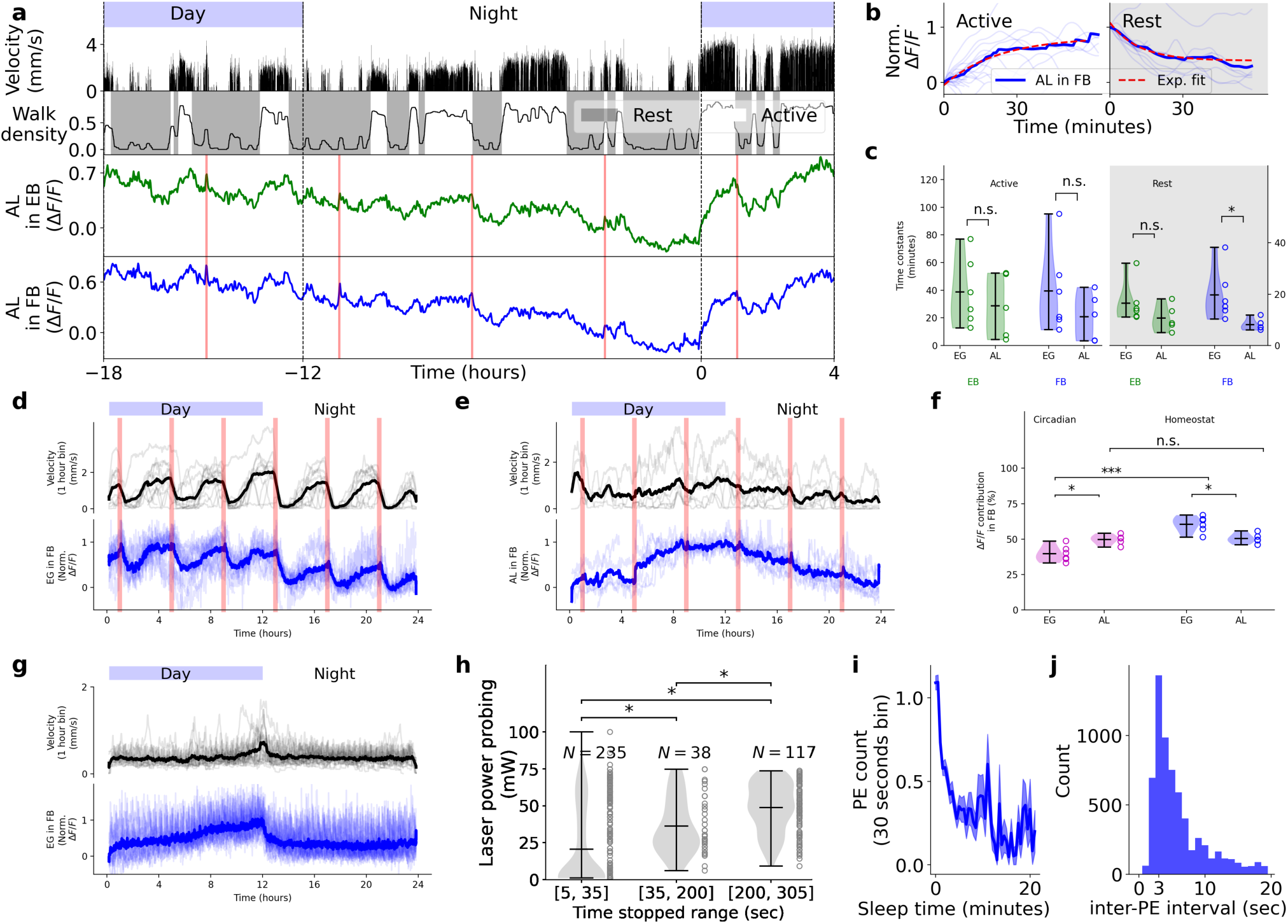
Calcium dynamics in astrocytes, circadian activity and sleep behavior during long term imaging experiments. **a** Recording of astrocytes-like glia (AL, or astrocytes) in the EB and FB for 22 hours (same as 1i). **b** Same as 1j and k but for astrocytes in the FB. **c** Time constants of ensheathing glia and astrocytes during wake and sleep in the EB and FB during active and rest behaviors (same as 1l). **d** Average velocity (second row) and activity of ensheathing glia in the FB (third row) over 24 hours. Thin lines represent individual days for each fly, while thick lines represent the average. Red vertical lines represent the feeding times. **e** Same as d but for astrocytes in the FB. **f** Percentage of circadian (magenta) and homeostat (blue) contributions to the activity of ensheathing glia and astrocytes in the FB (see Methods). One or three asterisks indicate statistical significance using t-test (p < 0.05 or p < 0.0005, respectively). **g** Same as d but for experiments where flies were fed every 25 minutes. **h** Laser power required to awaken the fly based on the previous durations of immobility during calcium imaging. Durations of immobility are grouped in three different time ranges; the number of probing trials is indicated at the top (for a total of 6 flies). Statistical significance was assessed using the Kolmogorov–Smirnov test. Asterisks in all panels indicate p-values < 0.05. **i** Proboscis extension (PE) count as a function of sleep time during long-term imaging experiments for 21 flies. **j** Histogram of time between proboscis extension events during sleep behavior during long-term imaging experiments (21 flies).

### Comparison of astrocytes and ensheathing glia activity

Circadian activity was weaker in ensheathing glia than in astrocytes (Fig. 2d-f). To better observe circadian oscillations in ensheathing glia largely independent of behavior, we sought to minimize the flies walking activity by keeping them well fed at all times. We fed the flies every 26 (Fig 1g, Supplementary Fig. S9) or 16 minutes (Supplementary Fig. S13), respectively, at a similar rate at which freely walking flies approached food (Supplementary Fig. S1d) for 2 minutes without interrupting the experiments using the automated feeder (Fig. 1h)^55^. The approaching feeder led to brief bouts of walking behavior at the time of feeding and corresponding, walking-induced (as described in the previous section, as well as potentially due to startling the fly, see below) spikes of calcium activity (Fig 2g, Supplementary Fig. S9, S13), but little walking activity between feeding events. Under these conditions, calcium activity in individual flies as well as averaged across flies slowly increased across the day and reset during the night with a circadian pattern independent of behavior (Fig. 2g). Overall, homeostatic fluctuations were stronger than circadian fluctuations in ensheathing glia, and also stronger than homeostatic signals in astrocytes. Circadian fluctuations were stronger in astrocytes than in ensheathing glia (Fig. 2f).

### Sleep behavior observed during imaging

The distributions of bouts of immobility during long-term imaging were similar during the day and identical during the night to those of freely moving flies (Supplementary Fig. S30a and b, respectively, see Methods). To determine whether epochs of immobility during long-term imaging experiments correspond to sleep, we tested whether the arousal threshold, the required stimulus strength to induce a behavioral response in resting flies, increased with the duration of immobility, as expected for sleep^18,61^. As a stimulus we used an infrared laser beam pointed at the fly’s abdomen. At the start of each trial of probing of the arousal threshold, we ensured that the fly was first awake by closing and opening the air stream to the air supported ball once per second for 3 seconds (see Methods), which stimulated the fly to walk. Then, using an automated control loop for monitoring ball velocity, we detected bouts of immobility. After either 30 seconds or 5 minutes of immobility, the power of the heating laser was gradually ramped up until the fly started to walk (see Methods). To ensure that flies were completely still during the detected epochs of immobility, we classified behavior recorded with a camera using machine learning in post-processing (see Methods and Supplementary Fig. S14). Removing those sections where the fly was grooming resulted in the final bouts of immobility between 5 seconds and 5 minutes. A representative trial of threshold probing is shown in Supplementary Fig. S15a. Longer periods of immobility required higher laser powers for a behavioral response (Fig. 2h), showing that the arousal threshold increases with the duration of immobility, a feature of sleep also observed in freely walking flies^16–18^ (see Supplementary Fig. S15b-g for individual flies). Proboscis extension, another sleep-related behavior^20,62^, was detected by classifying behavior using machine learning (see Methods) and occurred more frequently at the beginning of sleep bouts during long-term imaging (Fig. 2i), at similar frequencies as previously described (Fig. 2j)^20^.

### Ensheathing glia integrate behavioral effort

Since ensheathing glia show stronger homeostatic signals than astrocytes (Fig. 2f), we performed a more detailed analysis of the homeostatic features of these cells. Sleep behavior was often (but not always) correlated with feeding events (red vertical lines in Fig. 1i, 2a, Supplementary Fig. S3-S5), with flies increasingly walking more with time elapsed since feeding, and sleeping more after feeding (as observed in freely walking flies^14,63^).

We first asked whether the increase in calcium in ensheathing glia was specifically due to walking, or whether other effortful behaviors could also cause similar activity changes. To address this question, we immobilized the ball for epochs of 40 minutes while the fly was in the VR setup, and recorded behavior and glia activity in the EB and FB as before. Movement of the ball was blocked in such a way that the fly continually pushed or pulled (see Methods for details). As seen in the resulting fluorescence traces (Fig. 3a and Supplementary Fig. S16), this led to a steady increase in or saturation of calcium levels similar to the dynamics observed due to walking activity with slightly faster time constants (Fig. 3b, c and Fig. 1j, l). This shows that glia calcium activity increases not only due to coordinated walking but also due to other effortful behaviors (Fig. 3c, d), as similarly observed in zebrafish^64^.

**Figure 3.**
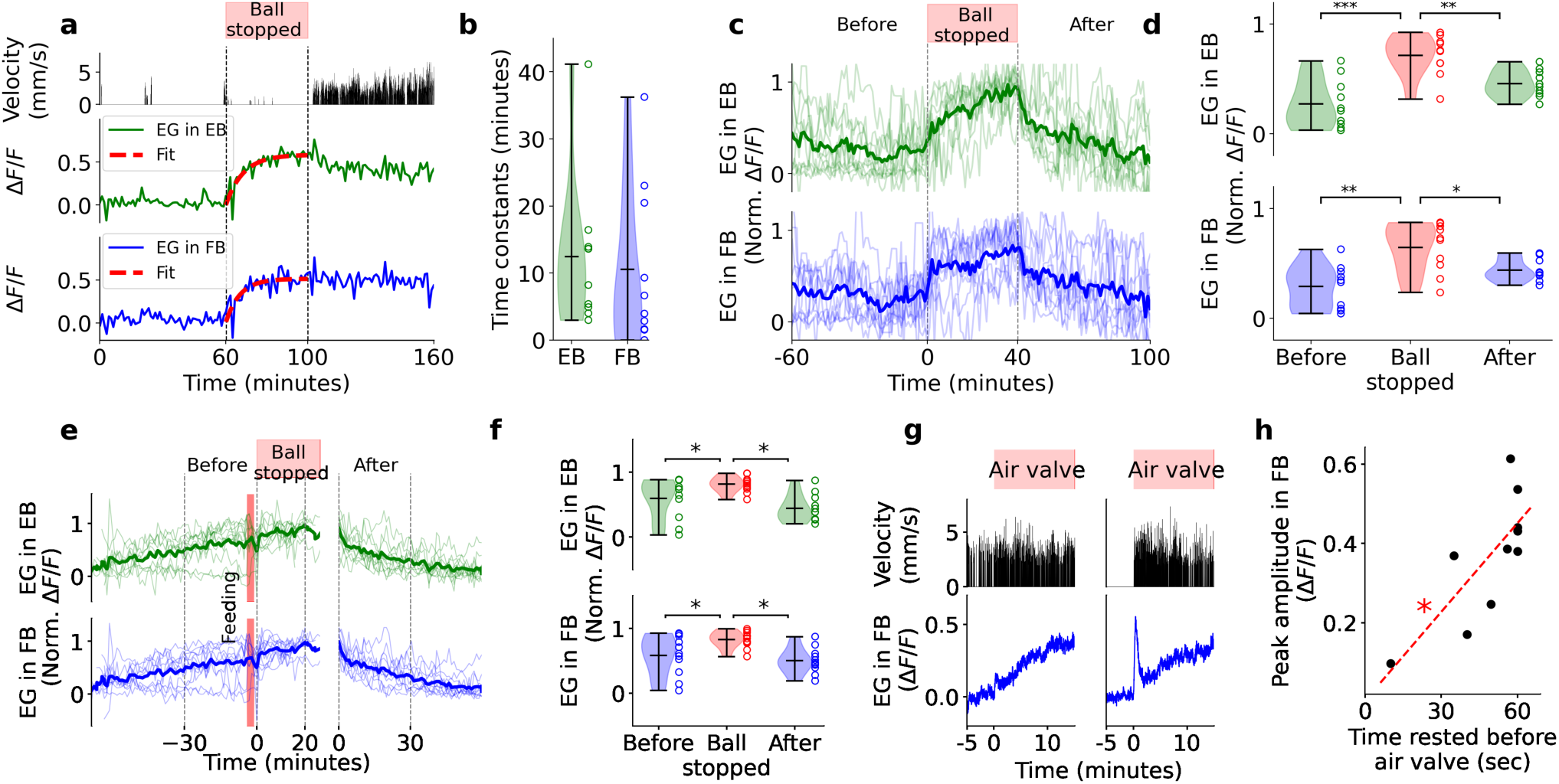
Calcium dynamics in EG increases due to effortful behavior, decreases during rest, and saturates under sleep deprivation. **a** Calcium activity during effort with blocked treadmill ball. Second row: velocity of fly over time (zero while stopping the ball). Third and fourth row: EG activity and exponential fit (red) in EB and FB, respectively. **b** Time constants of exponential fits while ball was stopped for *N* = 10 trials in 5 flies. **c** Normalized fluorescence traces of EG activity in EB and FB before, during, and after ball was stopped (top row). Second and third row: single trials (thin lines) and average (thick lines) in EB and FB, respectively. **d** Normalized fluorescence levels (y-axis shared with panel g) before, during, and after ball stopping (60, 40, and 60 minutes average), respectively. Statistical significance was assessed using t-test. **e** Calcium activity during effortful behavior after feeding. Normalized fluorescence traces of EG activity in EB and FB before feeding, after feeding while the ball was stopped, and after releasing the ball (for *N* = 11 trials in 5 flies). **f** Normalized fluorescence levels (y-axis shared with panel j) before feeding, after feeding while stopping the ball, and after releasing the ball (30, 26, and 30 minutes average); asterisks represent statistical significance for p-values less than 0.05 using t-test. **g** Velocity (second row) and EG activity in the FB (bottom row) while the air valve of the ball is switched on and off every second (top row, red) for two trials without and with a calcium peak at the onset of mechanical stimulation (left and right, respectively). **h** EG peak amplitude in the FB at the onset of mechanical stimulation as a function of time rested, measured 60 seconds prior the onset of ball stimulation. Red line shows linear fit, asterisk indicates statistical significance of the fit using Pearson correlation (p < 0.05).

Next, we verified that feeding was not the cause for the decay of calcium activity in those instances where flies were immobile or sleeping after feeding. We sleep deprived flies immediately after feeding by inhibiting ball movement as above (Fig. 3e and Supplementary Fig. S17). This not only suppressed resetting of calcium levels, but slightly increased calcium signals after feeding (Fig. 3e, f), demonstrating that rest or sleep, and not feeding, are responsible for the decay of calcium levels after feeding.

### Fast modulations of ensheathing glia

Ensheathing glia integrate activity over long timescales (tens of minutes) as indicated by the time constants (Fig. 1l, see also below for time constants from model fitting). Modulations of fluorescence signals were however already observed at the timescale of 1 minute when comparing changes in walking velocity with changes in high-pass filtered calcium signals in the EB and FB (DF/F, Supplementary Fig. S18, a-i and Methods). Thus, the accumulation of such faster fluctuations could lead to the integrated activity observed over longer timescales.

To investigate the transition from sleep to walking in more detail, we imaged calcium dynamics at higher time resolution (Fig. 3g, see Methods)^55^. We selected bouts where flies were immobile for several minutes, and then forced them to walk by repeatedly interrupting the air stream supporting the treadmill ball with an automated valve as above. Under these conditions ensheathing glia calcium activity showed a fast peak at walking onset, followed by the previously described slower integrating response (Fig. 3g, right side). Such calcium peaks were not observed when the fly was already walking before the onset of treadmill perturbations (Fig. 3g, left side). The peak amplitude increased when flies were sleeping longer before onset of forced walking (Fig. 3h). Potentially related signals were observed in a recent paper following electric shock in MB^65^.

### Sleep deprivation saturates glia activity

A signature of a sleep homeostat is that it saturates and plateaus under sleep deprivation^45,66^. To sleep deprive flies during long-term imaging, we periodically opened and closed the air stream supporting the ball at 1 second intervals for 6 seconds every 20 seconds, similar to sleep deprivation in freely walking flies^33,35,67^, which induced short bouts of fast walking (at least every 20 seconds). Calcium activity saturated in EB and FB (Fig. 4a and b, and Supplementary Fig. S21a) within 2 hours and plateaued (Fig. 4c), with occasional fluctuations around the saturation level (indicated by an exponential fit, red line in Fig. 4a). This experiment additionally again confirmed that behavioral activity, here walking, prevents resetting of calcium levels after feeding, similar to Fig. 3e (see Fig. 4a and Supplementary Fig. S21a for feeding events during sleep deprivation).

**Figure 4.**
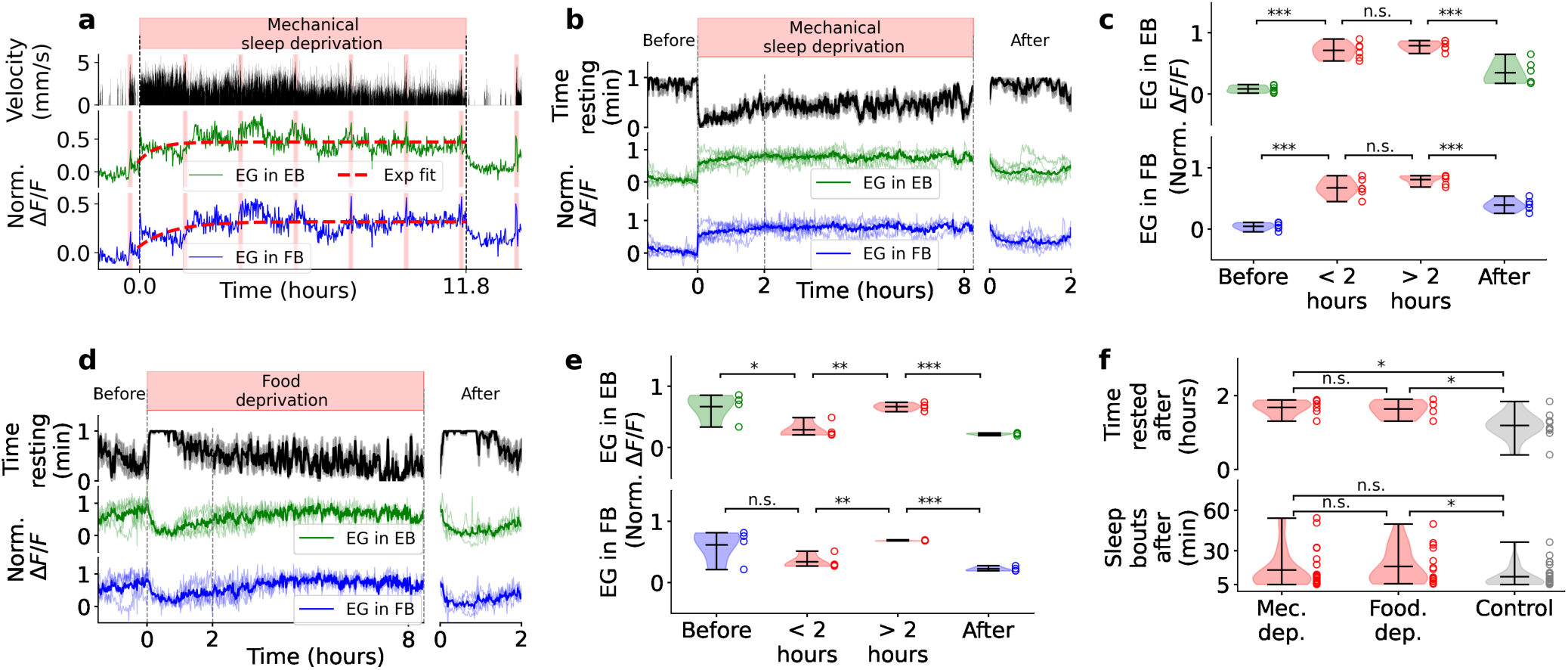
Glia activity due to sleep deprivation and effort. **a** Behavior and calcium activity in EB and FB ensheathing glia under mechanically induced sleep deprivation. Top row: time interval where sleep deprivation was applied. Second row: fly velocity. Third and fourth row: activity of glia with exponential fit for visualization of plateau level (red) during mechanical sleep deprivation for EB and FB, respectively. Vertical red lines indicate feeding events. **b** Mechanical sleep deprivation for *N* = 6 trials in 5 flies (including fly in a). Top row as in a. Second row: averaged time resting over time, calculated in bins of 1 minute. Third and fourth row: normalized fluorescence traces of ensheathing glia activity in each fly (thin lines) and average (thick line, at least 2 flies) in EB (green) and FB (blue), respectively, shown during up to 8 hours of mechanical sleep deprivation. Times of mechanical sleep deprivation varied between flies. Flies were fed during these experiments (see Supplementary Fig. S21). **c** Average fluorescence distributions before (over 1.5 hours), during the first 2 hours of sleep deprivation, between 2 and 8 hours of sleep deprivation, and after (2 hours). **d** Same as b but during food deprivation for *N* = 4 trials in different flies. Top row shows the time interval (in red) between two consecutive feeding events. Times of food deprivation varied between flies (see Supplementary Fig. S21b). **e** Same as c but for food deprivation. **f** Top row: time rested over 2 hours for each trial after mechanical sleep deprivation, food deprivation, and for control trials (*N* = 10 in 6 flies). Bottom row: distribution of sleep bouts (longer than 5 minutes) over the 2 hours after mechanical sleep deprivation, starvation induced sleep deprivation, and in control flies.. Asterisks in all panels indicate statistical significance (p < 0.05) using t-test (panels c,e and the top row of f) and Kolmogorov–Smirnov test in the bottom row of f.

For comparison, we also used food deprivation to sleep deprive flies. Absence of food and starvation induce hyperactivity or foraging behavior and prevent sleep^14,68–70^ and in some flies resulted in long epochs of continued fast walking. Single trials and average activity of 4 flies that were continuously walking for at least 3 hours following food deprivation with fluorescence traces normalized for comparison, are shown in Fig. 4d (see Supplementary Fig. S21b for individual trials). Starvation-induced walking produced a slow increase of glia calcium activity (Fig. 4d), and, similar to mechanical sleep deprivation, calcium activity saturated after 2 hours (Fig. 4e).

Sleep deprivation for extended periods of time leads to rebound sleep in flies^14–17^. The amount of rebound sleep however depends on the method of sleep deprivation. Sleep deprivation which avoids strong mechanical perturbation results in less than one hour of mean rebound sleep after 12 hours of sleep deprivation^67^, consistent with the dynamics of a sleep homeostat in the order of tens of minutes. After the offset of mechanically or food induced sleep deprivation, flies became less active (right side of Fig. 4b and d), and glia activity returned to lower levels. Flies displayed rebound sleep and showed an average of 1.68 and 1.64 hours of immobility in the two hours after the offset of mechanical and starvation induced sleep deprivation, respectively, compared to 1.19 hours in control flies (Fig. 4f, see Methods). The difference between sleep-deprived and control flies (around 30 minutes more in sleep-deprived flies on average) is consistent with the time that it takes to reset glia activity to baseline levels (Fig. 1l). Additionally, mechanical sleep deprivation resulted in more sleep bouts with a duration of more than 5 minutes (Fig. 4f). Starvation induce sleep deprivation is thought to not induce a homeostatic response compared to mechanical sleep deprivation^71^, but camera based methods might be required to detect the smaller amounts of rebound that we observe here^21,67^.

We additionally tested whether calcium responses could result from accumulation of glutamate, since ensheathing glia play a role in glutamate homeostasis^12,72^. However, time constants for glutamate were faster than those for calcium (Supplementary Fig. S26c).

### Homeostat model describes glia activity

Homeostatic activity can be modeled as exponentially decaying during sleep and exponentially approaching a saturation level during wakefulness (Fig. 1 e)^59,60,66^. We therefore quantified whether ensheathing glia calcium levels integrate behavioral history, that is, wake and sleep, according to such homeostatic dynamics. We fitted a differential equation with two time constants for charging and resetting of the homeostat, respectively, to glia calcium activity dependent on behavioral state (two-state model, see Methods). As illustrated in Fig. 5a, the behavioral states, here ’walk’ and ’stop’, are passed to the model, which represent the homeostat, and differentially increases or decreases activity with time constants characterizing the exponential dynamics. After fitting (see Methods), the homeostat model^46,73^ describes ensheathing glia activity over the time course of the experiments based on behavior (31 hours in Fig. 5b, see also Supplementary Figs. S22, S23, and S24). Similarly, astrocyte calcium dynamics can be fitted with a homeostatic model based on the fly’s behavioral state (Fig 5c and Supplementary Fig. S25). The model can therefore fit activity in EB and FB for both astrocytes and ensheathing glia based on behavior, as indicated by the absence of statistical difference in *L*2 error between the model fit and activity (Fig. 5d, also see Supplementary Information for models taking into account 3 and 7 different behavioral states, respectively).

**Figure 5.**
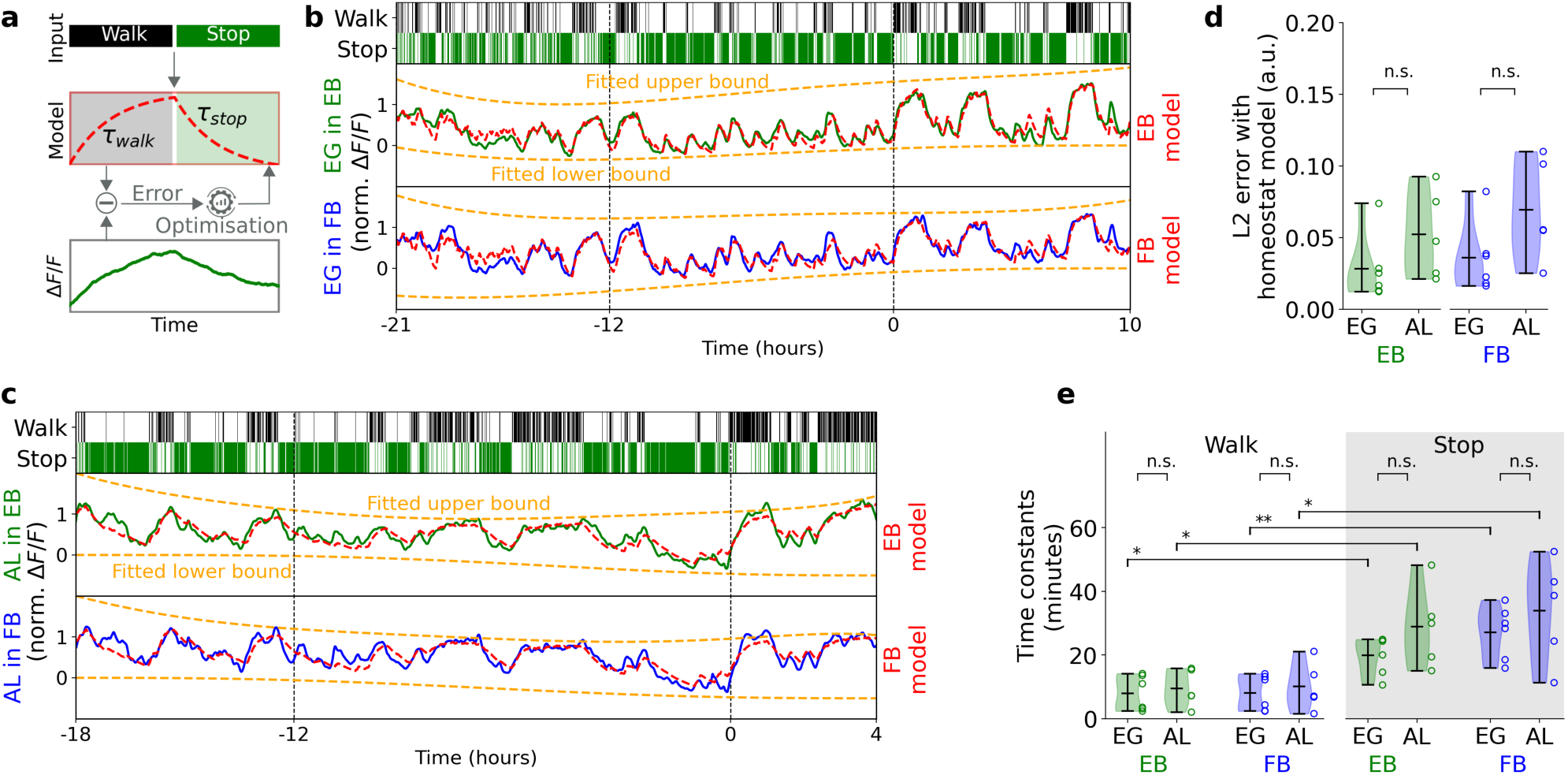
Two-state model for calcium dynamics. **a** Schematic of model fitting approach for calcium imaging: behavior (walk, stop) is integrated as model input with corresponding time constants and fitted to glia calcium activity using optimization. **b** Band-pass filtered (0.5 to 12 hours) calcium activity in EB and FB over time from ensheathing glia. Top row: ’walk’ state (velocity larger than zero in 1 second bins). Second and third row: activity in EB and FB. Red line: fitted homeostat model (two-state model); orange lines: fitted corrections for fluorescence levels (see Methods). **c** Same as b but for astrocytes in the EB and FB. **d** *L*2 error between fitted homeostat model and calcium activity of ensheathing glia and astrocytes in EB (left) and FB (right). **e** Distributions of time constants for ’stop’ and ’walk’ states from model fitting for ensheathing glia (*N* = 6 flies) and astrocytes (*N* = 5 flies) in EB (green) and FB (blue). One or two asterisks represent statistical significance with p-values lower than 0.05 or lower than 0.005, respectively, using t-test. (See Supplementary Information for models with 3 and 7 behavioral states, respectively.)

The resulting rise and decay time constants of ensheathing glia and astrocytes for 6 and 5 flies, respectively, in EB and FB are shown in Fig. 5d (see Supplementary Fig. S29c for ensheathing glia in the LAL, MB, and midline). The homeostat model fitted activity in the EB, FB, and LAL regions better than activity in the MB and midline, as shown by a lower fitting error (Supplementary Fig. S29d). Time constants in Fig. 2c were more similar between rise and decay than those obtained by model fitting (Fig. 5e and Supplementary Fig. S29c). This is due to the fact that active and rest states in Fig. 1j, k, and Fig.2b, were determined using low-pass filtering of walking activity (Fig. 1i and Fig. 2a), thus resulting in rest states that still contained a fraction of walking activity, and *vice versa* for active states. Time constants obtained from model fitting were not different between astrocytes and ensheathing glia (Fig. 5e).

Different behaviors contribute differently to sleep need and are therefore expected to charge the sleep homeostat with different time constants^1,14,15^. A model with 7 different behaviors, including proboscis extension and front or back grooming, improved model fitting (see Supplementary Information, ’7-state model’, Supplementary Figs. S31, S32, and Supplementary Tables S3, S4). Distinguishing two different sleep states depending on flies being immobile for less or more than 5 minutes (3-state model), resulted in similar quality of fits as the two-state model, with similar time constants independent of sleep state (see Supplementary Information and Supplementary Figs. S33, S34).

Thick line is low-pass filtered (0.1 hours cutoff period). **b** Same as a, but imaging in dFB neurons (labeled by R23E10). **c** Normalized individual (thin lines) and average (thick lines) fluorescence traces, as well as exponential fits (red lines, see Methods), during active and rest epochs for R5 neurons shown in a. **d** Same as c, but for dFB neurons shown in b. **e** Left side: Pearson correlation between the time flies are active and the average fluorescence traces for glia (in the EB, *N* = 6), and R5 neurons labeled by R88F06 (*N* = 8) and R58H05 (*N* = 5)). Right side: same as left side, but for glia (in the FB, *N* = 6) and dFB neurons (*N* = 8). **f** Pearson correlation between time resting and average fluorescence traces from glia and neurons, as in e. **g** *L*2 error between fitted homeostat model and calcium activity of glia and neurons in EB (left) and FB (right). **h** Approach followed to compute correlation between calcium activity and convolved ’walk’. First, walk is obtained from the behavior of the fly (first row). Then, ’walk’ is convolved with a triangular temporal filter with a defined size (second row) to obtain ’convolved walk’. Finally, the correlation between calcium activity and ’convolved walk’ is obtained. **i** Pearson correlation between ’convolved walk’ and calcium activity for different filter sizes (*x*-axis) for glia and neurons in EB (left) and FB (right). **j** Calcium activity in dFB neurons during sleep deprivation by perturbing the ball-supporting air stream after feeding. Normalized fluorescence traces of dFB activity (thin lines) and average (thick lines) before feeding, after feeding while the air valve was perturbing the ball, and after offset of the perturbation. **k** Normalized fluorescence levels (y-axis shared with panel j) before feeding, after feeding during sleep deprivation, and after offset of sleep deprivation. Sleep deprivation was performed by periodically interrupting air stream to ball. **l**, **m** Same as j and k, respectively, but with sleep deprivation by blocking the ball to induce effortful behaviors. Asterisks in all panels indicate statistical significance using t-test (*p <* 0.05).

### Comparison of glia and neuron dynamics

We next asked whether the observed dynamics in glia could result from activity of previously described homeostatic sleep circuits in the same brain compartments. In the EB, ring neurons (R5), increase calcium activity recorded in brain explants after extended sleep deprivation consistent with a sleep homeostat^33–35,74^, an effect which also depends on the circadian rhythm^75^. A second component of this circuit, dFB neurons, innervate the FB and are frequently used as a ’sleep switch’ to induce sleep^32,76,77^.

We recorded calcium activity of R5 neurons in the EB (with GAL4-lines R58H05 and R88F06, Supplementary Fig. S2d, e), and in dFB neurons (23E10-GAL4, Supplementary Fig. S2f), using multiple calcium indicators (GCaMP7f, GCaMP8f, and jGCaMP8m), as well as imaging protocols also with different time resolutions to potentially detect faster oscillatory dynamics^35^ (Fig. 6a, b, Supplementary Figs. S35, S38, S39, S43, S44, see Methods).

**Figure 6.**
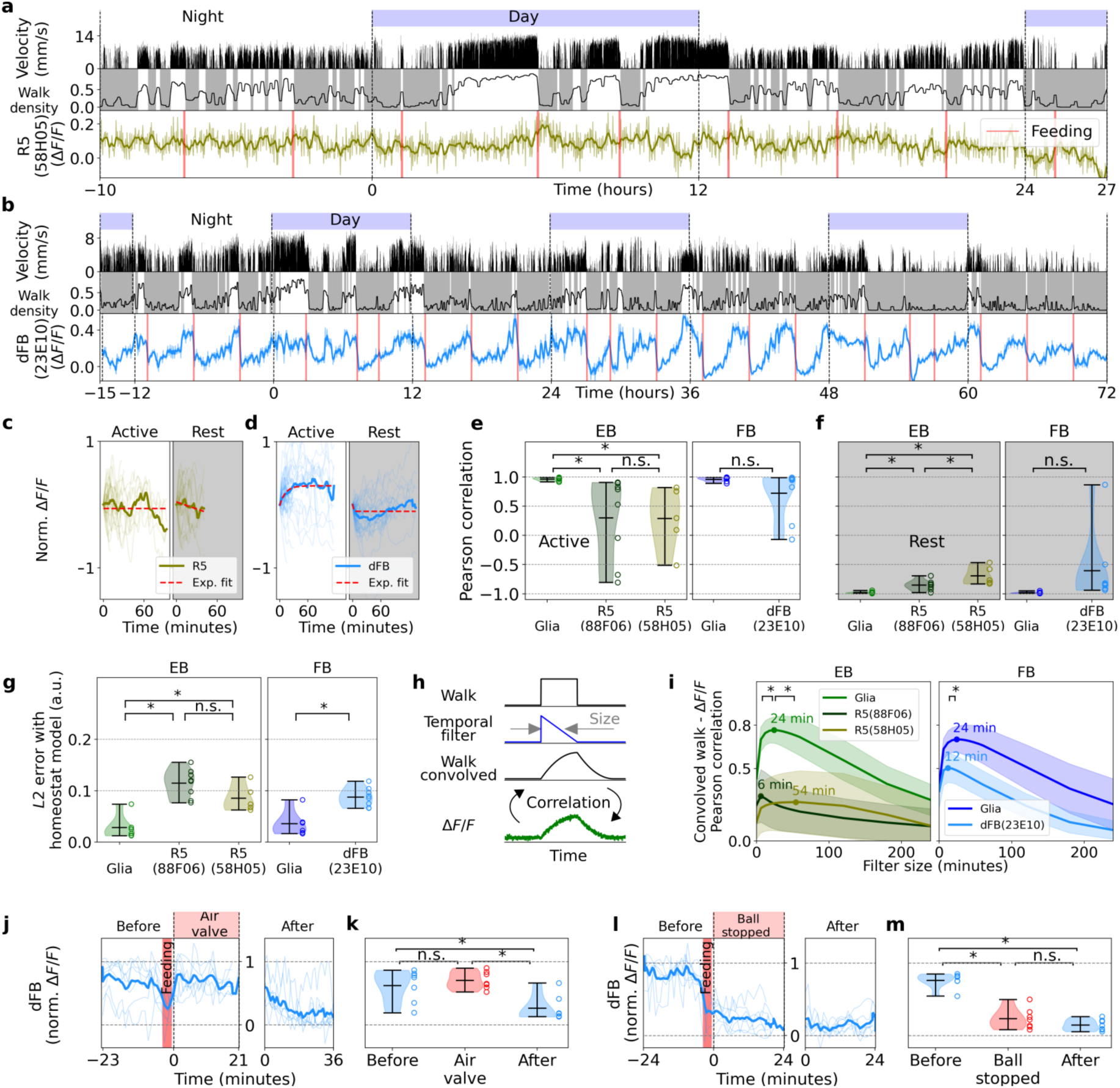
Calcium dynamics in sleep-related neurons and comparison with ensheathing glia. **a** Long-term imaging in R5 neurons (labeled by R58H05). Top row: day and night cycle in the VR. Second row: fly velocity. Third row: ’walk’ density, rest (grey region), and active epochs (white region). Fourth row: calcium activity in R5 neurons.

To determine whether these neurons encode sleep homeostasis similar to ensheathing glia in the respective neuropils, we first again determined active and rest epochs based on behavior (Fig. 6c, d, Supplementary Figs. S36, S40, S45). We then computed the correlation between active (Fig. 6e) or rest epochs (Fig. 6f) and the normalized calcium activity of neurons and glia (to avoid saturation we limited this analysis to traces of 30 minutes). For a sleep homeostat, the correlation should be positive during active epochs (increasing activity over time) and negative during rest epochs (decreasing activity over time). This was on average the case for both neurons and glia, with however considerably higher correlations for glia than for R5 neurons (Fig. 6e, f, left side). dFB neurons displayed high correlations during active epochs, while the slope of decreasing activity in dFB neurons was close to zero during rest epochs (Supplementary Fig. S53, see Methods).

We further analyzed whether a homeostat model as used for glia could fit activity of R5 and dFB neurons (Supplementary Fig. S37, S41, S42, S46, S47, and Methods). However, Fig. 6g shows that the L2 error between the fitted model and normalized activity was larger for both neural populations than for glia.

Finally, we computed the convolution between walking activity with a triangular-shaped temporal filter, which produces an exponential increase during ’walk’ bouts and a (symmetric) exponential decrease during ’stop’ bouts at a rate defined by the filter size (Fig. 6h, see Methods). We calculated the correlation between activity in neurons and glia with the corresponding signal resulting from the convolution of different temporal filter sizes with ’walk’ (Fig. 6i). The correlation with glia activity was high and peaked at a filter size of 24 minutes, similar to the dynamics obtained above. The correlation for R5 and dFB neurons was positive but lower compared to glia activity (Fig. 6i, see Methods).

Together, activity observed in ensheathing glia is not a simple reflection of activity of homeostatic circuits in the underlying compartments. Additionally, ensheathing glia better represent sleep homeostasis for the naturally occurring sleep and wake bouts than either R5 or dFB neuron activity recorded in the EB and FB, respectively.

### Hunger and walking state encoded in dFB neurons

While activity in dFB neurons was often correlated with walking and rapidly reset during feeding, resetting additionally depended on the behavioral state. If the fly was sleep deprived immediately after feeding by opening and closing the valve that controlled the air stream to the ball (mostly inducing quick bouts of fast walking), fluorescence activity increased again (Fig. 6j, k, and Supplementary Fig. S49). If the fly was sleep deprived by blocking the ball (which resulted in the fly pushing and pulling the ball but not coordinated walking), fluorescence reset (Fig. 6l, m, and Supplementary Fig. S50) indicating that dFB neurons track feeding and walking state.

Activity of dFB neurons additionally increased with time since the last feeding event, without a strong correlation with walking activity. We therefore averaged fluorescence traces of dFB neurons during hungry and fed epochs, defined as periods of 3.5 hours before and 0.5 hours after feeding events (separated by 4 hours), respectively (Fig. 7a, see below and Methods for a more detailed model). Activity of dFB neurons exponentially increased during hungry epochs, with time constants in the order of hours, and immediately decreased at feeding events with very fast time constants, in several cases limited by the time resolution of recordings of 1 minute (Fig. 7b, c, and Supplementary Fig. S48), indicating that dFB neurons could encode hunger, as has also been suggested for other FB neurons^78^.

**Figure 7.**
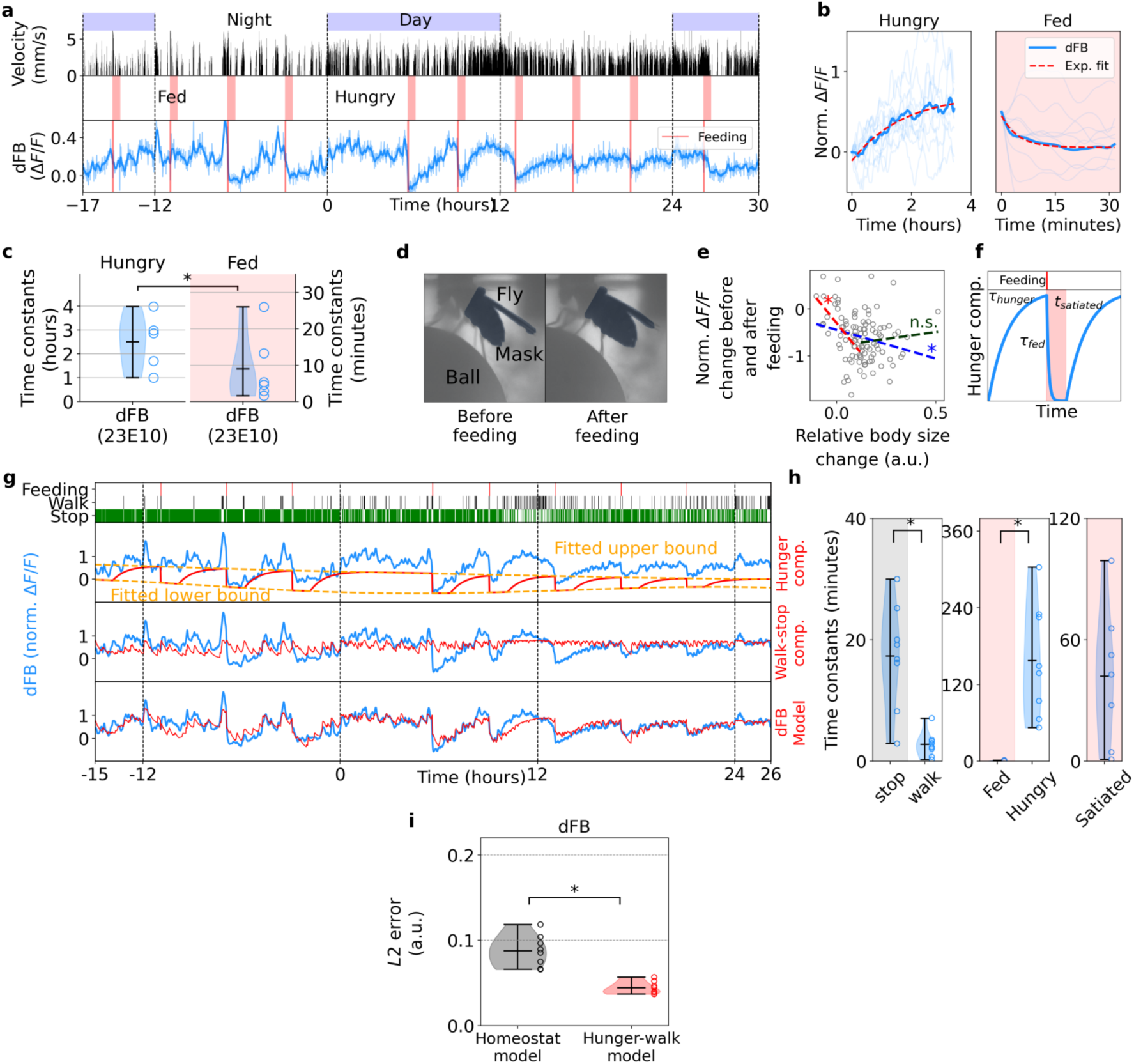
Feeding-related modulation in dFB neurons. **a** Long-term imaging in dFB neurons. Top row: day and night cycle in VR. Second row: fly velocity. Third row: ’hungry’ epochs before feeding (white region) and ’fed’ epochs after feeding (red region). Fourth row: calcium activity in dFB neurons. Thick line indicates low-pass filtering (with 0.1 hours cut-off period). **b** Normalized individual (thin lines) and average (thick lines) fluorescence traces and exponential fits (red lines, see Methods) during hungry and fed epochs from the recording in dFB neurons shown in a. **c** Time constants of fitted exponentials during hungry and fed epochs (*N* = 8 flies). Asterisk indicates statistical significance using t-test (p < 0.05). **d** Side view of a fly during a long-term imaging recording before feeding (hungry, left side) and after feeding (fed, right side). Mask of the abdomen (highlighted in dark color) for computing change in size. **e** Normalized fluorescence change against relative change in body size (see Methods) between 10 minutes before and 10 minutes after feeding. The blue line represents a linear fit over all points, while the red and green lines represent a linear fit over the first half and second half of relative body size change, respectively. Asterisks indicate the statistical significance of each fit using Pearson correlation (p < 0.05). **f** Parametrization (three parameters, *t_hunger_*, *t_f ed_* and *t_satiated_*) of the hunger component of the ’hunger-walk’ model. Before a feeding event (top row in red) the hunger component increases with the time constant *t_hunger_*and resets after feeding with the time constant *t_f ed_* for a given time *t_satiated_*. **g** Fitting of the hunger-walk model to activity of dFB neurons. Top row: events of feeding and walking and stopping bouts of the fly. Second, third and fourth row: normalized fluorescence (blue) and the fitted hunger component, ’walk-stop’ component, and the ’hunger-walk’ model (obtained from the two previous combined components), respectively (red). **h** Time constants (*N* = 8 flies) of stop and walk (left, from the stop-walk component), and time constants of ’fed’, ’hungry’, and ’satiated’ (center and right, from the hunger component). Asterisks indicate statistical significance using t-test (p < 0.05). *L*2 error between the fitted homeostat and ’hunger-walk’ models and the normalized activity of dFB neurons. Asterisk indicates statistical significance using paired t-test.

Using a dynamic model, activity of dFB neurons could be well described with a hunger component, which increases and decreases with respect to feeding events (Fig. 7f), and a walking component (Fig. 5a, see Methods), which were fitted together to the normalized fluorescence of dFB neurons ( ’hunger-walk’ model, Fig. 7g, Supplementary Figs. S51, and S52, see Methods for details). Fig. 7h shows the resulting model time constants, with slow calcium dynamics during the stop state (in the order of minutes), and feeding related time constants close to zero or the time resolution of our recordings, indicating that dFB activity resets immediately at feeding. Flies stayed satiated (with the hunger component not increasing) on average for around 40 minutes, although this varied between flies (Fig. 7h). The fitted ’hunger-walk’ model (Fig. 7g, Supplementary Figs. S51, and S52) produced a lower error with dFB activity (Fig. 7i) and statistically significant better fits (see Table S5 and Methods) than a homeostat model (Supplementary Figs. S46, S47). The very fast reset at feeding is not consistent with the behavior of a sleep homeostat.

Additionally, the decay amplitude during fed epochs was correlated with the amount of food ingested by the fly. The correlation between the relative change in abdomen size before feeding and after feeding events (as a proxy for the amount of food ingested, see Methods) with the change in fluorescence is shown in Fig. 7d and e. This correlation was stronger for small changes in body size (red line in Fig. 7e), suggesting that resetting of dFB neurons saturates after a certain increase in the volume of the abdomen (see Methods).

### Glia are linked to respiratory system

What mechanism could underlie the integration of behavioral activity over time in astrocytes and ensheathing glia? The structure of ensheathing glia, surrounding brain areas as a diffusion barriers, makes these cells well suited to detect the accumulation of metabolites inside these compartments. To further investigate this architecture, we took advantage of the fly connectome^37^ and noticed that glia surround trachea, which are responsible for gas exchange, as similarly observed in larvae^79^. By segmenting the tracheal system, glia matter, and EB and FB compartments (Fig. 8a), we found that trachea exclusively occupy the same space between brain compartments also covered by glia (Fig. 8b). Furthermore, tracheal tubes remain at the boundaries and do not enter neuropils (Fig. 8c). This suggests a close interaction between the respiratory system and ensheathing glia.

**Figure 8.**
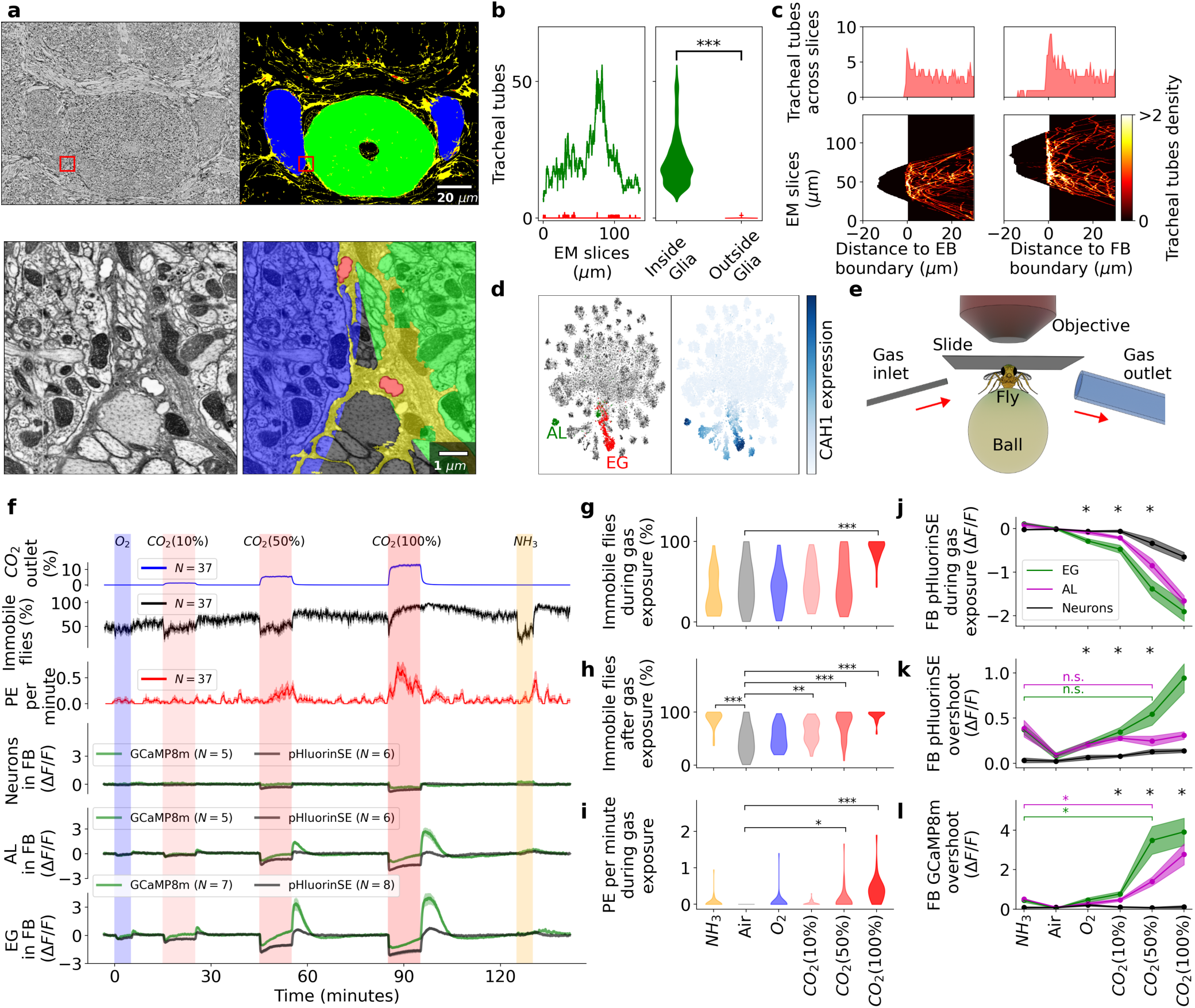
Tracheal system and CO_2_ / pH response in glia. **a** Left: slice from the EM hemibrain dataset (top) and zoom-in of red outline (bottom). Right: segmentation of EB (green) and FB (blue), glia (yellow) and tracheal tubes (red). Images are downloaded and adpapted from^37^. **b** Number of tracheal tubes in each slice (left) and distribution across all slices (right) of EM data detected inside (green) or outside (red) glia. **c** Distribution of tracheal tubes across slices (top) or for each slice (bottom) with respect to the boundary of the EB (left) and FB (right). Negative distance indicates tracheal tubes inside neuropils. **d** Right: scATAC sequencing of *Drosophila* brain projected in 2D using tSNE (data from^93^). Ensheathing glia and astrocytes are labeled red and green, respectively. Left: expression levels of CAH1. **e** Setup to deliver gases to the during imaging: inlet delivers gas to the fly walking on the ball during two-photon imaging, outlet removes gas. **f** First row: time of delivery of different gases. Air was delivered between the other gases at the same rate. Second row: measured CO_2_ levels at outlet. Third row: percentage of immobile flies. Fourth row: average of proboscis extensions per minute. Fifth, sixth and seventh rows: fluorescence levels of jGCaMP8m (green) and pHluorinSE (black) in neurons (N), ensheathing glia (EG), and astrocyte-like glia (AL), respectively. **g** Percentage of immobile flies during gas delivery **h** Percentage of immobile flies after gas delivery. **i** Proboscis extensions per minute during gas delivery. In g-i, one, two or three asterisks indicate different levels of statistical significance using a Kolmogorow-Smirnow test (p < 0.05, p < 0.005, or p < 0.0005, respectively). **j** pHluorinSE levels in the FB during gas delivery for N (black), EG (green) and AL (purple). **i** and **k** and **l** Overshoot of pHluorinSE and GCaMP8m levels in the FB after gas delivery, respectively. Asterisks at the top of j-l indicate statistical significance between N, EG, and AL for each gas using the one-way ANOVA test. See Supplementary Fig. S54 for the analysis in the EB.

We therefore hypothesized that the observed calcium dynamics could be linked to metabolic gasses such as CO_2_^80–82^. Due to higher metabolic demand of neurons^83^ more CO_2_ is produced during wakefulness than during sleep in flies^84^. In mammals, the glial support of metabolic activity of neurons includes the processing of gasses such as CO_2_^85–92^. Using a single-cell transcriptome dataset^93^, we further observed that carbonic anhydrase (CAH1), an enzyme which catalyzes the hydration of CO_2_, is highly expressed in astrocytes and ensheathing glia (Fig. 8d), further supporting the hypothesis that ensheathing glia and astrocytes are important for CO_2_ processing.

### Astrocytes and glia detect CO_2_ and pH changes

We therefore tested whether changes in CO_2_ concentration in the brain induces changes in glia and astrocyte calcium levels. Exposure of flies in the imaging setup to a CO_2_ plume (Fig. 8e) induced walking behavior at the two lower concentrations tested^94^ followed by sleep after offset of CO_2_ (Fig. 8f, top row, g, h), whereas the highest concentration induced immobility (Fig. 8f, top row, g, h). High CO_2_ concentrations also induced proboscis extension (Fig. 8i) . We imaged calcium activity in ensheathing glia (56F03-GAL4) and astrocytes (86E01-GAL4) as before, as well as in neurons by expressing sensors panneuronally using 57C10-GAL4^57,95^. To measure changes in pH we expressed the sensor pHluorinSE^96,97^ in these lines.

Exposure to CO_2_ (Fig. 8f, red sections) led to an initial rapid decrease in fluorescence of calcium as well as pH sensors expressed in neurons, astrocytes and ensheathing glia (Fig. 8j), as expected due to an induced change in pH^38–42^ (consistent with rapid diffusion of CO_2_ into tissue^98,99^). Interestingly, for astrocytes and ensheathing glia, but not for neurons, calcium levels slowly increased during CO_2_ application, and the CO_2_ offset was followed by an overshoot for both calcium as well as pH measurements (Fig. 8f). This suggests a slow homeostatic response counteracting pH changes in astrocytes and ensheathing glia, but not in neurons. Oxygen (O_2_, Fig. 8f, blue section) led to only a small decrease in calcium levels in EG. Ammonia (NH_3_)^100^ led to a long lasting increase in pH sensor signal expressed in ensheathing glia and astrocytes (Fig. 8j, k), consistent with alkalinization. Similar pH levels were observed both after NH_3_ exposure and after 50% CO_2_ exposure (Fig. 8k). However, calcium overshoot occurred only after CO_2_ exposure (Fig. 8l), indicating that calcium signals were not exclusively due to the impact of pH on the calcium sensor (see Methodsfor details).

### Optogenetic activation of astrocytes and glia induces sleep

The strong correlations between behavior and calcium activity, observed in awake and sleeping flies (Fig. 1, 2, 6), or following CO_2_ exposure (Fig. 8), raises the question of whether glia only track behavioral states, or also control them. We therefore expressed CSChrimson^43^, a cation and in particular proton channel^44^, together with jGCaMP8m in ensheathing glia or astrocytes^101^ for combined optogenetics and calcium imaging. A spatial light modulator (SLM) was used to activate CSChrimson in defined patterns using red light (Fig. 9a, see Methods and Supplementary Fig. S57) while at the same time imaging calcium activity in flies walking on an air supported ball as before. Optogenetic activation of a section of the EB (white wedge in Fig. 9a) induced sleep for both ensheathing glia and astrocytes, as seen by the percentage of immobile flies following activation (Fig. 9b-d). This shows that even local activation of astrocytes and ensheathing glia can cause sleep, as similarly observed using global thermogenetic manipulation of all ensheathing glia^49^ or astrocytes^50^.

**Figure 9.**
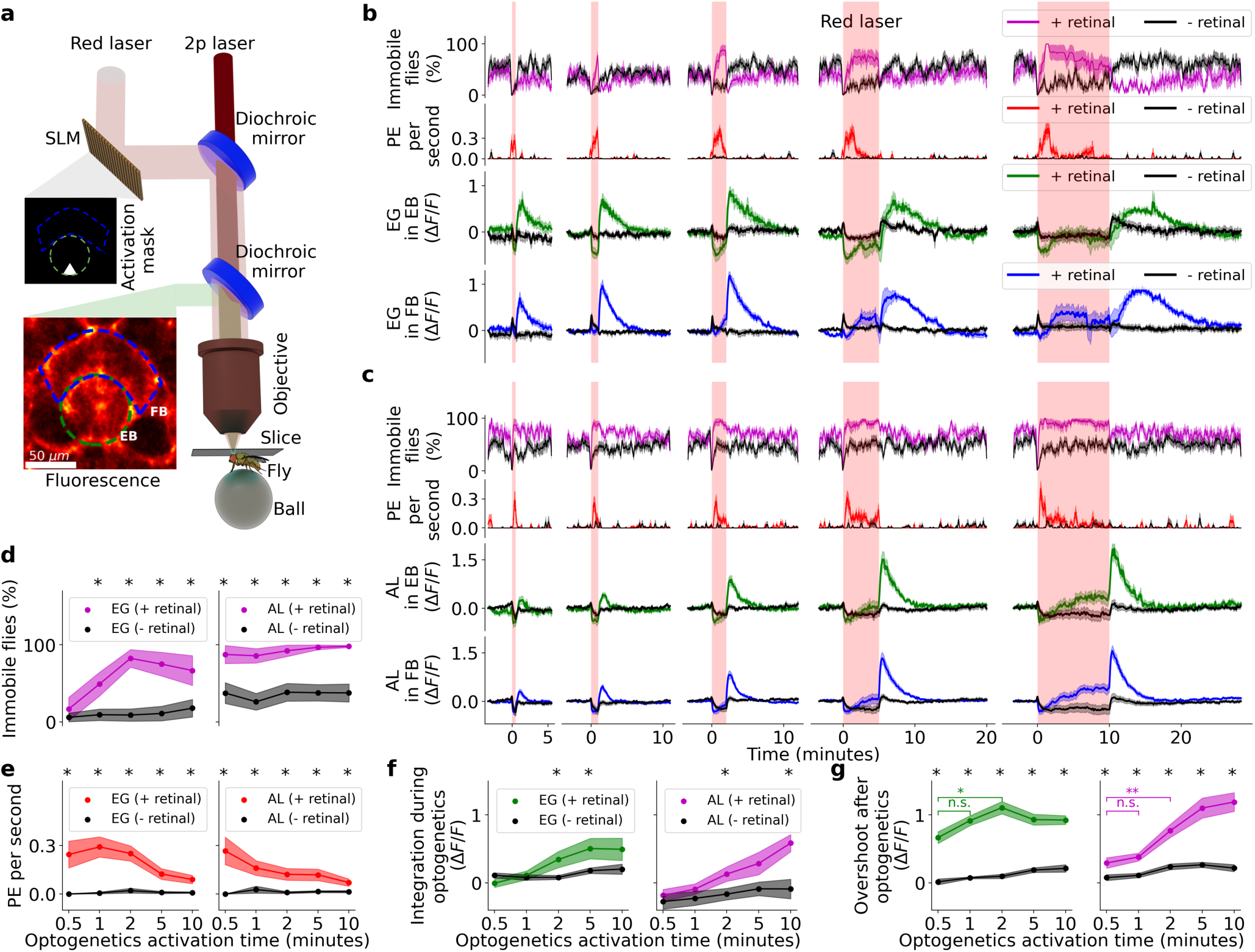
Combined optogenetics and imaging in ensheathing glia and astrocytes during behavior. **a** Optogenetics setup: custom light patterns for activation (shown as white triangle in EB) are generated by projecting a red laser into the microscope using a spatial light modulator (SLM). The activation and two-photon lasers are combined through a dichroic mirror and delivered to a fly walking on an air-supported ball. **b** Optogenetics in ensheathing glia: each column represents a trial with different durations of activation: 0.5, 1, 2, 5, or 10 minutes (red vertical areas). First row: percentage of immobile flies over time. Second row: average of proboscis extensions per second. Third and fourth row: recorded fluorescence from flies fed with retinal food (green for the EB activation site and blue in the FB, respectively) or with standard food as control (black). **c** Same as b but for astrocytes. **d** Percentage of immobile flies during optogenetics activation for each trial (x-axis) in ensheathing glia (left) and astrocytes (right). Purple: flies fed with retinal food. Black: flies fed with standard food. **e** Proboscis extensions per second during optogenetic activation for each trial (x-axis) in ensheathing glia (left) and astrocytes (right). Red: flies fed with retinal food. Black: flies fed with standard food. **f** Fluorescence drift or integration in FB, defined as the 0.9 quantile fluorescence value during optogenetics activation for each trial (x-axis) in ensheathing glia (left) and astrocytes (right). **g** Fluorescence overshoot in the FB, defined as the 0.9 quantile fluorescence value after optogenetics activation for each trial (x-axis) in ensheathing glia (green on the left) and astrocytes (purple on the right). Black lines represent control flies fed with standard food. Black asterisks at the top indicate statistical significance between flies fed with retinal food and control flies. Colored asterisks represent statistical significance across flies fed with retinal food (green for experiments in ensheathing glia, purple for experiments in astrocytes). All statistical significances were assessed using t test (one or two asterisks for p < 0.05 or p < 0.005, respectively). Black asterisks indicate statistical significance between + and *-* retinal flies, while colored asterisks compare + retinal flies in ensheathing glia (green) or astrocytes (magenta). See Supplementary Fig. S56 for EB analysis.

Fluorescence signal rapidly decreased at the onset of activation, consistent with an acidification of the activated compartment (EB)^44^, as similarly observed during CO_2_ exposure. As for CO_2_, this was followed by a slow drift of calcium levels during activation, both in glia and astrocytes (Fig. 9f), and offset of activation again led to an overshoot response (Fig. 9g). Control flies were not fed retinal food and immobility was significantly stronger in activated flies than in controls (Fig. 9d). Activation nevertheless led to small changes in calcium activity in controls, mostly during but not following optical stimulation, possibly due to residual or modified activity of CSChrimson. Interestingly, local optogenetic activation of EB also induced calcium signals in FB, suggesting that ensheathing glia interact across neuropils.

Optogenetic activation also induced proboscis extension (Fig. 9e), as observed before^101^, a behavior that occurs during sleep^20,62^, although at higher frequency than during natural sleep (Supplementary Fig. S56a). As for sleep bouts during long-term imaging, where proboscis extension occurred most frequently at the onset of immobility (Fig. 2i, k), proboscis extension was most frequent at the onset of optogenetic activation (Fig. 9b and c).

### Controller model of calcium dynamics

Both CO_2_ and optogenetics experiments revealed similar calcium and pH dynamics with drift during stimulation and overshoot after offset of stimulation (Fig. 8, 9), suggesting a pH-dependent mechanism of calcium dynamics. CO_2_ in solution is in equilibrium with bicarbonate and protons, and this equilibrium can be modulated in (mammalian) glia by carbonic anhydrases, pH buffers, and ion channels^38,40,41,102,103^. We use a model with two compartments to describe the observed dynamics, a neuropil compartment and a glia compartment (Fig. 10a, see Methods). We assume that pH in neurons is detected by glia or astrocytes and that a calcium dependent mechanism maintains pH at a physiological setpoint. Deviations from the setpoint in turn lead to changes in glia or astrocyte calcium levels.

**Figure 10.**
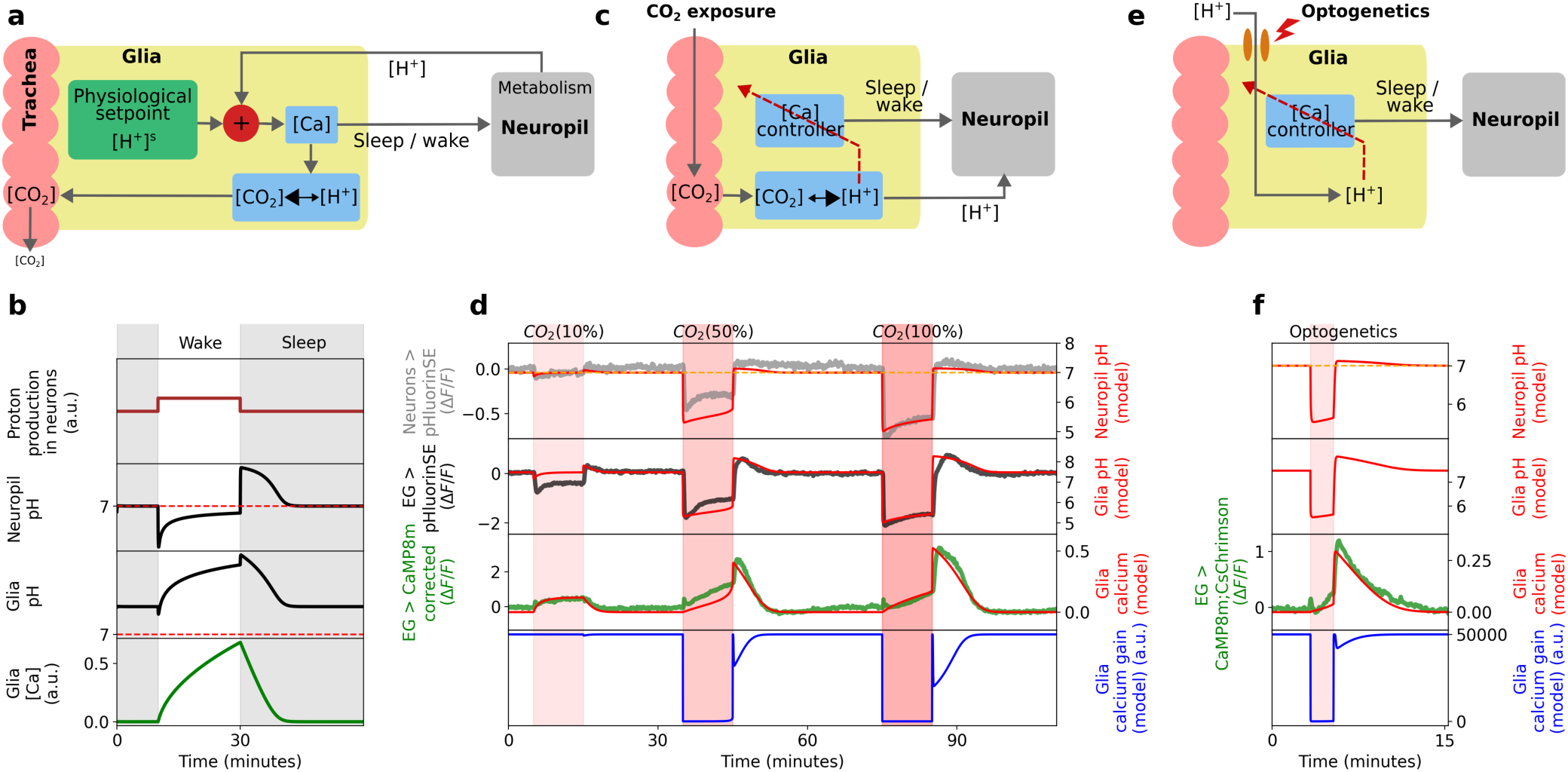
Controller model for glia calcium dynamics **a** Schematic of glia acting as pH controller: ensheathing glia (yellow) compare optimal physiological setpoint with pH changes (protons) in neuropil (grey) due to metabolic activity. Negative deviations of pH from setpoint increases calcium, which is assumed to shift the equilibrium between CO_2_ and protons to the left, increasing CO_2_ concentration for diffusive removal. **b** Model simulation during wake and sleep bouts. During wake (white area), more protons are produced in the neuropil (first row) compared to sleep (grey area). This leads to a decrease in pH in neurons (second row) and a pH increase in glia (third row). Calcium in glia increases (fourth row) to maintain pH in the neuropil back at setpoint (*pH* = 7). **C** Controller schematic during CO_2_ experiments. High concentration of CO_2_ enters the trachea, shifting the equilibrium between CO_2_ and protons to the right, lowering pH in glia as well as in neurons through diffusion. At very low pH, the calcium controller is partially inhibited (red dashed arrow). **d** Model dynamics during CO_2_ exposure. First row: application of 10%, 50%, and 100% CO_2_. Second and third row: average of fluorescence traces in neurons (measured with pHluorinSE, grey) and EG (black), as in Fig. 8f. In red, simulated dynamics oh pH inside the neuropil and glia. Fourth row: pH-corrected jGCaMP8m in EG (see Methods adn Supplementary Information), together with simulated calcium controller (red). Fifth row: simulated gain of the calcium controller, partially inhibited during CO_2_ application, produces slow dynamics and calcium overshoot afterwards (see Methods). Partial inhibition of the controller leads to a weak response, leading to slow integration. **e** Controller schematic during optogenetics. During optical stimulation, protons enter glia cells and lower pH. At very low the calcium controller is partially inhibited (red dashed arrow). **f** Model dynamics during optogenetics experiments. Second row and third row: simulated pH dynamics in neurons and glia. Fourth row: average of calcium traces recorded in EG during optogenetics experiments in FB (taken from Fig 9b, third column) compared to simulated calcium dynamics (red). Fifth row: simulated gain of the calcium controller partially inhibited during optogenetics activation.

Simulating the model (Fig. 10b) shows that neural metabolic activity during the wake state lowers pH in the neuronal compartment. This drop in pH is detected by glia, triggering an increase in calcium, which works to restore pH to the setpoint by shifting the equilibrium from protons to CO_2_ for removal through respiration (Fig. 10a, see Methods). This produces an increase of pH in the glial compartment. Glia alkalinization in the model accelerates proton diffusion from the neuronal to the glial compartments, bringing the neuropil pH back to its setpoint. When calcium activity reaches an upper threshold, it triggers sleep, which inhibits neural activity and prevents further acidification and alows resetting.

Drift and overshoot dynamics resulting from CO_2_ exposure or optogenetics can be modeled (Fig. 10d, f) by assuming that calcium regulation is partially inhibited when pH is outside normal physiological limits (Fig 10c and e), as is likely the case for the strong perturbations induced during CO_2_ exposure or optogenetics activation. When pH rapidly rises after CO_2_ exposure or optogenetic stimulation, the inhibition of the calcium controller is lifted (Fig. 10c, e, blue line on the last row), now detecting a sharp pH deviation from the setpoint. This results in a strong overshoot, similar to an impulse response in a control system (see Methods, see Supplementary Information 2.5 for correction of jGCaMP for pH, and 2.9 for possible biological mechanisms underlying model assumptions ).

## Discussion

Homeostatic control , that is sensing deviations from and resetting to a preferred physiological state, is key for the restorative and beneficial functions of sleep across species^1–5,7,8^. Many homeostatic processes, such as restoring energy levels or removing waste, are closely linked to the function of glia in the brain^89,90^. The dynamics of glia in the mammalian brain have therefore been a main target for the identification of sleep control^23–30^. In *Drosophila*, sleep homeostasis has been investigated extensively in neurons^14,15,34^, but the importance of glia has long been recognized^10,14^ and recent work points to multiple links between different classes of glia and sleep^49–52^. Here, by monitoring calcium dynamics in glia in flies navigating in virtual reality, we link sleep homeostasis to metabolism. We find that ensheathing glia and astrocytes encode all the information necessary for sensing sleep need. Both cell types show homeostatic calcium transients for the short sleep bouts typical for the fly. Both also show calcium fluctuations on circadian timescales (Figs. 1, 2), as also observed for gene expression^52,104^.

Ensheathing glia show all the characteristics expected of a sleep homeostat^45,66^: calcium levels increase with behavioral activity or effort during wakefulness, decay when the fly is sleeping, resting, extending its proboscis, or grooming (Fig. 1i-l, Fig. 3 and Supplementary Figs. S3, S6, S22, and Supplementary Videos S1-S6), and saturate during sleep deprivation (Fig. 4), all with an exponential time course. For example resetting of the homeostat to below 2% from baseline occurs over about 4 time constants or 60 to 120 minutes for the CX, or 80 to 200 minutes in the MB, a time span that covers a large fraction of the sleep bouts observed in freely walking flies^16,21,22^ (Figs. 1d and 5e). The time constants were faster for charging than for resetting of the homeostat (Fig. 5e and Supplementary Fig. S32a), consistent with the observation that flies sleep more than they are awake over a 24 hour period^21,22^. Reaching the upper or lower thresholds of the sleep homeostat did not automatically trigger sleep or awakening. However, long bouts of activity with a saturated homeostat were only observed under mechanically or hunger induced sleep deprivation (Fig. 4b and d).

Homeostatic sleep signals were observed across brain areas, such as the CX and MBs, but also in areas that have not been previously implicated in sleep, such as the LAL or the midline. That glia around different brain compartments show different calcium dynamics could be due to locally different activity and resulting varying sleep needs in different brain compartments, consistent with the idea of local sleep^47,50,73,105,106^. Local optogenetic activation of ensheathing glia as well as astrocytes in the central brain was sufficient to induce sleep (Fig. 9), and triggered proboscis extension, a behavior observed during sleep. Ensheathing glia and astrocytes can therefore both sense sleep need and control sleep.

While a frequently used definition for fly sleep relies on a threshold of 5 minutes, the arousal threshold increases already in the first 30 seconds to one minute after onset of immobility^18,62^. Additionally, most bouts of immobility are shorter than 5 minutes (Supplementary Fig. S30c), while short bouts of inactivity of maximally 20 seconds are sufficient for long-term survival of flies^67^, suggesting that already such short epochs of rest might share features of longer sleep bouts. Nevertheless, time from onset of immobility to onset of deeper sleep needs to be measured in the brain, as accomplished here by the dynamics of astrocytes and glia.

Both ensheathing glia and astrocytes, but not neurons, showed long-lasting calcium responses when the fly was exposed to a CO_2_ plume during tethered walking (Fig. 8). Similar calcium dynamics was observed when activating glia using optogenetics (Fig. 9). In both cases, continued stimulus application lead to a slow drift counteracting the induced changes, followed by an overshoot upon stimulus offset.

These dynamics can be described with a model which assumes that calcium levels in glia rise during active behavior, CO_2_ application, or optogenetic stimulation due to a concomitant decrease in pH (Fig. 10). While the strong perturbations due to high concentrations of CO_2_ or optogenetics are likely unphysiological, neural activity could lead to weaker acidification. Indeed, calcium dynamics similar to those observed during wake and sleep behavior are observed at low concentrations of CO_2_ (10% in Fig. 10d). In the model, sleep is triggered when calcium reaches an upper threshold, which could for example prevent excessive energy expenditure on the glia controller^103^, allow to replenish energy storages, or to restore pH bicarbonate buffers and substrates that are coupled with proton shuttles such as lactate or glutamine^107^. Dysregulation of astrocytic pH is linked to various diseases of the brain^103^.

That glia do not only show slow integration (Figs. 1, 2, 3a-d), but also calcium peaks at activity onset after extended bouts of immobility (Fig. 3g), suggests that a more detailed controller model should also include a differential component. Such a differential control element, which anticipates future changes, would also be consistent with the anticipatory metabolic activity observed in neurons^83^.

Proboscis extension has been linked to waste clearance during sleep^20^, potentially serving to remove CO_2_ from the brain^108^, and in turn reducing elevated proton concentrations. That proboscis extension events are more frequent at the onset of sleep, when acidification of neurons is likely higher due to continued neural activity during prior wakefulness^41,42,109,110^, would also agree with such a mechanism (Fig. 2i). Moreover, proboscis extension is also triggered by optogenetic activation of ensheathing glia and astrocytes (Fig. 9e), and by high concentrations of CO_2_ (Fig. 8i), consistent with a role in maintaining pH balance in the brain.

The arousal threshold increases with sleep duration (Fig. 2h), at least initially^18^, while calcium levels in glia reset over time. At the end of wake bouts, increased neural excitability, and corresponding lower arousal threshold^111,112^, could result from increased connectivity strength, as expected based on the synaptic homeostasis hypothesis^113^, higher excitability due to pH changes^114^, or due to accumulation of gliotransmitters. This agrees with increased EEG slow-wave amplitudes in mammals^47,115^ and flies (specifically in R5^33,35^ and dFB neurons^116^) after sleep deprivation. Thus the homeostat is expected to be at a maximum at the end of wake or beginning of sleep bouts as observed.

Of the different types of neurons that have been linked to sleep behavior^14,15^, we here studied a subset of neurons in the center of the fly brain^33–35^. However, glia activity describes wake and sleep bouts more reliably than the investigated neurons inside the corresponding brain compartments (Fig. 6e-i). Ring neuron activity in the EB^33,35^ was not well described with a homeostatic model. Interestingly, activity of dFB neurons in the FB^32^ could be described by taking into account walking activity and hunger^78,117,118^, but not with a sleep homeostat (Fig. 7g-i and Supplementary Figs. S46, S47, S51 and S52). The often very rapid reset of activity after feeding in dFB neurons, which is not consistent with the behavior of a sleep homeostat, depends on the behavioral state of the fly, which is not the case for ensheathing glia. Glia activity remained high during sleep deprivation after feeding (Figs. 3e, f, 4a, Supplementary Figs. S21a, and S17), as expected of a sleep homeostat, whereas in dFB neurons only walking activity, and not other effortful behaviors, could suppress resetting of calcium levels (Fig. 6j-m, Supplementary Figs. S50, and S49). These findings are also consistent with recent observations that dFB neurons might not have the previously assumed role for sleep control^119^, but would be consistent with a role of the FB in feeding behavior^78,120^. The mechanism by which glia act on neurons likely involves the calcium dependent release of gliotransmitters^121^.

Modulation of sleep circuits by glia has for example been described in C. elegans^122,123^. In mice, microglia, which share some of the functionalities of ensheathing glia in flies, generate inhibitory feedback and suppress activity of glutamatergic neurons^124^, a mechanism which could also play a role in sleep regulation in an adenosine-dependent manner^125^. The action of ensheathing glia could also include the regulation of glutamate in the enclosed circuits^12,65,72^. We observed an increase or respectively a decrease of extracellular glutamate at ensheathing glia in the CX during active and inactive, or wake and sleep states (see Supplementary Information and Supplementary Figs. S26a-c,S27, and S28). Ensheathing glia release adenosine with increasing calcium levels^126^, and adenosine signaling is more broadly important for metabolic control in other *Drosophila* glia as well, including astrocytes^101^. Additionally, increased extracelullar adenosine is also modulated by decreasing pH^127^. Another gliotransmitter in ensheathing glia could be taurine which promotes sleep in flies^49^. Further, astrocytes release D-serine^128^ which is also important for sleep in flies^129^.

Together, ensheathing glia and astrocytes offer homeostatic controllers for locally sensing and controlling sleep distributed across the entire brain for the short, naturally occurring sleep bouts typical of the fly. In the future, it will be of great interest to use local manipulation and control of sleep in different behavioral paradigms, including learning, to gain an better understanding of the interplay between behavior and sleep function, in particular its homeostatic and metabolic aspects.

## Data availability

All of the behavior data, raw functional imaging data, and associated behavior data can be made available upon request.

## Code availability

All of the code for data processing, analysis and figure generation can be made available upon request.

## Acknowledgements

We thank Anna Kaatz for fly crosses and dissections for imaging experiments. We thank Marcelo Cintra, Christoph Geisen, Omar Valerio Minero, and Sercan Yegin for help with the MPINB cluster, Jasmin Beinert, Alexandra Neu, Tim Krause, Tolga Ates and Anne Buecker for maintaining flies, and Vivek Jayaraman for Chrimson flies. We thank Jason Kerr for the Mai Tai Laser as well as other equipment, and Alexandr Klioutchnikov for helpful discussion. We thank Kevin Briggman and Jason Kerr for comments on the manuscript.

## Author contributions

AFV and JDS designed the study, built the experimental setups, performed experiments, and wrote the manuscript. AFV performed data analysis. IV developed software for optogenetics.

## 1 Methods

### *Drosophila* preparation

All flies were reared in an incubator at 25 degrees with a 12 hour light/dark cycle. For experiments with freely moving flies, we used 7 days old females expressing jGCaMP8m in ensheathing glia (UAS-jGCaMP8m;56F03-GAL4). Flies were anesthetized on ice and placed individually in rectangular chambers with food (Fig. 1a).

The same 12 hour light/dark cycle as in the incubator was used in VR during imaging experiments. All imaging experiments were performed with female flies between 3 and 8 days old at the beginning of the experiment. To record calcium activity of ensheathing glia, astrocytes and in neurons, we expressed jGCaMP8m using the GAL4 lines 56F03 (UAS-jGCaMP8m;56F03-GAL4), 86E01 (UAS-jGCaMP8m;86E01-GAL4), and 57C10 (UAS-jGCaMP8m;57C10-GAL4), respectively.. Experiments for monitoring extracellular glutamate were performed by expressing the glutamate sensor iGluSnFR in ensheathing glia (UAS-iGluSnFR;56F03-GAL4). To monitor activity of R5 ring neurons we used two different GAL4 lines: 58H05-GAL4 expressing jGCaMP8m (UAS-jGCaMP8m;58H05-GAL4), and 88F06-GAL4 expressing jGCaMP8m, 8f or 7f (UAS-jGCaMP8m;88F06-GAL4, UAS-jGCaMP8f;88F06-GAL4 and UAS-jGCaMP7f;88F06-GAL4), respectively. Activity of dFB neurons was monitored by expressing jGCaMP8m or jGCaMP8f in 23E10-GAL4 (UAS-jGCaMP8m;23E10-GAL4, UAS-jGCaMP8f;23E10-GAL4). To record pH in ensheathing glia, astrocytes and neurons during gas exposure experiments, we expressed pHluorinSE (Bloomington stock number 82176) in the respective cell types (UAS-pHluorinSE;56F03-GAL4, UAS-pHluorinSE;86E01-GAL4 and UAS-pHluorinSE;57C10-GAL4). For optogenetics experiments we expressed both jGCaMP8m and CSChrimson-mCherry (Bloominton stock number: 82180) in ensheathing glia (UAS-CSChrimson, UAS-jGcAMP8m;56F03-GAL4) and astrocytes (UAS-CSChrimson, UAS-jGcAMP8m;86E10-GAL4).

For imaging, flies were dissected using laser surgery as described^55^ to insert a transparent window into the cuticle^55,130^. The cut cuticle and air sacks were removed under a dissection microscope using forceps, either manually or with a microrobotic arm^55^. The opening was sealed with a drop of transparent UV glue (Detax 02829, freeform fixgel 3g syringe, purchased before 2023^55^, or Norland Optical Adhesive NOA 68^130^). Between 2 and 5 flies were dissected at a time and were left to recover in vials with food for 1 to 3 days before imaging. Flies were then glued to a glass slide using UV glue and transferred to the long-term imaging setup or to a second microscope modified for gas experiments. Flies were selected for imaging based on the optical access to the structures of interest which varied dependent on dissections. Only data recorded at least 48 hours after surgery were included in the analysis.

For optogenetics experiments, flies expressing CSChrimson and jGcAMP8m were initially reared on regular food, with dissections carried out as previously described. After dissection, control flies were transferred to a vial containing regular food, while non-control flies were placed in a vial with retinal food. Both control and non-control flies were left to recover in darkness for two days before imaging experiments. Retinal food was prepared by adding 50 *µ*L of a solution containing 100 mg of all-trans-retinal dissolved in 3.52 mL of 96% ethanol to vials containing regular food.

### Setup for experiments with freely walking flies

Flies were placed in transparent chambers of 70 ⇥ 4 mm (Fig. 1a) and activity was recorded in 15 chambers using a camera from above with a resolution of 1920 ⇥ 1080 pixels at 30 Hz. A rectangular grid of IR LEDs illuminated the chambers from below through a diffuser. A longpass filter (Thorlabs FGL780S) in front of the camera rejected white light from LEDs which generated the same 12 hour light /dark cycle as in the incubator. Since flies needed to adapt to the new environment, the first day of the experiment was discarded, and we only considered the subsequent two days for the following analysis.

Tracking of fly positions was done offline, using color segmentation in OpenCV and Python, by extracting the dark color of the flies from the bright background (Supplementary Video S8). The position and velocity of each fly were smoothed using a Kalman filter. Fig. 1b shows examples of the position along the chamber and the velocity of one fly. Supplementary Figs. S1a and b show the mean velocity of 15 flies with a light/dark cycle and in darkness over 48 hours, respectively. Under both conditions flies displayed circadian activity.

To find epochs of immobility, we set the velocity of each fly to zero if a fly did not move for at least 0.25 body lengths in a second (a body length was defined as 2.5 mm) (Fig. 1c, first row). We then computed each fly’s ’stop’ binary profile over time from the velocity (Fig. 1c, second row). To compute sleep bouts, we used different temporal filters with a size of 1, 2, 5, 8, and 10 minutes (Fig. 1c, third row) and convolved them with the binary ’stop’ signal (Fig. 1c, third row). These temporal filters removed small bouts of movement of the fly (depending on the filter size) between stop epochs. Sleep was then obtained by thresholding the convolved ’walk’ signal at the value of 0.5 (Fig. 1c, fourth row). The distribution of continuous sleep bouts was computed for each filter size during the day and night (Fig. 1d), and during darkness (Supplementary Fig. S1c), and the 90% quantile of the distribution is highlighted. A filter size of 2 minutes, which already removes up to 1 minute of walking in between stop epochs, produces a distribution where 90% of sleep bouts are below 50 minutes (Fig. 1d). Sleep bouts were even shorter in dark conditions, where even using a filter size of 10 minutes, which removes 5 minutes of walking in between stop bouts, led to a distribution where 90% of sleep bouts were below 40 minutes (Supplementary Fig. S1c).

To compute how frequently flies ate in these behavior experiments, we determined all occasions where the fly was close to the food (at around 7 *mm* in the chamber, see Fig. 1b). As before, we used temporal filters with different sizes (30, 60, 90, 120, and 150 seconds) to filter out small movements of the fly entering and leaving the food. We then obtained the distribution of time intervals between consecutive feeding events during the day and night depending on filter size (Supplementary Fig. S1d). Using a filter size of 120 seconds, 90 % of the times between consecutive feeding events were around 26 minutes during both day and night. Similar rates of feeding were used in some of the imaging experiments, where flies were fed every 26 minutes (Fig. 1g and Supplementary Fig. S9) or every 16 minutes (Supplementary Fig. S13).

### Imaging setup

The setup for long-term imaging was as described in^55^. For recording volumetric calcium activity and to reduce brain motion artifacts, two axially offset focal planes were recorded at the same time with beams with an extended focal length, with temporally multiplexed acquisition^55^. This setup allowed motion correction at high time resolution in all three dimensions as described in^58^. Virtual reality projection and closed loop behavior were implemented as previously described^55,131^.

### Data acquisition

At the start of each long-term imaging experiment, a z-stack of 100 *µ*m depth with an axial step size of 0.25 *µ*m was recorded, which was used for z-motion correction in post-processing^58^ (see next section). Additionally, an automated robotic feeder was configured at the beginning of the experiment, for example to feed flies every 4 hours for 2 minutes^55^. Imaging data was recorded at a resolution of 256 ⇥ 256 pixels at 60 Hz. In most of the experiments, we recorded for 1 second with a wait time of 60 seconds between consecutive recordings, resulting in a total imaging time of 1.6% of the duration of the experiment, which helped to reduce photobleaching and phototoxicity. However, we additionally used a different protocol in four other flies, where we recorded in trials of 30 seconds with a wait time of 5 minutes between trials, which resulted in a total imaging time of 11% of the duration of the experiment (Supplementary Figs. S44g and S39f-h).

The VR, implemented as described^74,131^ displayed a dark stripe on a bright background using a blue laser (488 nm), which was switched off during the night for 12 hours. The VR night and day cycle was the same as the one used in the incubator. Ball movement was tracked at 200 Hz and a calibration factor was determined for calculating ball velocities. The water temperature at the objective was maintained at 22 degrees with a perfusion system^55^.

Flies were left to recover for at least 2 days after surgery and before imaging started. In some flies, imaging started after 1 day of recovery and the first day was excluded from data analysis. Recordings of calcium activity in ensheathing glia are shown in Figs. 1i and Supplementary Figs. S3, S16, S9 and S13 for EB and FB, Supplementary Fig. S4 for LAL, and Supplementary Figs. S5 and S19 for MB and midline. Recordings for astrocytes are shown in Fig. 2a and Supplementary Fig. S11. Recordings of glutamate activity in ensheathing glia are shown in Fig. S26a and Supplementary Fig. S27. Recordings of calcium activity in R5 neurons using 58H05-GAL4 are shown in Fig. 6a and Supplementary Fig. S35, and recordings using 88F06-GAL4 are shown in Supplementary Figs. S38 and S39. Finally, recordings of calcium activity in dFB neurons are shown in Fig. 6b and Supplementary Figs. S43, S43, S49 and S50.

### Data post-processing

Imaging data was corrected for lateral motion by alignment with respect to a template (average of the first 60 frames) using cross-correlation (Fig. 1g). The z-stack was also aligned with respect to this template. Then, 24 regions of interest (ROIs) were defined for recordings in ensheathing glia in different brain areas (the 24 ROIs merged into a single one are shown in Fig. 1g for EB and FB, Supplementary Fig. S2b for LAL and Supplementary Fig. S2c for MB and midline). ROIs were also defined for the case of R5 and dFB neurons (the combined 24 ROIs are shown in Supplementary Fig. S2d,e for R5 neurons, and f for dFB neurons). The intensity of each ROI was computed by summing all pixel values within each ROI per frame as well as in each frame in the z-stack. The intensity of the ROIs was then used to estimate the z-motion of the sample for each imaging frame as described^58^. The resulting z-motion over the time of the experiment was filtered using a median filter with 1000 points to discard high-frequency motion, and maximum displacements of around 10*um* along the z-axis were detected. The filtered z-motion was used to estimate the fluorescence of each ROI corrected for z-motion^58^. Differences between corrected and uncorrected fluorescence was generally low, with maximum differences of 0.1 D*F/F* due to the extended focal volume used for imaging^58^.

The 200 Hz used to track ball motion underestimated ball displacements in the 3 axes by a factor of 0.50 compared to using 500 Hz tracking as in^131^. This factor was experimentally measured by rotating the ball by 3600 degrees. Ball displacements during long-term imaging were corrected with this calibration factor in post-processing and used to compute the absolute velocity of the fly in bins of 1 second using the ball radius (3*mm*).

For fitting models to fluorescence traces (see sections below), we first smoothed the fluorescence signal using a low-pass filter with a cut-off period of 30 minutes. Then, we normalized the filtered fluorescence, *F*(*t*), by remapping the 10% and 90% quantiles of *F*(*t*) (*Q*_10%_ and *Q*_90%_, respectively) to the values 0 and 1, respectively, with the following equation:

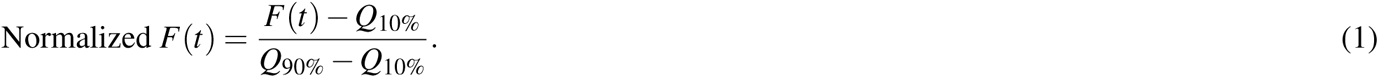

This was done to reliably compare glia and neurons in Fig. 6e-i, since different recordings have different fluorescence dynamic ranges and different noise levels. By smoothing and normalizing fluorescence traces from all experiments, we ensured that correlations (Fig. 6e, f and i) and errors from model fittings (Fig. 6g and Fig. 7i) were correctly compared across recordings.

### Behavior classification

Fly behavior was monitored during the entire experiment using a camera (Supplementary Fig. S14a) under IR illumination at 10 frames per second (fps). Based on blocks of 10 consecutive frames (1 second time resolution), we classified the following 7 behaviors (rows in Supplementary Fig. S14A). Stop: the fly does not move. Proboscis: the fly does not move but extends and retracts the proboscis. Walk: the fly walks on the ball. Discomfort: the fly pushes or pulls the ball. Grooming front: grooming of head and proboscis with front legs. Grooming back: grooming of the abdomen and wings with hind legs. Feeding: fly is fed with the feeding robot.

We used a 3D convolutional neural network (3D CNN) to classify behavior based on 10 frames at a time. The field of view of the camera (with resolution 640 ⇥ 480 pixels) was cropped around the fly and was resized to a resolution of 256 ⇥ 256 pixels. The architecture of the 3D CNN is shown in Supplementary Fig. S14b, where each convolutional block consisted of a 3D convolutional layer with a 3 ⇥ 3 ⇥ 3 kernel size and rectified linear unit activation function (ReLU), a dropout layer with a rate of 0.2 to prevent overfitting, and 3D max pooling with size 2 ⇥ 2 ⇥ 1. The number of filters of the convolutional layer increased by a factor of 2 in each consecutive convolutional block. Finally, a dense layer with a softmax activation function assigned a value to each class. The 3D CNN had a total of around 6.5M trainable parameters.

A total of 45610 manually labeled frames from 10 different flies were used for training. The dataset was normalized to train the network with the same number of samples for each class. We further increased this dataset with data augmentation that randomly transformed the input frames by changing illumination, translation, rotation, and/or scale during training. To train the network we used TensorFlow with a binary cross-entropy loss function and Adam as optimizer. After 150 epochs with a batch size of 50 samples, the loss function reached a minimum saturation value in about 2 hours, using 4 QUADRO GPUs.

To test the performance of the network, we used 2410 labeled frames, which were augmented to a total of 20000 with the previously described augmentation transformations. The accuracy of the network to predict the right class of behavior was 98.6%, and the normalized confusion matrix for each class is shown in Supplementary Fig. S14c. An example of behavior classification is shown in Supplementary Video S7.

### Probing of arousal threshold

For probing the arousal threshold of the fly during long-term imaging, we used a total of 6 flies where we recorded calcium activity (data not included) from ensheathing glia using the same protocol as before. We recorded multiple trials of probing the arousal threshold in each fly. Trials were initiated automatically using custom software (see number of trials, *N*, for each fly in Supplementary Fig. S15b-g). At the beginning of each trial, the amount of expected sleep time for probing the arousal threshold was selected randomly, either 30 seconds or 5 minutes. Then the fly was stimulated to walk by closing and opening the valve controlling the air stream supporting the treadmill ball repeatedly 3 times during 6 seconds (1 second closed and 1 second open). This ensured that the fly was awake at the beginning of each trial. After this, the velocity of the fly on the ball (thresholded to remove tracking noise) was monitored in real-time to detect epochs of walking or immobility. A timer was used to measure the time the fly was not walking since the end of the last walking bout (non-zero velocity). If this time reached the previously selected duration (30 seconds or 5 minutes), probing of the arousal threshold started. Probing was performed with an IR laser focused on the abdomen (Toptica, ibeam-smart-785-S-HP with pulse option, 785 nm) by ramping the power from 0 to 100*mW* in steps 1*mW* every second. When the fly started to walk due to the heating effect of the IR laser, the trial finished and the IR laser was turned off. During probing with the IR laser, the shutter of the two-photon laser was closed and imaging data was not recorded, since the combined power of the two-photon and heating lasers would distort the detection of the threshold. After a trial had finished, the next trial started after 10 minutes of waiting. Experiments for each fly were performed for at least 1 day, resulting in several hundreds of probing trials in each fly. We used the 3D CNN to classify the behavior in post-processing. For each trial, we computed the actual elapsed time that the fly was immobile (either in the ’stop’ state or ’proboscis’ state) prior to the arousal threshold probing, in order to remove grooming events. Only trials with prior times of complete immobility larger than 5 seconds were considered, therefore the resulting times of immobility ranged from 5 seconds to 5 minutes. During the probing period in each trial, we obtained the required laser power to awake the fly. The fly was considered awake when it showed walking, discomfort, or grooming behaviors (obtained from the 3D CNN classification). An example of a trial for arousal threshold probing is shown in Supplementary Fig. S15a.

We computed the distribution of IR laser powers required to awake the fly prior to three different time intervals of immobility: [5, 35], [35, 200], and [200, 305] seconds. These distributions are shown in Supplementary Fig. S15b-g for individual flies, and in Fig. 2h for all flies. Asterisks indicate statistical significance using t-test (p-value < 0.05). As shown for individual flies as well as for all flies (Fig. 2h and Supplementary Fig. S15b-g), longer times of immobility required higher IR laser powers to awake the flies, confirming that the arousal threshold increases with time of immobility during our long-term recordings.

### Proboscis extension during sleep

We used the trained convolutional neural network to classify the behavior of 21 flies during long-term imaging sessions, which included recordings from ensheathing glia, dFB, and R5 neurons. Based on this classification, we identified periods of immobility, defined as the fly either being in the ’stop’ state or engaged in proboscis extension (’proboscis’ state). We excluded bouts of immobility lasting less than one minute and computed the distribution of proboscis extension events in 30-second intervals during sleep. Fig. 2i shows the average distribution of proboscis extension events across all sleep bouts for the 21 flies. For each time bin, we required at least 10 sleep bouts to be included in the average. Since we did not find more than 10 sleep bouts with more than 20 minutes of uninterrupted immobility, the distribution of proboscis extension events in Fig. 2i is limited to the first 20 minutes of sleep. Proboscis extension data beyond 20 minutes are therefore not schown.

We also computed the inter-PE interval, defined as the time between two consecutive proboscis extension events (Fig. 2j), which shows a peak at around 3 seconds, at previously described^20^.

### Fitting of fluorescence traces during ’active’ and ’rest’ states

Sleep is commonly identified in flies as bouts of immobility that last for at least 5 minutes^17,132^, a threshold that has also been used for assessing sleep in tethered walking flies^133,134^. However, shorter bouts of quiescence already show many of the characteristics of sleep^19^. The distribution of sleep bouts, defined as bouts of at least 1 minute of immobility during long term recordings in ensheathing glia, astrocytes, R5 neurons, and dFB neurons is shown in Supplementary Fig. S30a and b for the day and night, respectively, including a total of 21 flies. We compared these distributions to sleep bout distributions in freely moving flies using the filter size of 0 minutes to define sleep (Fig. 1c and d). During the day, the distribution of sleep bouts in flies during imaging compared to freely moving flies are highly similar (statistically different as asses with a Kolmogorow-Smirnow test (p-value < 0.05, Supplementary Fig. S30a). During the night, they are not significantly different (Supplementary Fig. S30b). We also confirmed that around 80% to 90% of sleep bouts in flies during imaging and freely moving flies are shorter than 5 minutes (Supplementary Fig. S30c).

We compare the amount of sleep according to a 5 minutes threshold, only distinguishing between walking and stopping, and according to the ’rest’ and ’active’ states (used in Fig. 1i) in Table S1.

To find epochs of at least 10 minutes during which the fly was walking most of the time (active) or sleeping most of the time (rest), first, walking velocity on the ball (with a diameter of 3 mm) was averaged over one second.

Then, velocities below a threshold of 0.25 body lengths per second (body length defined again as 2.5 *mm*) were set to zero to remove tracking noise, and we defined a binary walk state of the fly, *w*(*t*), when the thresholded velocity was non-zero (*w*(*t*)= 1). We defined active and rest states where the fly was walking or standing still most of the time. To calculate these states, we used a walk density, which was obtained by low-pass filtering of the walk state with a period of 0.1 hours (6 minutes). Active and rest states were defined based on the walk density being above or below a threshold of 0.5, respectively (Fig. 1i and 2a and Supplementary Figs. S3, S4, S5, S11, S27, S35, S38, S39, S43, S44). We subdivided the normalized fluorescence of each recording (equation (1)) into *N_a_* traces of at leas *active* 10 minutes of continuous active epochs, 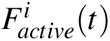 for *i* = 1*, …N_a_*. At least 4 trace traces were used from each fly to obtain the normalized averaged fluorescence over active epochs, which was fitted with an exponential (Figs. 1j, 2b, and 6c, and left side in Supplementary Figs. S6a-e, S7a-e, S8a-e, S12a-d, S28a-d, S36a-d, S40a-h and S45a-g). The exponential was defined by the following equation:

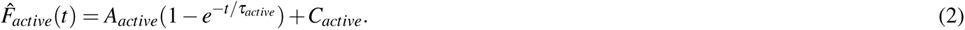

Here, *A_active_*is the saturation level, *C_active_*is an offset, and *t_active_*is the time constant.

Fluorescence traces for the rest state were similarly selected as epochs of at least 10 minutes. At least 4 normalized fluorescence traces were used to obtain the normalized average fluorescence over rest epochs, which was fitted with an exponential (Figs. 1k, 2b, and 6d, and right side in Supplementary Figs. S6a-e, S7a-e, S8a-e, S12a-d, S28a-d, S36a-d, S40a-h and S45a-g). The exponential was defined by the following equation:

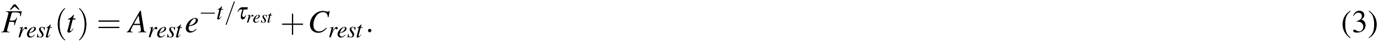

Here, *A_rest_* is the amplitude, *C_rest_* is an offset and *t_rest_* is the time constant of the decay. The time constants during active and rest states for activity in ensheathing glia, astrocytes, and extracellular glutamate are shown in Figs. 1l, 2c and Supplementary Fig. S26c.

### Circadian and homeo3static modulation of calcium in glia

We observed 24-hour calcium fluctuations associated with circadian activity in astrocytes and ensheathing glia when analyzing the averaged normalized calcium activity over 24 hours (Fig. 2d, e, and g). To distinguish between homeostatic and circadian contributions in calcium activity, we first applied a 48-hour lowpass filter to eliminate bleaching effects. Next, we used a 12-hour highpass filter to capture fast calcium modulations related to sleep homeostasis and a 12-hour lowpass filter to isolate circadian modulation. The contributions of each modulation type were calculated by comparing the ranges of the filtered signals to that of the original signal, obtaining the contribution percentage on calcium activity, shown in Fig. 2f (magenta for the circadian contribution and blue for the homeostatic contribution).

**Table S2.**
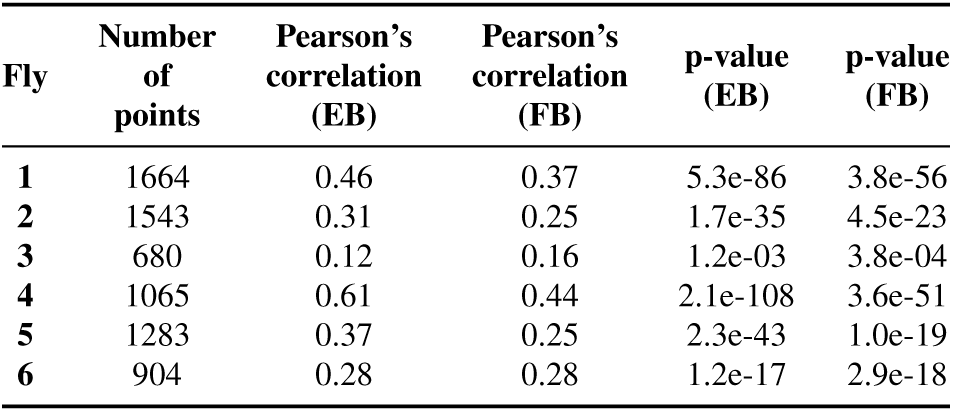
Pearson’s correlation between the changes in high frequency fluorescence fluctuations (change in DF/F) and changes in velocity.

| Fly | Number of points | Pearson's correlation (EB) | Pearson's correlation (FB) | p-value (EB) | p-value (FB) |
| --- | --- | --- | --- | --- | --- |
| 1 | 1664 | 0.46 | 0.37 | 5.3e-86 | 3.8e-56 |
| 2 | 1543 | 0.31 | 0.25 | 1.7e-35 | 4.5e-23 |
| 3 | 680 | 0.12 | 0.16 | 1.2e-03 | 3.8e-04 |
| 4 | 1065 | 0.61 | 0.44 | 2.1e-108 | 3.6e-51 |
| 5 | 1283 | 0.37 | 0.25 | 2.3e-43 | 1.0e-19 |
| 6 | 904 | 0.28 | 0.28 | 1.2e-17 | 2.9e-18 |

### GCaMP sensor saturation

Calcium activity follows exponential saturation dynamics during walking behavior. To determine if this saturation is caused by glia calcium dynamics or if it is an artifact of GCaMP sensor saturation, we analyzed calcium activity during active bouts in the EB and FB of ensheathing glia (Fig. 1i and Supplementary Fig. S3) and astrocytes (Fig. S11a and Supplementary Fig. S11). We used the rising exponentials fitted to the normalized fluorescence data of active bouts (see section above) and calculated the saturation levels, as shown in Supplementary Fig. S10a. Next, we computed the distribution of saturation levels across all active bouts for ensheathing glia (Supplementary Fig. S11b) and astrocytes (Supplementary Fig. S11c). The distributions show that saturation levels are below 1 (the maximum normalized fluorescence value), indicating that calcium saturates during some active periods at values lower than the maximum value of the sensor. This confirms that the saturation observed is due to calcium dynamics, not sensor limitations.

### Velocity modulation of ensheathing glia activity

To assess the influence of fly velocity at short timescales in calcium activity of ensheathing glia in the EB and FB, we computed the mean velocity of the fly during each 1 second imaging epoch, and used a high-pass filter to calculate fluorescence oscillations below periods of 0.5 hours. Supplementary Fig. S18a shows the mean velocity of fly 1 in each trial (black) as well as the high-pass filtered fluorescence in EB (green) and FB (blue). Each data point in Supplementary Fig. S18a was separated by the time difference between consecutive imaging recordings of 60 seconds. We computed the change in mean velocity and the change of DF/F in EB and FB between consecutive epochs, Supplementary Fig. S18b-g. We fitted a linear regression (red line in Supplementary Fig. S18b-g) and computed the Pearson’s correlation between the change in velocity and change in DF/F for the EB and FB, finding a positive correlation for each fly with p-values lower than 0.05 (see Fig. S18i and Table S2).

### Fast calcium modulations in ensheathing glia

To investigate calcium dynamics during the transition between wakefulness and sleep, we recorded calcium activity in ensheathing glia from the FB at a high temporal resolution (60 Hz) while applying mechanical stimulation by opening and closing the ball air valve every second. Fig. 3g shows two examples of this experiment. The left panel shows calcium increases during stimulation when the fly was already walking, while the right panel displays a rapid calcium peak when the fly was asleep prior to stimulation. This phenomenon was observed across ten trials from seven flies, with a significant positive correlation (p < 0.05) between the amplitude of the calcium peak and the duration of rest before stimulation (Fig. 3h).

### Sleep deprivation

Sleep deprivation was performed for recordings in ensheathing glia in the EB and FB. Mechanical sleep deprivation was performed in a total of 5 flies between 3.6 and 8.2 hours, by opening and closing the air stream of the ball repeatedly 3 times for 6 seconds (alternating between 1 second closed and 1 second open) every 20 seconds. This stimulated flies to walk and prevented sleep. Food deprivation was performed for a total of 4 flies between 4 and 16 hours. Individual recordings for both conditions are shown in Supplementary Fig. S21a and b, respectively, where exponentials were fitted for visualization (equation (2)).

For each fly, we computed the ’stop’ state of the fly (zero velocity) in 1 second bins. We then averaged the ’stop’ time for all flies over 1.5 hours before sleep deprivation and over 8 hours of sleep deprivation (left side, first row in Fig. 4b and d). We computed the average of the fluorescence in the EB and FB, which were first normalized through equation (1) (left side, second and third row in Fig. 4b and d, respectively). At least two flies were considered for the average over the sleep deprivation period. The average ’stop’ time and fluorescence were also computed over 2 hours after sleep deprivation (right side in Fig. 4b and d). We then computed the distribution of the mean values of the normalized fluorescence in each fly over 1.5 hours before sleep deprivation, over the first 2 hours of sleep deprivation, after the first 2 hours of sleep deprivation, and over 2 hours after sleep deprivation (Fig. 4c and e). Statistical significance between these distributions was assessed using t-test (p-value < 0.05).

We also calculated the distribution of ’stop’ time after 2 hours of sleep deprivation from the ’stop’ state of each fly, as well as the duration of bouts of immobility that were larger than 5 minutes (Fig. 4f). For control flies, we used the recordings from Supplementary Fig. S3, which had identical conditions as sleep-deprived flies except for sleep deprivation. We then computed the distribution of ’stop’ time over 2 hours at the time of the day when sleep deprivation finished for the sleep-deprived flies, as well as the distribution of sleep bout duration for all flies (Fig. 4f). Statistical difference between distributions was assessed again using t-test (p-value < 0.05).

### Homeostat model

The homeostat model was fitted to activity of glia in the EB, FB, LAL, MB, and midline, as well as activity of R5 neurons, and dFB neurons. For model fitting, fluorescence was filtered and normalized as described before (equation (1)). In the homeostat model we did not distinguish between immobility and sleep, but distinguished two behavioral states based on the fly’s walking activity for charging and resetting of the homeostat, respectively: ’walk’ and ’stop’. The homeostat model was fitted over the time range of each experiment, defined by [*T_min_, T_max_*] for ’stop’ and ’walk’ periods of the fly. The ’stop’ behavior of the fly, *s*(*t*), was set to 1 when the velocity of the fly was below a threshold of (0.25 body lengths per second, to remove tracking noise level) and a value of 0 otherwise. Conversely, the ’walk’ state of the fly, *w*(*t*), was assigned a value of 1 for velocities higher than the threshold and 0 otherwise (top rows in Fig. 5b, Supplementary Figs. S22, S23, S24, S37, S38, S39, S46 and S47). The time resolution for distinguishing between ’stop’ and ’walk’ was 1 second. The following model was used for fitting:

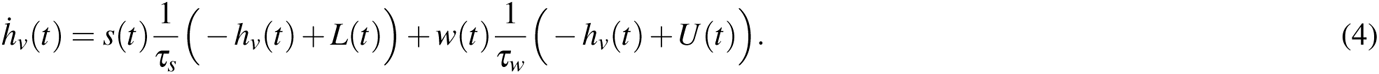

Here, *h_v_*(*t*) describes the fluorescence signal (D*F/F*) or homeostat, while *s*(*t*) and *w*(*t*) act as binary weights for each behavior. Therefore only one behavioral state contributes to the homeostat at any given time. *t_s_* and *t_w_*are the time constants for the stop and walk states, and *L*(*t*) and *U* (*t*) are functions that describe the lower and upper bounds of the homeostat, respectively (yellow lines in Fig. 5b and C, Supplementary Figs. S22, S23, S24, S37, S38, S39, S46 and S47). These bounds were allowed to vary with time to take into account slow modulations in fluorescence that are unavoidable during long-term imaging recordings, such as photobleaching, circadian modulation, or slow changes in calcium levels potentially due to phototoxicity. These changes affect both baseline levels as well as the dynamic range of fluorescence signals over time. Therefore these upper and lower bounds allowed to correct for dynamic range and baseline changes, and were defined as Bezier curves:

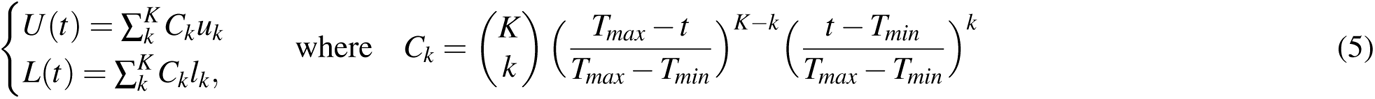

These curves were fitted at the same time as the parameters of the model. The Bezier curves were parameterized by points separated by at least 6 hours, which defined the number of parameters *K* for each experiment as

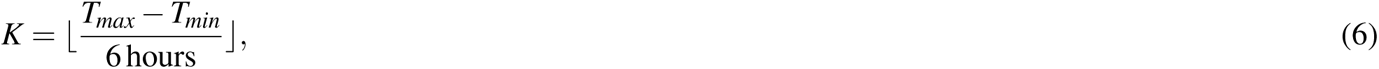

where ⌊.⌋ indicates the floor division operation. For each experiment we fitted the 2 time constants *t_s_*and *t_w_*, and the *K* upper and lower bound parameters, *u_k_* and *l_k_*: *{t_s_, t_w_, u*_1_*, …, u_K_, l*_1_*, …, l_k_}*

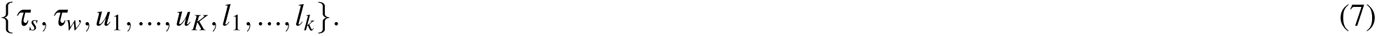

The fitting procedure is described in the next section. The fitted homeostat model for glia activity is shown in Fig. 5b and c and Supplementary Fig. S22 for EB and FB, S23 for LAL, and S24 for MB and midline; for astrocytes in Fig. 5c and Supplementary Fig. S25; for R5 neurons in Supplementary Figs. S37, S38 and S39; and for dFB neurons in Supplementary Figs. S46 and S47. All the fitted time constants for stop and walk and their estimated errors are shown on the left side of all the previous Supplementary Figs., and the combined time constants for glia are shown in Fig. 5e for EB and FB and Supplementary Fig. S29c for EB, FB, LAL, MB, and midline, where asterisks show statistical significance (p-values lower than 0.05) between the walk and stop states in the EB (green) and FB (blue) using t-test.

#### Three-state homeostat model

This model was only fitted for glia activity in the EB and FB, where fluorescence was filtered and normalized as described before (equation (1)). In this model, we distinguished two states for resetting the homeostat: a ’stop’ state where the fly was at rest (as assessed by ball velocity) for less than 5 minutes, and a sleep state, where the fly was at rest for epochs lasting more than 5 minutes.

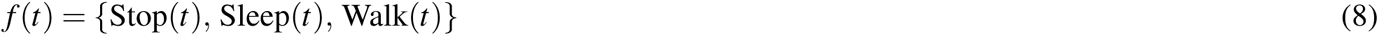

The following model was used for fitting:

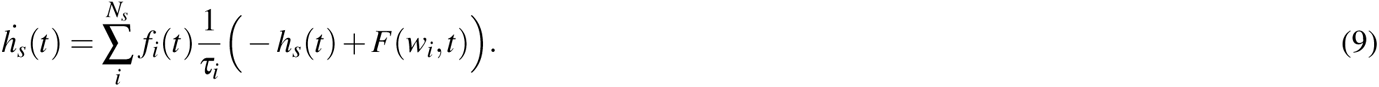

Here, *h_s_*(*t*) describes the fluorescence signal (D*F/F*) or homeostat, and *f_i_*(*t*) acts as a mask for each state, and therefore only the assigned behavior contributes to the homeostat at any given time (similar to the 2-state model). *t_i_* is a time constant for each behavior *i*, and *w_i_* is the weight with which each behavior contributes to the homeostat, -1 for ’stop’ and ’sleep’ states and +1 for ’wake’. *F* is a function that describes the upper and lower bounds of the homeostat,

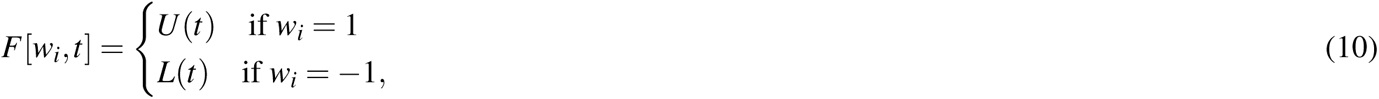

where functions *U* (*t*) and *L*(*t*) are the upper and lower bounds defined in equation (5), similarly to the previous homeostat model.

For each model we fitted the *N_s_* time constants *t_i_* and the *K* upper and lower bound parameters, *u_k_* and *l_k_*:

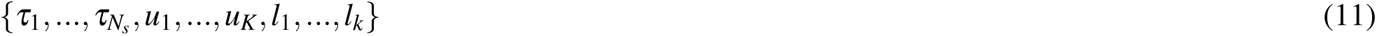

An example of model fitting for glia activity in the EB and FB is shown in Supplementary Fig. S33a. The fitted time constants for each of the states, as well as their estimated errors in EB and FB, are shown in Supplementary Fig. S33b. The distribution of time constants for each state in the EB and FB is shown in Supplementary Fig. S33c. Only time constants with an estimated error lower than 0.2 times its fitted value were included to discard estimated time constants with high error. Asterisks in Supplementary Fig. S33c indicate statistical significance (p-values lower than 0.05) between different behaviors in EB (green) and FB (blue) using t-test.

We also asked if the homeostat would reset differently after 5 minutes of immobility. For this purpose, we defined a sleep state only after the fly was stopped for 5 minutes and fitted again equation 9. Supplementary Fig. S34a shows an example for glia activity in the EB and FB, together with the fitted time constants in Supplementary Fig. S34b. The distribution for time constants with an estimated error lower than 0.2 times its value are shown in Supplementary Fig. S34c for EB and FB. We did not find statistical significance between the time constants of the ’sleep’ and ’stop’ states using t-test.

#### Seven-state homeostat model

Activity of glia in the EB and FB, which was filtered and normalized as described before (equation (1)), was fitted to a homeostat model using the classified behaviors (an example is shown in the top row of Supplementary Fig. S31b.

We fitted the dynamics of the homeostat with a model taking into account the *N_b_*= 7 classified behaviors from the 3D CNN:

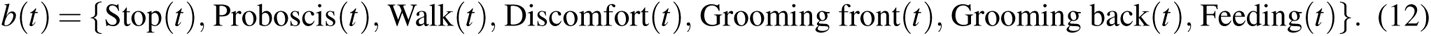

Each behavior, *b_i_*(*t*), was assigned a binary value, *b_i_*(*t*) *2 {*0, 1*}*, and only one behavior had value 1 at any given time, corresponding to the maximum class value predicted by the 3D CNN. The time resolution for the classification of behavior was 1 second.

We used the following model to fit the data:

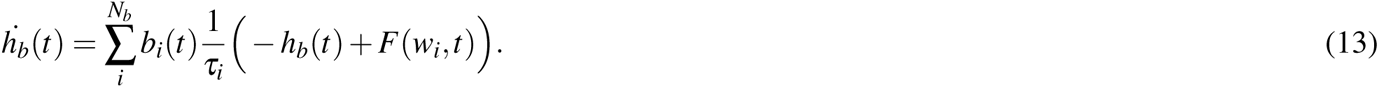

Here, *h_b_*(*t*) describes the fluorescence signal (D*F/F*) or homeostat, and *b_i_*(*t*) acts as a mask for each behavior, and therefore only the assigned behavior contributes to the homeostat at any given time (similar to the 2-state model). *t_i_* is a time constant for each behavior *i*, and *w_i_* is the weight with which each behavior contributes to the homeostat, which can take only two values: *w_i_ 2 {-*1, 1*}*. When a weight is *w_i_* = *-*1, the homeostat decreases while the fly performs the behavior, while with *w_i_* = 1, the homeostat increases. *F* is a function that describes the upper and lower bounds of the homeostat, given by equation (10), similar to the previous 2- and 3-states model.

Fitting this differential equation requires determining for each behavior whether it charges or resets the homeostat, that is, contributes with a negative or positive weight to the model. To determine the weight of each behavior in the model, we used fluorescence traces from flies that performed one of the 7 behaviors for at least two consecutive imaging epochs (120 seconds). For ’discomfort’ we used fluorescence traces recorded when the ball was stopped (Fig. 3c and Supplementary Fig. S16), since this behavior was not observed continuously for 120 seconds when the ball was free to rotate. For the rest of the behaviors, we used fluorescence traces from the recordings of ensheathing glia in 6 flies. For each behavior, all traces were aligned at the origin (thin lines in Supplementary Fig. S31a for the FB) and linear regression was used to determine the slope. If the slope was positive for a behavior, the weight was set to 1 and otherwise to -1 (negative slope). We repeated the same procedure for FB and found the same weights as in EB (magenta lines in Supplementary Fig. S31a). For each model we fitted the *N_b_* time constants *t_i_* and the *K* upper and lower bound parameters, *u_k_* and *l_k_*:

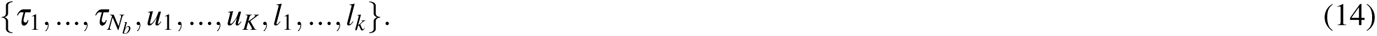

Models were fitted independently to EB and FB data.

The fitting procedure is explained in the next section. Since some behaviors were very rare in some flies, the estimation error for model fitting was large for some of the time constants. Therefore only time constants with an estimated error lower than 0.2 times its fitted value were included in the distribution of the time constants for each behavior in Supplementary Fig. S32a. Asterisks in Supplementary Fig. S32a indicate statistical significance (p-values lower than 0.05) for behaviors in the EB (green) and FB (blue) using t-test.

#### Hunger-walk model

We analyzed the dynamics of dFB activity with respect to feeding by fitting exponentials (see below for a more detailed model). The average fluorescence traces were divided into two epochs, one where the fly was hungry (defined as 3.5 hours before the next feeding event) and one where the fly was fed (defined as 30 minutes after feeding, see Fig. 7a, b, Supplementary Fig. S48, and Methods). The time constants during epochs where the fly was hungry were in the order of hours (Fig. 7c). The time constants for resetting after feeding were in the range of minutes, but they might not reflect the actual dynamics of dFB neurons after feeding since resetting depended also on the fly behavioral state (Fig. 6j-m).

For a more detailed model-based analysis, fluorescence traces were first filtered and normalized as described before (equation (1)). The model contained two different components that contributed differently: a hunger component, and a walk-stop component. The hunger component was fitted based on the time relative to the feeding events. We defined a "hungry" variable, hungry(*t*), defined as 0 between a feeding event and a time after feeding, called satiated time *t_satiated_*, and as 1 after the satiated time and the next feeding event. Another variable, fed(*t*), was defined as 1 after feeding and before the satiated time, and as 0 otherwise. Therefore fed(*t*)= 1 *-* hungry(*t*). The satiated time, *t_satiated_*, interpreted as the time during which the hunger component resets and stays low, was fitted as a model parameter. The hunger component, *h_hunger_*, was therefore fitted using the following differential equation:

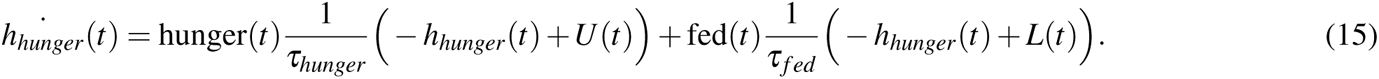

Here, hungry(*t*) and fed(*t*) act as a mask and therefore only the assigned variable contributes to the hunger component at any given time (similar to the 2-state model). *t_hungry_* and *t_f ed_* are the time constants that describe the increase and decrease of the hunger component, respectively. The hunger component and its parameters are shown in Fig. 7f. *U* (*t*) and *L*(*t*) represent the upper and lower bounds, defined in equation (5), which were fitted together with the parameters. The walk-stop component, *h_v_*(*t*), was defined as in the homeostat model (equation (4), but the upper and lower bouts were set constant, *U* (*t*)= 1 and *L*(*t*)= 0.

Finally, the hunger-walk model was defined as the weighted sum of the hunger and walk-stop components as follows:

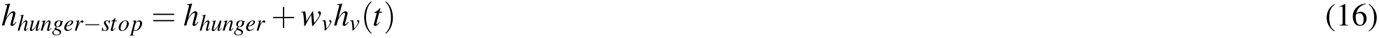

where *w_v_*was a weight that indicated the contribution of the walk-stop component, which was also fitted. In summary, for this model we fitted the following 6 parameters,

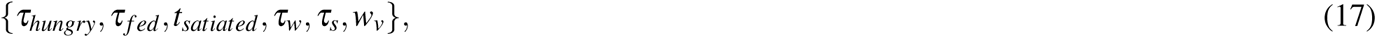

as well as the parameters that defined the upper and lower bounds of the hunger component, *u_k_* and *l_k_* from equation (5). The fitting procedure is explained in the next section, and the fitted models are shown in Fig. 7g and Supplementary Figs. S51 and S52. The fitted time constants of all flies are shown in Fig. 7h, where only parameters with an estimated error lower than 0.2 times their fitted value were included. Asterisks in Supplementary Fig. 7h indicate statistical significance (p-values lower than 0.05) for behaviors in the EB (green) and FB (blue) using t-test.

#### Model fitting

Fitting of models was performed using the function *curve fit* in Scipy^135^ in Python. Given a set of initial values of the parameters, we integrated each model with the corresponding equation, for example using equation 4 for the 2-state model, equation 9 for the 3-state model and equation 13 for the 7-state model, from their corresponding behavioral states (’stop’, ’walk’, …) over the experiment timeline using Euler integration (with 1 second steps). Since the fluorescence data was not recorded continuously, we then interpolated the values of the model in agreement with the times of the recordings. Finally, we obtained an *L*2-error function between the interpolated model values and fluorescence data. The parameter values were updated iteratively from the Jacobian of the error function, and the minimum was found using the trust region reflective algorithm as optimization method.

To prevent parameters from taking forbidden values during optimization, we used parameter bounds. For the homeostat model, as well as the 3-state and 7-state homeostat models, the allowed range for time constants was [0, 2] hours. For the hunger-walk model, time constants in the range of [0, 2] hours were allowed, except for the time constant for "hungry", _*tau_hungry_*, where the allowed range was [0, 6] hours. The allowed range for the weight *w_v_* in the hunger model was [0,1], and the allowed range for the upper and lower bound parameters, *u_k_* and *l_k_*, were [*-*1, 0] and [0, 2] normalized D*F/F*, respectively.

We computed the error or variance of each parameter from the covariance matrix between all parameters, which was returned by the function *curve fit*. This error indicated one standard deviation error of the parameter, obtained as the square root of the corresponding diagonal element in the covariance matrix. In all models, parameters with errors lower than 0.2 times the magnitude of the fitted parameter were considered for distribution analysis (Fig. 5e, 1h and Supplementary Figs. S33, S34 and S32a).

#### Comparison between models

We asked whether the 7-state homeostat model fitted the glia activity better than the homeostat model. In this case, the homeostat model is a nested model, the 7-state homeostat model, which means that the homeostat model and its parameters are included in the 7-state homeostat model and parameters. We therefore asked if a null hypothesis holds, which establishes that the more complex (7-state homeostat) model does not fit fluorescence data significantly better than the simpler model (homeostat model). For this purpose we used a F-test^136^.. Since the more complex model has different numbers of parameters than the simpler model, we can compute the F-statistic of the F-test, as:

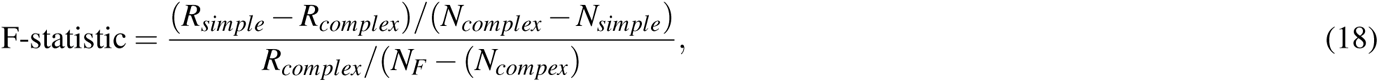

where *R_simple_*and *R_complex_*are the residual sum of squares for the simpler and the more complex model, respectively (same as *L*2-error), *N_simple_* and *N_complex_* are the number of parameters for the simpler and more complex model, respectively (in this case *N_complex_ - N_homeostat_* = 5) and *N_F_* is the number of fluorescence points that was used for fitting both models. The F-statistic value can then be used to generate a p-value, rejecting the null hypothesis stated above if it is lower than 0.05. If the null hypothesis is rejected, we can conclude that the more complex model fits the data significantly better than the simpler model.

The L2-errors between each pair of models for each fly in EB and FB are shown in Fig. S32b, while the L2-errors, F-statistic values, and p-values comparing each pair of models in EB and FB are shown in Tables S3, S4. This analysis concludes that the 7-state homeostat model provides a better fit for the glia data than the homeostat model.

We performed a similar analysis to compare the homeostat model with the hunger-walk model fitted to dFB neurons. As before, the homeostat model was a nested model of the hunger-walk model. We computed the F-statistic between the two models (being the homeostat model the simpler model, while the hunger-walk model is the more complex model) from equation 18 and obtained the corresponding p-values. The L2-errors between each pair of models fitted for each fly in shown in Fig. 7i, while the L2-errors, F-statistic values, and p-values are shown in Table S5. Since all the p-values were lower than 0.05, we conclude that the hunger-walk model fits the activity of dFB neurons better than the homeostat model.

**Table S3.**
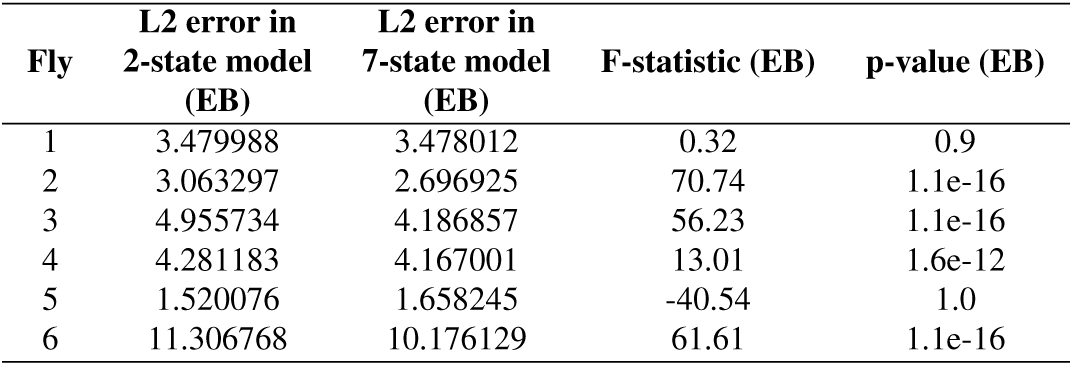
Comparison between 2-state and 7-state models in the EB.

| Fly | L2 error in<br>2-state model<br>(EB) | L2 error in<br>7-state model<br>(EB) | F-statistic (EB) | p-value (EB) |
| --- | --- | --- | --- | --- |
| 1 | 3.479988 | 3.478012 | 0.32 | 0.9 |
| 2 | 3.063297 | 2.696925 | 70.74 | 1.1e-16 |
| 3 | 4.955734 | 4.186857 | 56.23 | 1.1e-16 |
| 4 | 4.281183 | 4.167001 | 13.01 | 1.6e-12 |
| 5 | 1.520076 | 1.658245 | -40.54 | 1.0 |
| 6 | 11.306768 | 10.176129 | 61.61 | 1.1e-16 |

**Table S4.** Comparison between the 2-state and 7-state models in the FB.

| Fly | L2 error in<br>2-state model<br>(FB) | L2 error in<br>7-state model<br>(FB) | F-statistic (FB) | p-value (FB) |
| --- | --- | --- | --- | --- |
| 1 | 3.588580 | 3.246188 | 59.59 | 1.1e-16 |
| 2 | 0.784993 | 0.733569 | 36.50 | 1.1e-16 |
| 3 | 3.402890 | 2.818227 | 63.52 | 1.1e-16 |
| 4 | 1.604965 | 1.512774 | 28.95 | 1.1e-16 |
| 5 | 2.616210 | 2.554074 | 11.84 | 2.4e-11 |
| 6 | 5.747171 | 5.153419 | 63.90 | 1.1e-16 |

**Table S5.**
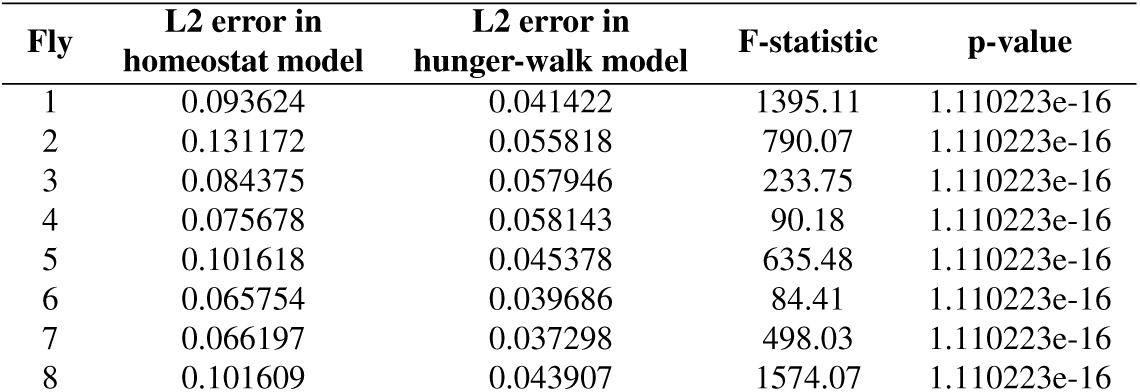
Comparison between the homeostat model and hunger-walk model for dFB neurons.

| Fly | L2 error in<br>homeostat model | L2 error in<br>hunger-walk model | F-statistic | p-value |
| --- | --- | --- | --- | --- |
| 1 | 0.093624 | 0.041422 | 1395.11 | 1.110223e-16 |
| 2 | 0.131172 | 0.055818 | 790.07 | 1.110223e-16 |
| 3 | 0.084375 | 0.057946 | 233.75 | 1.110223e-16 |
| 4 | 0.075678 | 0.058143 | 90.18 | 1.110223e-16 |
| 5 | 0.101618 | 0.045378 | 635.48 | 1.110223e-16 |
| 6 | 0.065754 | 0.039686 | 84.41 | 1.110223e-16 |
| 7 | 0.066197 | 0.037298 | 498.03 | 1.110223e-16 |
| 8 | 0.101609 | 0.043907 | 1574.07 | 1.110223e-16 |

#### Correlation between ’convolved walk’ and calcium activity

To compare how ensheathing glia calcium activity, and R5 and dFB neurons integrate wake time in the different compartments, we first computed the convolution of the binary ’walk’ state of the fly, *w*(*t*) (assigned with a value of 1 for velocities higher than the threshold and 0 otherwise), with a temporal filter, *T_f_* (*t*). The temporal filter had a triangular shape (second row in Fig. 6h) defined by a filter size, *s _f_* , as:

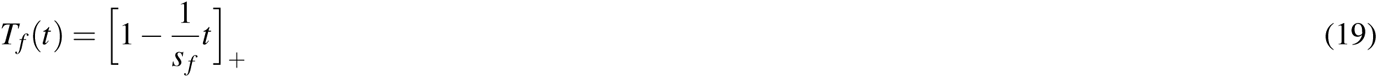

**Table S6.**
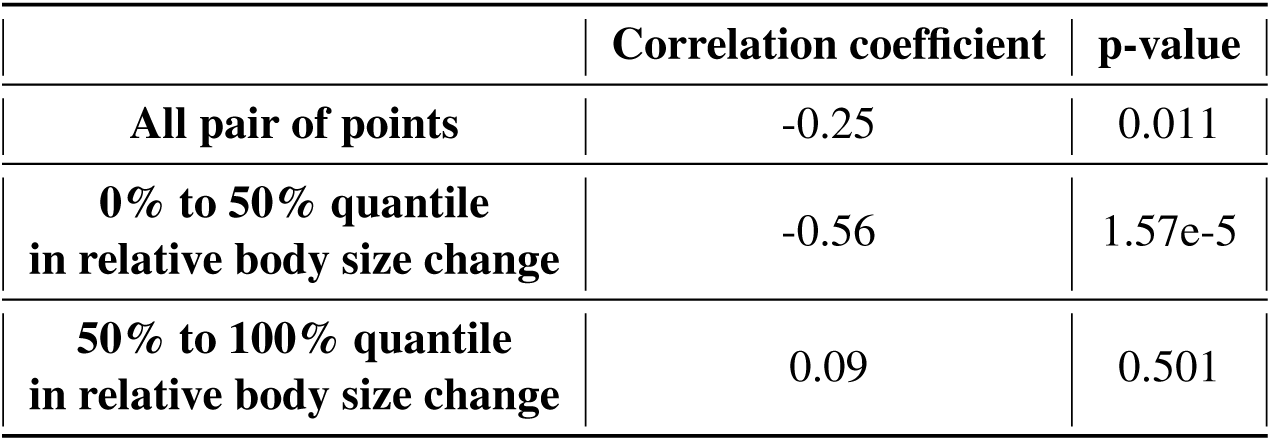
Pearson correlation and p-values for the relative body size changes and changes in activity of dFB neurons before and after feeding events.

|  | Correlation coefficient | p-value |
| --- | --- | --- |
| All pair of points | -0.25 | 0.011 |
| 0% to 50% quantile<br>in relative body size change | -0.56 | 1.57e-5 |
| 50% to 100% quantile<br>in relative body size change | 0.09 | 0.501 |

where ⌊.⌋ represents a threshold-linear function to ensure only positive values. The convolution between the ’walk’ state and the triangular filter produced the ’convolved walk’ (third row in Fig. 6h), which exponentially increased when the ’walk’ state was 1 and exponentially decreased otherwise. The rate of increase and decrease was the same, defined by the filter size *s _f_* . The ’convolved walk’ can therefore be used to compute the Pearson correlation with the normalized fluorescence (as previously described, see equation (1)) for different filter sizes *s _f_* . We used a total of 34 temporal filters, ranging from 0 to 60 minutes in steps of 6 minutes, and from 1 hour to 25 hours in steps of 1 hour. The average correlation for ensheathing glia in EB, FB, LAL, MB, and midline is shown with colored lines in Fig. S29a, while the standard deviation of the correlation from each group is represented by semi-transparent colored regions around the average in Fig. S29a. Only filter sizes lower than 240 minutes were considered, as larger filter sizes produced close to zero correlation. We obtained the maximum value of the averaged correlation and its corresponding filter size (colored numbers in Fig. S29a) and assessed the statistical difference between these maxima using t-test. Statistical significance was indicated by asterisks in Fig. S29a (p-values < 0.05 represented by one asterisk, and p-values < 0.005 represented by two asterisks). The same analysis was performed for glutamate recordings, shown in Supplementary Fig. 6b, as well as for R5 and dFB neurons to compare with glia activity in the EB and FB (Fig. 6i).

#### Comparison between activity of ensheathing glia and neurons

The dynamics of activity in ensheathing glia, R5, and dFB neurons were compared with respect to how well they described a sleep homeostat. For this, we used three different methods. In a first approach, we reasoned that a homeostatic signal should increase over active epochs, producing a positive correlation with the time during which the fly was active, and decrease over epochs of rest or sleep, thus producing a negative correlation with time spent resting or sleeping. We therefore computed the Pearson correlation between the averaged normalized traces over 30 minutes of active and rest epochs (Figs. 1j, k, 6c, d, Supplementary Figs. S6, S36, S40, and S45). We distinguished between the EB and FB neuropils and compared the distributions of correlation values in the EB between glia and R5 neurons and the distribution of correlation values in the FB between glia and dFB neurons (Fig. 6e, f). To assess whether the distributions of correlation values were statistically different we used t-test. Asterisks in Fig. 6e and f represent p-values lower than 0.05.

Alternatively, we compared the normalized average traces to a linear fit (Figs. 1j, k, 6c, d, Supplementary Figs. S6, S36, S40, and S45) during the first 30 minutes of rest and activity. The slope of this fit for each fly, shown in Fig. S53, represents how fast activity increases towards a maximum value during active epochs (Fig. S53, left side) and how fast activity decreases during epochs of rest to baseline (Fig. S53, right side). Statistical differences of the slope distributions were again assessed using t-test and are represented by asterisks (p-value < 0.05) in Fig. S53. The slope for a homeostatic signal that increases and decreases linearly over 30 minutes of activity or rest should be 1 and -1, respectively. Glia activity was very close to these values (Fig. S53), while the slope for dFB neurons during rest was close to zero.

As a second approach, we calculated the *L*2-error of the homeostat model fitted for glia and R5 neurons. We again distinguished between neuropils and compared the *L*2-error of glia in the EB and of R5 neurons, as well as glia in the FB and dFB neurons (Fig. 6g). Statistical significance was assessed using t-test, where asterisks indicate that p-values were lower than 0.05.

As a third approach, we computed the Pearson correlation between the normalized fluorescence and the ’convolved walk’ (third row in Fig. 6h) for different filter sizes (equation (19), as described previously. A total of 34 temporal filters were used, ranging from 0 to 60 minutes in steps of 6 minutes, and from 1 hour to 25 hours in steps of 1 hour. The average correlation for glia in the EB and R5 neurons, and for glia in the FB and dFB neurons, is shown with colored lines in Fig. 6i, while the standard deviation of the correlation from each group is represented by semi-transparent colored regions around the average in Fig. 6i. Filter sizes larger than 240 minutes are not shown, since they produced close to zero correlation. The maximum value of the averaged correlation was obtained together with its corresponding filter size (colored numbers in Fig. 6i). Statistical significance between these maxima was assessed using t-test, indicated by asterisks in Fig. 6i (p-values < 0.05).

#### Fitting of fluorescence traces during ’hungry’ and ’fed’ states

This analysis was only performed for dFB neurons and was similar to the analysis using active and rest epochs. We defined ’hunger’ and ’fed’ epochs to characterize the trend of activity of dFB neurons. Fed epochs were defined over the first 30 minutes after a feeding event, while hungry epochs were defined as starting 30 minutes after a feeding event and lasting until the next feeding. At least 4 normalized fluorescence traces were used to obtain the normalized average fluorescence over hungry or fed states, which was fitted with a rising exponential (equation (2)) or a resetting exponential (equation (3)), respectively (Fig. 7b and Supplementary Fig. S48). The time constants of the fitted exponentials are shown in Fig. 7c, where the asterisk indicates statistical significance (p-value < 0.05) using t-test.

#### Change in body size before and after feeding

This analysis was only performed for dFB neurons. We asked whether the reset of activity of dFB neurons was linked with how much food flies were ingesting at each feeding event. We therefore used the side view of the fly in the video recorded during the experiment 10 minutes before feeding and 10 minutes after feeding. Using color segmentation in OpenCV, we obtained a mask of the fly that only included the abdomen and the wings (Fig. 7d). We did not include the legs or the head in this mask, as flies could for example move the legs or extend the proboscis, producing an enlarged body size. We computed the area of the mask before and after feeding, *A_be f ore_* and *A_a fter_*, respectively, and calculated the relative body size change between feeding events, D*A_r_*, as follows:

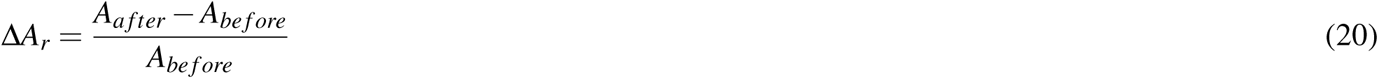

The relative body size changes between feeding events were then compared to the corresponding differences in activity of dFB neurons before and after feeding. For this, we averaged the activity of dFB neurons over the last 10 minutes before a feeding event, *< F >_be f ore_*, and over the next 10 minutes after feeding, *< F >_a fter_*, and obtained the difference, *<* D*F >*=*< F >_be f ore_ - < F >_a fter_*. We obtained a total of 107 pairs of relative body size changes and fluorescence changes from 8 flies, shown in Fig. 7e. We computed the Pearson correlation between all pairs of points, as well as during the first half of the lowest relative body size changes (50% quantile) and the second half, shown in Table S6. While the correlation was positive and statistically significant (p-value < 0.05), this correlation was stronger for lower relative body size changes (Table S6).

#### EM data analysis

We downloaded a portion of the raw EM data from the hemibrain dataset^37,137^. The analyzed data, containing EB and FB, was a cube with dimensions of 136*µm*, centered at the coordinates (25711, 27000, 19798). Although the hemibrain dataset contains the segmentation of glia matter and tracheal tubes, we noticed that a custom glia mask for each slice, obtained by selecting those parts of the image where neurons were not segmented, resulted in more reliable results. Therefore, we used the available segmentation of neurons in each slice and defined the glia matter as the non-segmented regions. An example of this approach for the segmentation of glia matter for one slice is shown in Fig. 8a (right side, glia in yellow).

Tracheal tubes appear in a different color and we used a simple color threshold segmentation method. Once all slices were segmented, mislabeled non-tracheal structures were manually removed using the software Blender. Fig. 8a shows on the right in red the segmentation of the tracheal tubes for one slice. Finally, we used the available hemibrain defined neuropil boundaries to get the EB and FB masks (green and blue, respectively, in Fig. 8a, right side).

To compute whether tracheal tubes were always found inside glia, we compute the overlap between each of the tracheal tubes and the glia matter for each slice. If the overlap was non-zero, the tracheal tube was considered to be inside the glia mask. Fig 8b shows the amount of tracheal tubes inside (green) and outside (red) glia for each slice (left side), and the distribution of tracheal tubes across all slices inside and outside glia (right side). Statistical significance between tracheal tubes inside and outside glia was addressed using the Kolmogorow-Smirnow test.

To compute the distance between tracheal tubes and the boundaries of the EB and FB neuropils, we generated a 3D distance transform for each neuropil using Python’s *edt* library. This process produced a 3D distance map for each neuropil, which is a 3D image where the voxel values indicated the distance to the nearest neuropil boundary. Voxels located outside the neuropil had positive distance values, while those inside had negative values. Next, for each slice, we identified the center of each tracheal tube using the tracheal mask and determined its distance to the EB and FB neuropils based on its position in the 3D distance map. After calculating the distances of the tracheal tubes for each slice, we grouped them into 1*µm* bins to create a distribution of the number of tracheal tubes for each binned distance. The bottom panel of Fig. 8c shows this distribution, with slices along the y-axis and the binned distances along the x-axis for both the EB (left) and FB (right) neuropils. For slices that did not intersect with the neuropil (above or below its boundaries), negative distances (representing positions inside the neuropil) are shown in white. This indicates that negative distances are not possible in these slices, as no part of the neuropil is present. The top panel of Fig. 8c shows the average distribution of tracheal tubes across slices for the EB (left) and FB (right)l that contain at least one tracheal tube. Specifically, for each distance bin, we calculated the average number of tracheal tubes, considering only the slices where tracheal tubes are present. This approach offers a clearer view of the number of tracheal tubes near the boundaries by focusing on slices that contain tracheal tubes. Tracheal tubes were rarely observed inside the neuropil (at negative distances) and were primarily concentrated around its boundary (at zero distance, see Fig.8c). This suggests that the tracheal tubes are embedded in ensheathing glia, which are located at the neuropil boundary.

#### Gas exposure during imaging

To deliver gas to flies during imaging, we used an outlet at a distance of approximately 1*cm* from the fly positioned to the side (Fig. S54e). The outlet delivered either air, oxygen (100 % concentration), CO_2_ (at three different concentrations: 10 %, 50% or 100% in air), or ammonia. For ammonia, air was humidified with a solution that contained a concentration of 5% ammonia (NH_3_) diluted in water. The CO_2_ concentration was obtained by mixing different volumes of air with CO_2_. A suction outlet running at 1 liter per minute was positioned opposite the inlet, on the other side of the fly and, at a distance of approximately 1*cm* from the fly (Fig. 8e). The outlet was connected through a tube to a sensor (SPRINTIR-WF-20 sensor) to measure the concentration of CO_2_ (first row, blue line in 8f).

Switching between the different gases was performed through a system of automated air valves, which were controlled using an Arduino MEGA board. The experiment consisted of delivering first oxygen (for 5 minutes), then CO_2_ (for 10 minutes for each different concentration), and then ammonia (for 5 minutes). Between these gases, air was delivered (Fig 8f). All gases were delivered at a rate of 0.5 liters per minute.

Flies expressing either jGCaMP8m or pHluorinSE on ensheathing glia (56F03-GAL4), astrocytes (86E01-GAL4), or neurons (57C07-GAL4) were used during gas exposure experiments (sample size is shown in Fig. 8f).

We calculated the percentage of immobile flies during gas exposure in 10-second bins, as well as proboscis extensions per minute (black and red lines, Fig. 8f). The average fluorescence traces from jGCaMP8m and pHluorinSE in neurons, astrocytes, and ensheathing glia are shown in the last 3 rows of Fig. 8f for the FB and Supplementary Fig. S54a for the EB (black line for pHluorinSE, green line for jGCaMP8m). We analyzed the distribution of immobile flies during (Fig. 8g) and after gas exposure (Fig. 8h). Immobility was significantly higher during 100% CO_2_ exposure, and after all CO_2_ concentrations and ammonia. Proboscis extension significantly increased during 50% and 100% CO_2_ exposure (Fig. 8i). Statistical significance in Fig. 8h-i is indicated by asterisks using t-tests (p < 0.05).

We also computed the average pHluorinSE fluorescence during gas exposure in neurons, astrocytes, and ensheathing glia in the FB (black, magenta and green lines in Fig. 8j, respectively) and EB (Supplementary Fig. S54b). The drop in pHluorinSE was largest in ensheathing glia, followed by astrocytes, and smallest in neurons, reflecting a layered hierarchy where glia are most affected due to proximity to the gas. We also computed the pHluorinSE overshoot 5 minutes post-exposure, showing that both CO_2_ and ammonia induced significant overshoots in ensheathing glia and astrocytes (Fig. 8k and Supplementary Fig. 8c).

Next, we measured the jGCaMP8m calcium overshoot (Fig. 8l and Supplementary Fig. 8d), revealing significant differences between ammonia and 50% CO_2_, indicated by colored asterisks (using t-tests). This calcium overshoot is different from the pH overshoot, showing that calcium activity during CO2 exposure is not simply a reflection of pH changes. This difference motivated the use of ammonia, as both CO2 and ammonia raise pH similarly. However, ammonia did not trigger the strong calcium responses observed with CO2, confirming that the jGCaMP8m signal after CO2 exposure reflects actual calcium overshoot dynamics in astrocytes and ensheathing glia. Statistical differences between neurons, astrocytes, and glia were assessed using a one-way ANOVA, as indicated by black asterisks at the top of Fig. 8i-l and Supplementary Fig. S54b-d.

Finally, we calculated the time constants for the decay of pHluorinSE and jGCaMP8m signals following CO_2_exposure in ensheathing glia and astrocytes, both in the EB (Supplementary Fig. S54e-h) and FB (Supplementary Fig. S54i-l). The time constants for both cell types were similar, averaging around 2 minutes.

#### Optogenetics during imaging

Optogenetic activation is achieved by illumination of the select areas of the brain with computer generated holograms, projected with a red laser (640 nm, Toptica, iBEAM-SMART-640-S-HP) and phase-SLM (Meadowlark Optics, HSP1920-1064-HSP8). The activation laser beam is expanded, modulated, and multiplexed into the scanned fluorescence illumination path with a dichroic mirror. The phase SLM, displaying the hologram, is located in the plane conjugate with the back focal plane of the microscope objective (Fig. 9 a).

We use the classic Gerchberg-Saxton algorithm^138^ with 20 iterations, where the desired activation intensity distribution in the field of view serves as the amplitude constraint to find phase modulation in the Fourier plane to be displayed on the SLM.

We have integrated a custom GUI with Scanimage^139^, allowing the user to mark areas of activation by painting over them in the two-photon field-of-view (FOV). To schedule the activation, the laser was controlled by an arduino UNO board to switch it on and off automatically.

To allow for precise activation, a correspondence needs to be established between the 2-photon FOV and the area illuminated by the SLM. This calibration was done with an auxiliary area scan camera (Basler acA640-750um) imaging reflected light and fluorescence from sample with beads through the objective (procedure is shown in Fig. S57).

#### Optogenetics protocol during imaging

We crossed flies expressing R56F03-GAL with UAS-CSChrimson and UAS-jGcAMP8m. The flies were raised on regular food and, after surgery, were placed on retinal food, while control flies remained on regular food for recovery. After a two-day recovery, experiments were conducted using a power of 0.3 mW in the optogenetics laser. We used a total of 6 and 9 flies for ensheathing glia and astrocytes optogenetics activation, and 8 and 9 flies for ensheathing glia and astrocytes controls, respectively.

Trials involved continuous laser activation for varying durations (30, 60, 120, 300, and 600 seconds), followed by 20 minutes of recovery. Before each trial, we mechanically stimulated the flies for 10 seconds to ensure that they were awake. Optogenetics activation was performed on the lower edge of the EB (as defined in Fig. 9a). Figures 9b and c show that flies were less immobile 10 seconds before activation due to mechanical stimulation. We also measured proboscis extension events per second during the trials (red lines in Fig. 9b and c). Calcium dynamics were recorded at the activation site in the EB and in the FB compared to control flies (green, blue and black lines in Fig. 9b and c, respectively).

The percentage of immobile flies during optogenetic activation in ensheathing glia and astrocytes is shown in Fig. 9d and was higher than in control flies. Asterisks at the top of the panels indicate significant differences compared to control, as determined by t-tests (p < 0.05). Additionally, proboscis extensions per second were significantly greater during optogenetic activation in these cell types (Fig. 9e). The inter-proboscis extension period during activation averaged around 1.5 seconds, faster than during natural sleep (Supplementary Fig. S56a).

We analyzed calcium dynamics in the FB during optogenetic activation. For calcium drift or integration, we calculated the 0.95 quantile fluorescence value for ensheathing glia and astrocytes (Fig. 9f). We also measured the calcium overshoot in the FB five minutes post-activation, defined by the 0.95 quantile calcium value (Fig. 9g). Both drift and overshoot were significantly different from control (indicated by black asterisks in Figs. 9f and g). The overshoot amplitude also increased with longer optogenetic activation times in ensheathing glia and astrocytes (indicated by colored asterisks, using t-tests). The same analysis was performed for the EB, as shown in Supplementary Fig. S56b-c. Finally, we computed the time constants for calcium decay after the overshoot, revealing that calcium dynamics were faster in astrocytes compared to ensheathing glia (Fig. S56d). Time constants were also similar to those obtained after CO_2_ exposure (S54e-l).

### 2 Supplementary Information

#### 2.1 Comparison of homeostat activity in different brain areas

Glia activity in the MB and midline showed in some cases weaker correlations with walking activity (Supplementary Fig. S29a and Methods than the central complex. However, intermittently opening and closing the air stream supporting the ball with a valve every second, which induced fast walking activity, reliably induced increased calcium activity in MB and midline similar to the central complex (Supplementary Figs. S19 and S20). That activity in the EB, FB, and LAL was better correlated with walking activity compared to activity in the MB and midline (see Methods and Supplementary Fig. S29a, is likely due to neurons in EB, FB and LAL being important for visual navigation, the behavioral paradigm used in these experiments.

#### 2.2 Glutamate changes faster than calcium

Ensheathing glia are important for glutamatergic signaling^12^ and glutamatergic neurons show activity correlated with walking^140^. We therefore tested whether the slow calcium dynamics observed in glia corresponded to the accumulation of extracellular glutamate. The glutamate sensor iGluSnFR was expressed in ensheathing glia and we performed long-term imaging experiments as before^141,142^.

Glutamate activity was correlated with walking (Supplementary Fig. S26a, S27 and S29b) and we again defined active and rest epochs based on the velocity of the fly and fitted exponential curves for EB and FB fluorescence traces (Supplementary Fig. S26b and S28). Extracellular glutamate increased during active epochs and decreased during rest (Supplementary Fig. S26b). However, time constants were faster than those obtained for calcium (compare Supplementary Fig. S26c and Fig. 1l, and Supplementary Fig. S29a and b), indicating that ensheathing glia calcium levels do not simply reflect extracellular glutamate concentration.

#### 2.3 Homeostat model with two different rest states based on 5 minutes threshold for sleep

Calcium dynamics in ensheathing glia and astrocytes can be described with a simple model based on stop and walk behaviors (Fig. 5). Additionally, we fitted models to ensheathing glia activity to determine whether distinct sleep stages could be inferred from the calcium dynamics. Models for the EB and FB assuming two different rest states, distinguishing between epochs with less and more than 5 minutes of immobility (defining sleep^132^), did not result in significantly different time constants between the two rest state, even when defining a second state as starting only 5 minutes after the onset of immobility (see Supplementary Figs. S33, S34 and Methods). This shows that the homeostat resets similarly in short and long bouts of immobility, consistent with the observation that already short bouts of immobility show features of sleep^18,20^.

#### 2.4 Homeostat integrates multiple behaviors

We also tested whether a more detailed analysis, distinguishing multiple behaviors, could improve model fitting. Behavior on the ball, which was monitored with a camera throughout the experiments, was classified into the following 7 categories using machine learning with 1 second time resolution (similar to^143^, Supplementary Fig. S14): immobility (stop), proboscis extension, walking, discomfort (where the fly was pushing or pulling the ball), grooming of the front, grooming of the back, and feeding (see Methods and Supplementary Video S7).

We used the intervals where flies performed one of the defined behaviors continuously over consecutive imaging epochs to assess whether they contributed to an increase or decrease of calcium activity. As expected, glia activity in the EB and FB decreased when flies were stopped, but also during proboscis extension and grooming. Glia activity increased with walking, feeding (during which flies were often walking), and discomfort (Supplementary Fig. S31a). We next fitted a model to activity in EB and FB to obtain the time constants for each of the seven behaviors (Supplementary Fig. S31b). This model predicts that resetting of the homeostat occurs faster during proboscis extension and back-grooming than during immobility, and also resets during front-grooming, although more slowly than during immobility. Proboscis extension and back-grooming was only observed in short bouts, and rarely over more than two consecutive imaging epochs (Supplementary Fig. S32a). Consistent with these observations, proboscis extension has been described to occur during a deep sleep state, and other brief movements have also been observed during sleep^16,20,21^.

#### 2.5 Correction of jGCaMP8 fluorescence for pH induced changes

Calcium activity in ensheathing glia, measured using the pH-sensitive jGCaMP8m sensor during CO_2_ exposure (Fig. 8f), can be corrected by subtracting the pH component from the signal (see Methodsand Supplementary Fig. S55). This adjustment provides a corrected GCaMP signal that represents calcium activity without the signal drop typically seen during CO_2_ exposure, which is caused by the decrease in pH (Fig. 10d). The corrected calcium dynamics show a gradual increase during CO_2_ exposure and overshoots when CO_2_ exposure ends. These dynamics closely resemble the drift and overshoot observed during optogenetic activation (see Fig. 9b, FB, bottom row and blue line).

#### 2.6 Controller glia model during wakefulness and sleep

We modeled glia (referring to both ensheathing glia and astrocytes) as a controller that tries to maintains an optimal pH for neuronal activity, given by *H^s^*, assumed to be *-log*(*H^s^*)= 7. The model includes two compartments: a neuronal or neuropil compartment, where protons are produced at rate *r_N_*, and a glial compartment, where protons diffuse at a rate *d_H_* and are converted to CO_2_ through an equilibrium constant *k_eq_* via carbonic anhydrase at rate *k*. The model schematic is illustrated in Fig. 10a and is governed by the following system of differential equations:

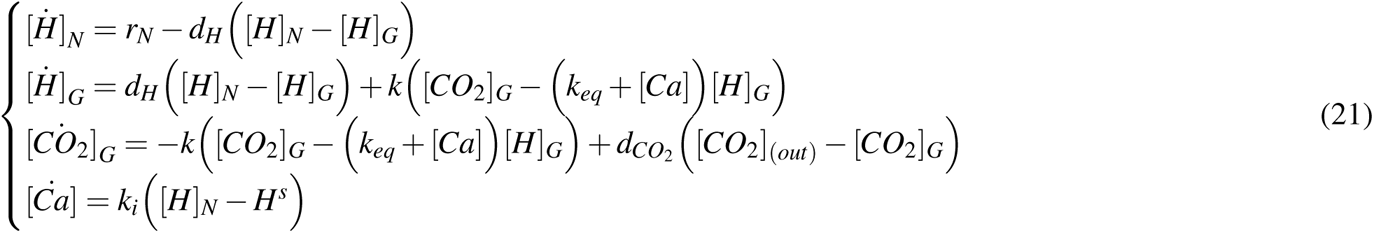

where [*H*]*_N_* is the concentration of protons in the neuronal compartment, [*H*]*_G_* is the concentration of protons in the glial compartment, and [*CO*_2_]*_G_* is the concentration of CO_2_ in the glial compartment. CO_2_ diffuses out of glia at a rate *d_CO_*_2_ through the tracheal system, where the external concentration is assumed to be [*CO*_2_]_(_*_out_*_)_ = 0. Calcium in glia [*Ca*] serves as a controller, modulating the proton-CO_2_ equilibrium to maintain the neuronal proton concentration setpoint *H^s^*. The response of the controller to pH deviations is defined by the integral gain constant *k_i_*.

**Table S7.** Parameter values for the model simulations in Fig. 10b,e, and f.

| $H^s$<br>(a.u.) | $r_N$<br>(seconds<sup>-1</sup>) | $d_H$<br>(seconds<sup>-1</sup>) | $k$<br>(seconds<sup>-1</sup>) | $k_{eq}$<br>(a.u.) | $d_{CO_2}$<br>(seconds<sup>-1</sup>) | $[CO_2]_{(out)}$<br>(a.u.) | $k_i$<br>(seconds<sup>-1</sup>) |
| --- | --- | --- | --- | --- | --- | --- | --- |
| 1e-7 | 3.3e-08 | 0.5 | 1e7 | 0.1 | 10 | 0 | 5e4 |

**Table S8.** Values for [*CO*_2_]_(_*_out_*_)_ during the simulated CO_2_ exposure.

| Simulated 10% CO<sub>2</sub> | Simulated 50% CO<sub>2</sub> | Simulated 100% CO<sub>2</sub> |
| --- | --- | --- |
| $[CO_2]_{(out)}$ | $[CO_2]_{(out)}$ | $[CO_2]_{(out)}$ |
| 1e-7 | 5e-7 | 1e-6 |

Note that we did not include a potential CO_2_ source from neuronal metabolic activity in our model. Although a more complex model incorporating CO_2_ concentration in neurons is possible, we opted for a simpler approach with fewer differential equations.

The model proposes that glia regulate the equilibrium between CO_2_ and protons, based on previous findings that astrocytes can influence this balance through pH bicarbonate buffers and ion channels^102,103^ . The model simplifies the various ion channels and buffering mechanisms, offering an abstract representation of glial regulation of pH by adjusting the equilibrium between CO_2_ and protons. This conceptualization focuses on glial modulation of the equilibrium without detailing every contributing ion channel and buffer.

We used the model to simulate calcium activity profiles in ensheathing glia and astrocytes during sleep and wake cycles (Fig. 10b). Model parameters are listed in Table S7. The baseline proton production rate in the neuronal compartment, *r_N_*, was calculated from the rest of parameters (equation 22) to obtain a pH of exactly 7 in the neuronal compartment ([*H*]*_N_* = *H^s^*). This baseline rate is assumed to occur during lower metabolic states, such as sleep in the fly, when the glial controller does not need to counteract any deviations from the pH setpoint.

During wakefulness however, higher neural activity leads to CO_2_ production and ATP hydrolysis and corresponding acidification^41,42,109,110^. We therefore assume that the proton production rate is increased by 1.5 times during wakefulness (Fig. 10b)

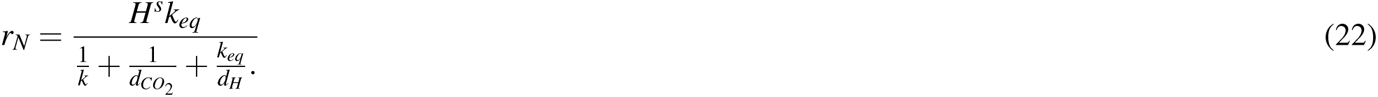

During wakefulness, the pH in the neuronal compartment decreases and diffuses into the glial compartment. Calcium senses this deviation from the setpoint and begins to rise. As calcium levels increase, it shifts the equilibrium between CO_2_ and protons, as indicated by the term *k_eq_* + [*Ca*] in equation (21). The chosen value of the equilibrium constant *k_eq_* reflects the limited conversion of protons to CO_2_, which also requires bicarbonate ions.. This calcium driven shift leads to a reduction in proton concentration and an increase in pH within the glial compartment during wakefulness (Fig. 10b, fourth row). The alkalinization of glia during neural activity aligns with previous findings in mammalian astrocytes^144,145^. An alternative model could result in glial acidification if calcium aimed to maintain the pH setpoint in the glial compartment rather than the neuronal one (model not shown). The converted CO_2_ diffuses through the trachea, with *d_CO_*_2_ chosen to reflect CO_2_ diffusion in air. Additionally, the value of the integral gain of the calcium controller, *k_i_* is manually adjusted to align the model dynamics with the observed data.

When calcium levels reach a high threshold, glial cells induce sleep by inhibiting neural activity, thus reducing proton production. In the model, the proton production rate is reset to the value of *r_N_* in Table S7. This transition from wakefulness to sleep could helps prevent excessive energy expenditure on the glial controller (not modeled here), facilitating the replenishment of energy stores and restoration of pH bicarbonate buffers and other substrates. During sleep, calcium levels gradually decrease while pH returns to the setpoint (Fig. 10b).

#### 2.7 Controller glia model during CO_2_ exposure

The previous model can explain the calcium slow drift and overshoot during and after CO_2_ exposure and optogenetics activation.

To simulate CO_2_ exposure in the model, we assume that the external CO_2_ concentration, [*CO*_2_]_(_*_out_*_)_ is nonzero. The values of [*CO*_2_]_(_*_out_*_)_ during 10%, 50% and 100% of CO_2_ exposure are shown in table S8. Figure 10d shows the response of the model to the three CO_2_ concentrations, overlaid with recorded data from pHluorinSE in both neurons and ensheathing glia, as well as the corrected jGCaMP8 data from ensheathing glia (see section 2.5 and Supplementary Fig. S55). When [*CO*_2_]_(_*_out_*_)_ is present, it diffuses into the glial compartment, converting to protons and causing a decrease in pH within both the glial and neuronal compartments. To account for the observed slow drift of calcium and the overshoot following 50% and 100% CO_2_ exposure, we propose that the significant drop in glial pH partially inhibits the calcium controller. Specifically, we suggest that the integral gain parameter, *k_i_* is influenced by the pH levels in the glial compartment (Fig. 10c). As pH decreases, *k_i_* is assumed to decrease as well, indicating a reduction in the effectiveness of the controller under acidic conditions. This partial inhibition leads to a slow response to pH deviations, resulting in a gradual rise in calcium and a modest increase in pH during 50% and 100% CO_2_ exposure (Fig. 10d).

At the end of 50% and 100% CO_2_ exposure, respectively, the external CO_2_ concentration, [*CO*_2_]_(_*_out_*_)_ is set to zero, leading to a decrease in proton concentration in the glial compartment. As the pH returns to physiological levels, the partial inhibition of the calcium controller is lifted, restoring *k_i_*to its normal value. For a brief period, the pH in the neuronal compartment is still low (as the protons have not diffused yet), causing the calcium controller to detect a significant deviation from the setpoint and respond strongly. This generates the observed overshoot dynamics after CO_2_ exposure (Fig. 10d), similarly to an impulse response in control systems.

The calcium overshoot leads to an increase in pH in both the neuronal and glial compartments. This elevated alkaline pH inhibits the controller, as indicated by the brief decrease in *k_i_* at the end of CO_2_ exposure (shown in Fig. 10d). Consequently, this inhibition results in a delayed response in calcium dynamics, as seen in the different overshoot patterns observed after 50% and 100% CO_2_ exposure. Once the glial pH returns to baseline, calcium levels also normalize. This model explains the drift and overshoot dynamics during CO_2_ exposure by suggesting that the glia controller is pH-dependent and partially inhibited outside its physiological range (For the biological rationale behind controller inhibition during low pH, see section 2.9.)

#### 2.8 Controller glia model during optogenetics activation

Similar dynamics are observed during optogenetic activation compared to CO_2_ exposure experiments. Since we express CSChrimson in glia, which acts as a proton channel^44^, we simulate the effects of optogenetic activation by adding an additional term, *r_G_*, into the glial compartment:

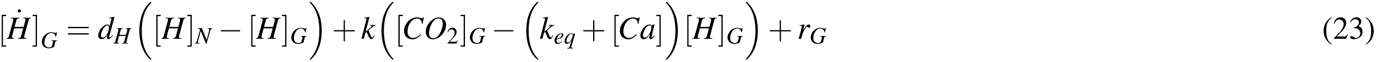

The term *r_G_* represents the rate at which protons enter the glial compartment during optogenetic activation, and it is set at *r_G_* = 90*r_N_* during this activation period. Because glial cells are negatively charged, the opening of proton channels allows protons to flow into the glial compartment through the electric gradient, resulting in acidification of this compartment (Fig. 10e).

The model simulation during optogenetic activation for 120 seconds is displayed in Fig. 10f, overlaid with the recorded calcium patterns from ensheathing glia in the FB. The observed slow drift and overshoot during and after optogenetic activation can be explained by the partial inhibition of the controller due to acidification, similar to the mechanisms described in the previous section regarding CO_2_ exposure.

#### 2.9 Potential mechanisms for biological implementation of the proposed model

The proposed model is consistent with carbonic anhydrase observed specifically in glia (Fig. 8d), which can also contribute to calcium regulation in some species^146^, and the close connection between trachea and glia facilitating gas exchange (Fig. 8). Further, neural activity leads to CO_2_ production and ATP hydrolysis and corresponding acidification^41,42,109,110^. In the proposed model, alkalinization of glia during neural activity (Fig. 10b, fourth row), is consistent with previous data in mammalian astrocytes^144,145^. However, a similar model with acidification in both compartments could similarly explain the data. The model is further consistent with homeostatic chemosensory function of glia described in mammals^85–90^. Additionally, glia are known to regulate intracellular and extracellular pH through membrane-bound transporters and ion channels, activated by calcium signaling^102,145^. Controller inhibition could for example result from inhibition of pH-regulatory ion channels under acidic conditions^41^.

**Supplementary Video S1.** Video for experiment of fly 1. Top: time during experiment. Middle, right: raw imaging data (60 frames or 1 second average in epochs of 1 minute). Rotating blue (during the day) or grey (during the night) arch represents stripe orientation in VR with respect to the fly (the fly’s head and abdomen are pointing toward the upper and lower axis of the video, respectively). Right: side view of the fly on the ball. Behavior classification is shown in white in the left corner. The bottom panel shows the velocity of the fly (first row), behavior classification (second row), and glia activity in EB (green) and FB (blue) over time. Grey areas in top and bottom rows indicate night (VR display is off). The time of extracted movies in the experiment are shown are indicated by the green bar.

**Supplementary Video S2.** Same as Supplementary Video S1 for fly 2.

**Supplementary Video S3.** Same as Supplementary Video S1 for fly 3.

**Supplementary Video S4.** Same as Supplementary Video S1 for fly 4.

**Supplementary Video S5.** Same as Supplementary Video S1 for fly 5.

**Supplementary Video S6.** Same as Supplementary Video S1 for fly 6.

**Supplementary Video S7.** Classification of fly behavior using the 3D CNN. The video shows different moments where the fly performs different behaviors.

**Supplementary Video S8.** Example of tracking freely walking flies in optogenetics setup under infrared illumination.

---

**Figure S1.**
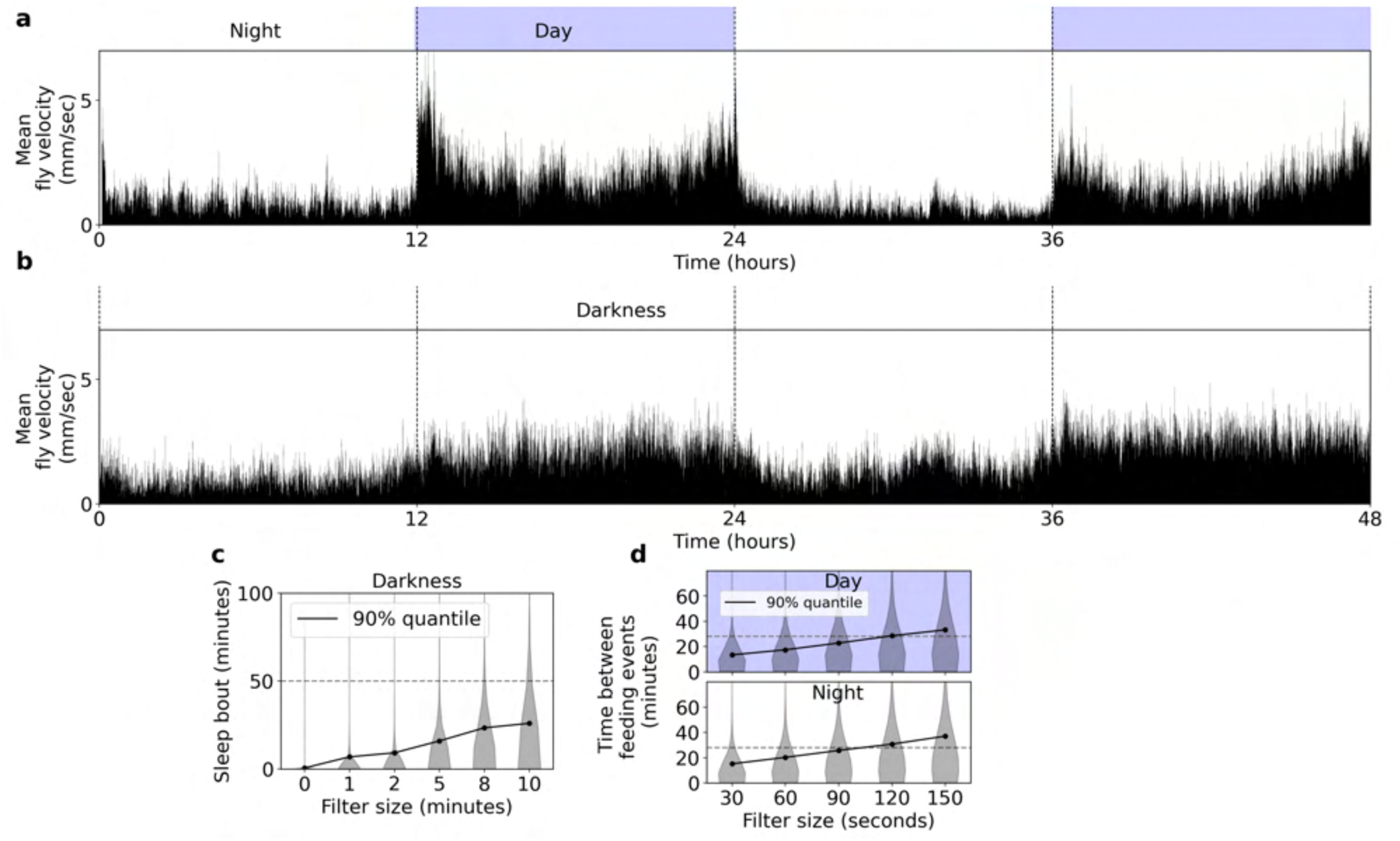
Behavioral activity of freely moving flies. **a** Mean fly velocity in 1 second bins over 48 hours with 12 hours of light (day) and 12 hours of darkness (night). **b** Same as in a, but in constant darkness. **c** Distribution of sleep bouts for a total of 15 flies in complete darkness (from b) as function of filter size. **d** Distribution of time between feeding events in freely moving flies as a function of filter size (see Methods).

**Figure S2.**
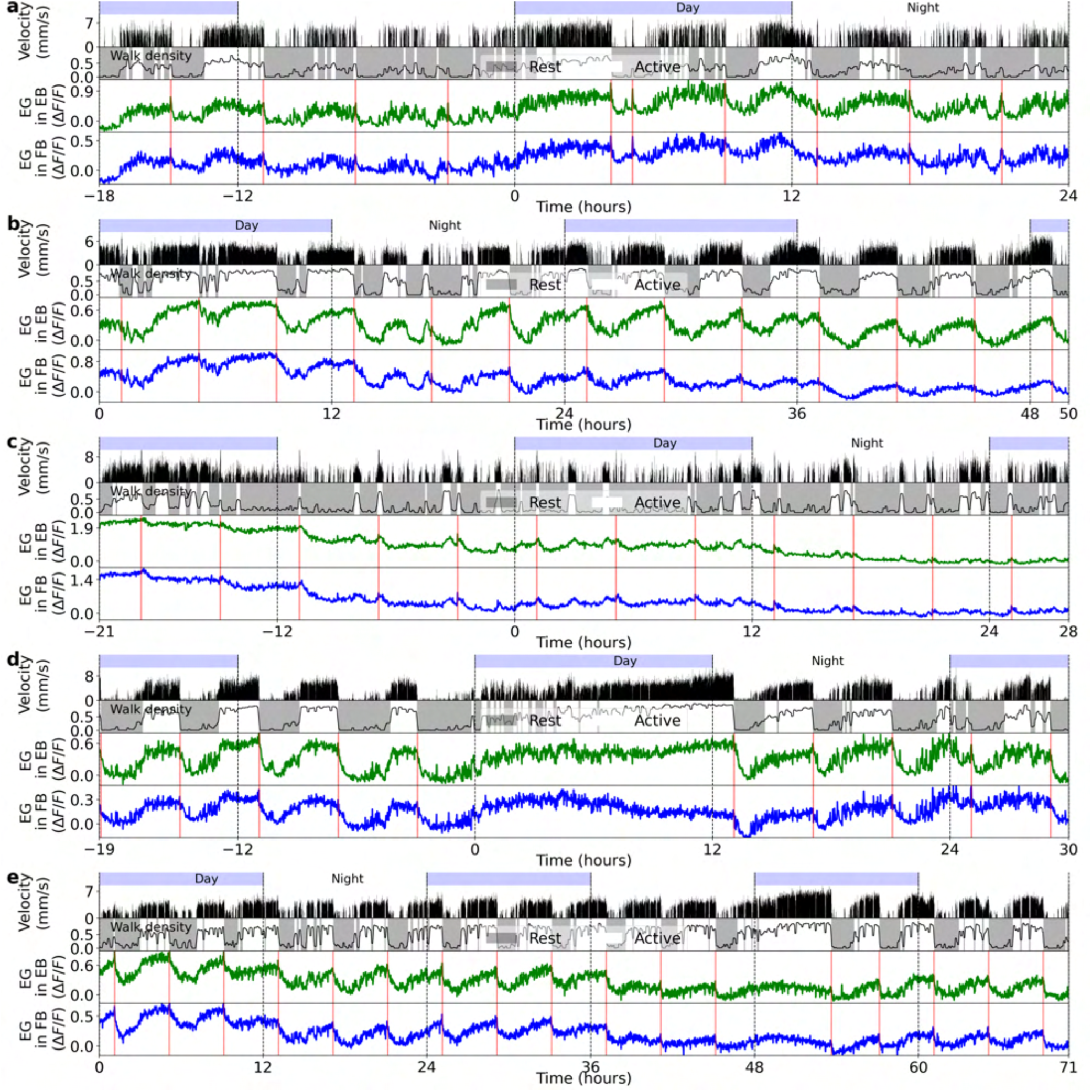
Regions of interests (ROIs) for recordings of ensheathing glia in LAL, MB, and midline, as well as of R5 and dFB neurons. **a** Schematic of EB, FB, and left and right LAL in the fly brain. **b** On the left, ensheathing glia in the central complex and LAL. On the right, ROI along the right (R) and left (L) LAL **c** On the left, *Drosophila* brain area where ensheathing glia in MB and midline were recorded (center). On the right, ROI along the MB (brown) and midline (dark orange). **d** On the left, R5 neurons labeled by 58H05-GAL4. On the right, ROI along the ring-shaped projections of R5 neurons for calculating fluorescence changes. **e** Same as in a, but for R5 neurons labeled by 88F06-GAL4. **f** Same as a and b, but for dFB neurons labeled by 23E10-GAL4.

**Figure S3.**
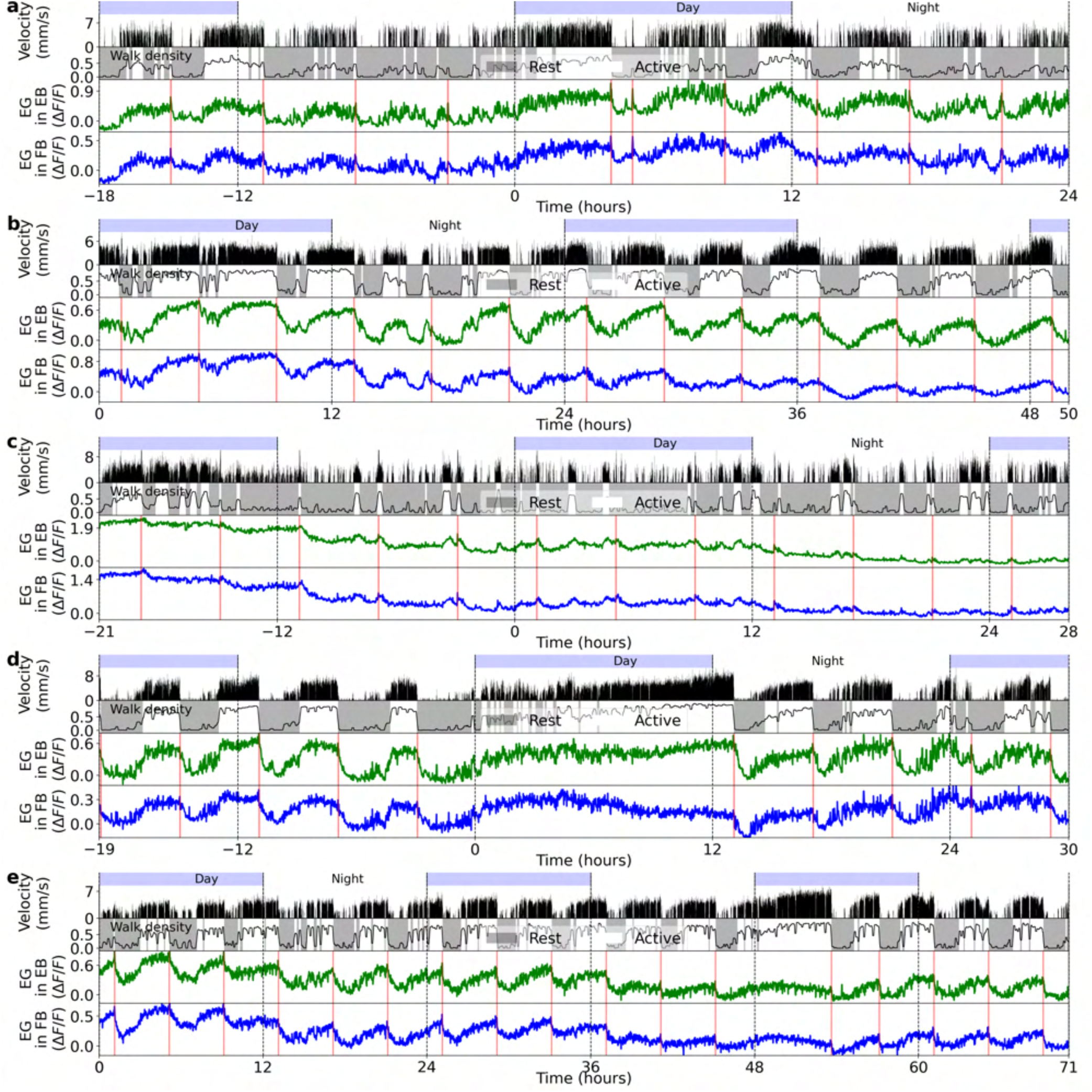
Five different recordings in ensheathing glia where flies are fed every 4 hours. **a** Top row: day and night cycle in VR. Second row: velocity of the fly in 1 second bins. Third row: walk density (see Methods) and rest (grey region) and active (white region) epochs. Fourth and fifth row: Calcium activity of ensheathing glia in the EB (green) and FB (blue), respectively. Thick lines indicate low-pass filter with a 0.1 hours cut-off period, while vertical red lines represent feeding events. **b-e** Same as a. Each panel shows different flies.

**Figure S4.**
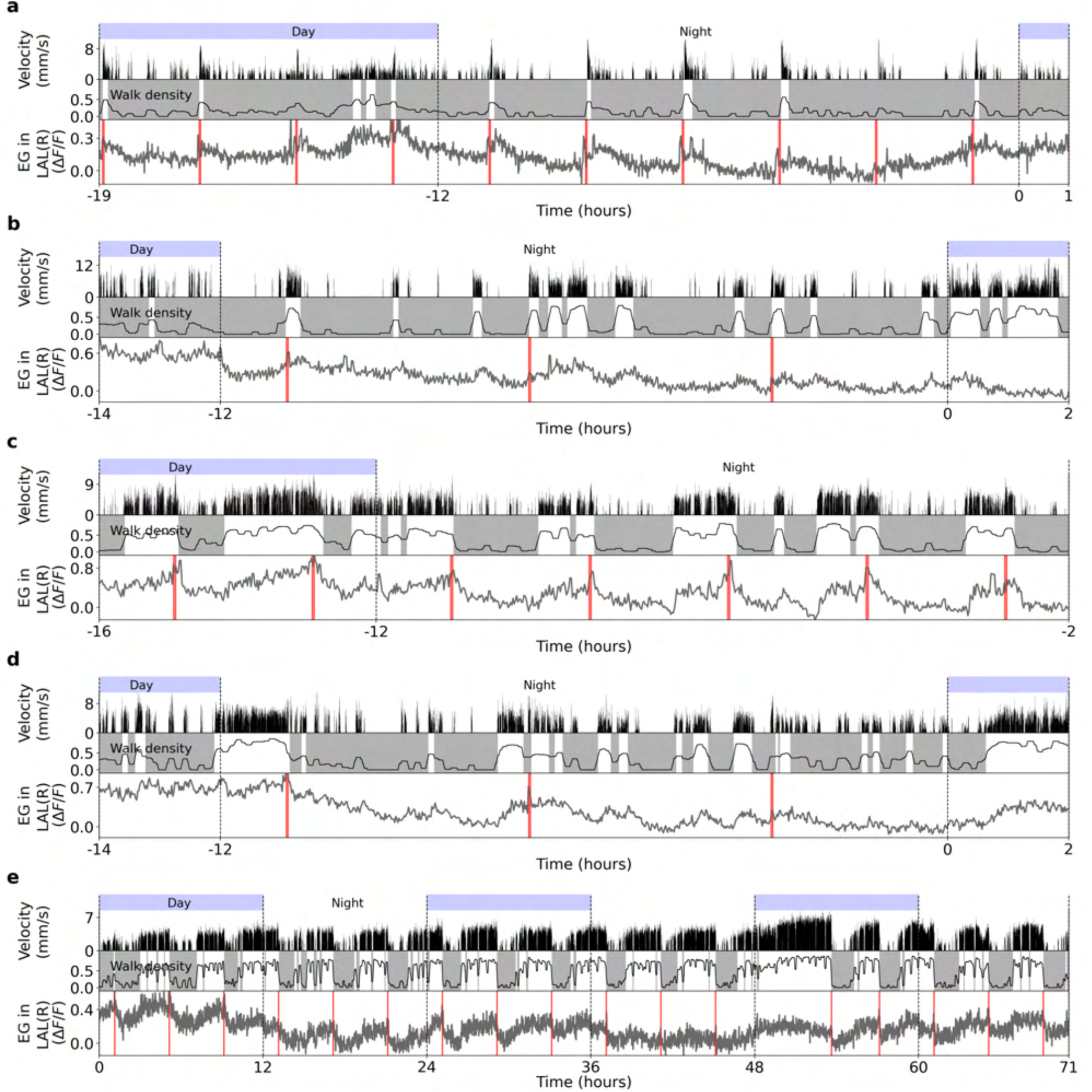
Ensheathing glia calcium activity in the LAL for 5 flies fed every 4 hours. **a** Top row: day and night cycle in VR. Second row: velocity of the fly in 1 second bins. Third row: walk density (see Methods) and rest (grey region) and active (white region) epochs. Fourth row: Calcium activity of ensheathing glia in the LAL (grey). Thick lines indicate a low-pass filter with 0.1 hour cut-off period, while vertical red lines represent feeding events. **b-e** Same as a. Each panel shows a different fly.

**Figure S5.**
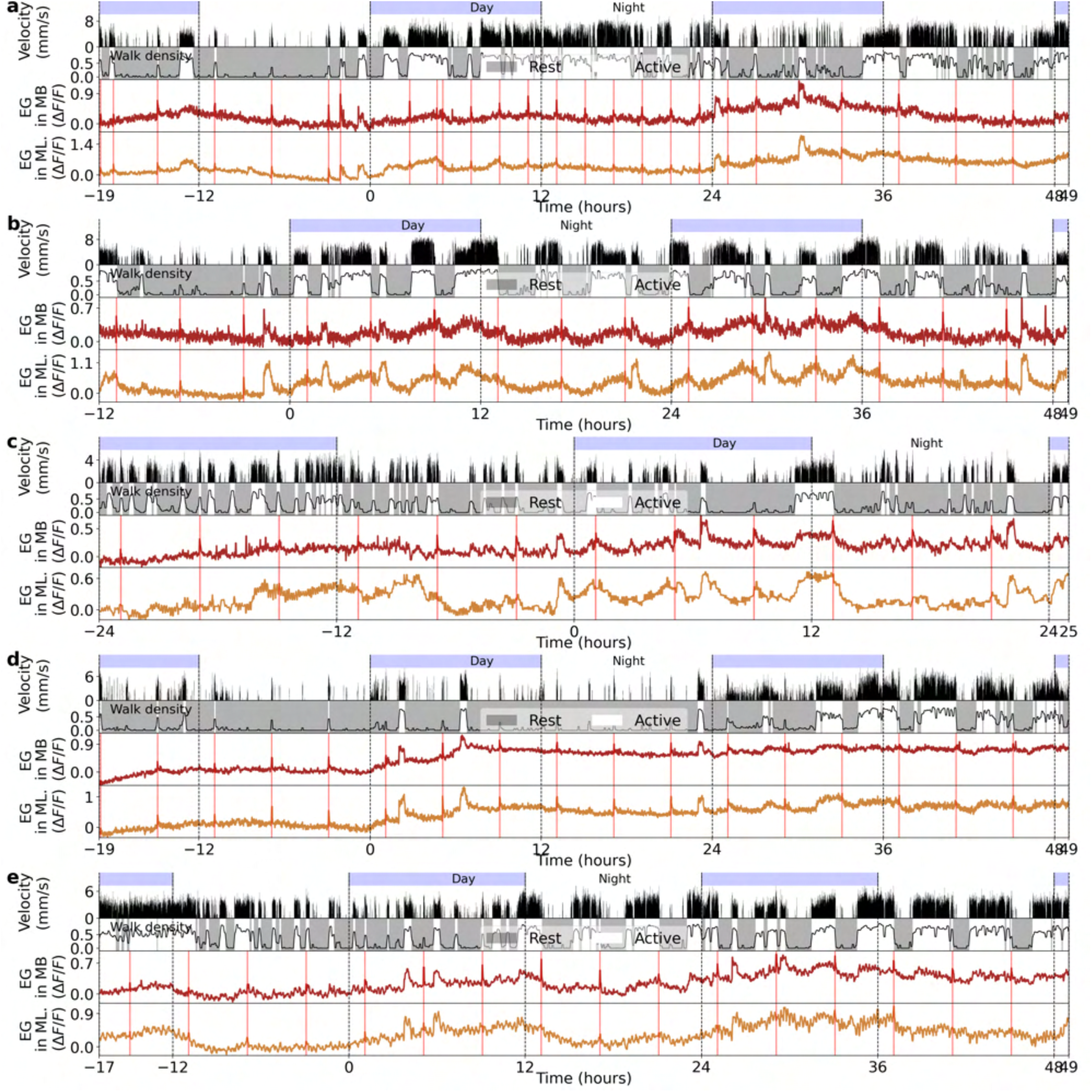
Ensheathing glia calcium activity in the MB and midline for 5 flies fed every 4 hours. **a** Top row: day and night cycle in VR. Second row: velocity of the fly in 1 second bins. Third row: walk density (see Methods) and rest (grey region) and active (white region) epochs. Fourth and fifth row: Calcium activity of ensheathing glia in the MB (brown) and midline (dark orange). Thick lines indicate a low-pass filter with 0.1 hour cut-off period, while vertical red lines represent feeding events. **b-e** Same as a. Each panel shows a different fly.

**Figure S6.**
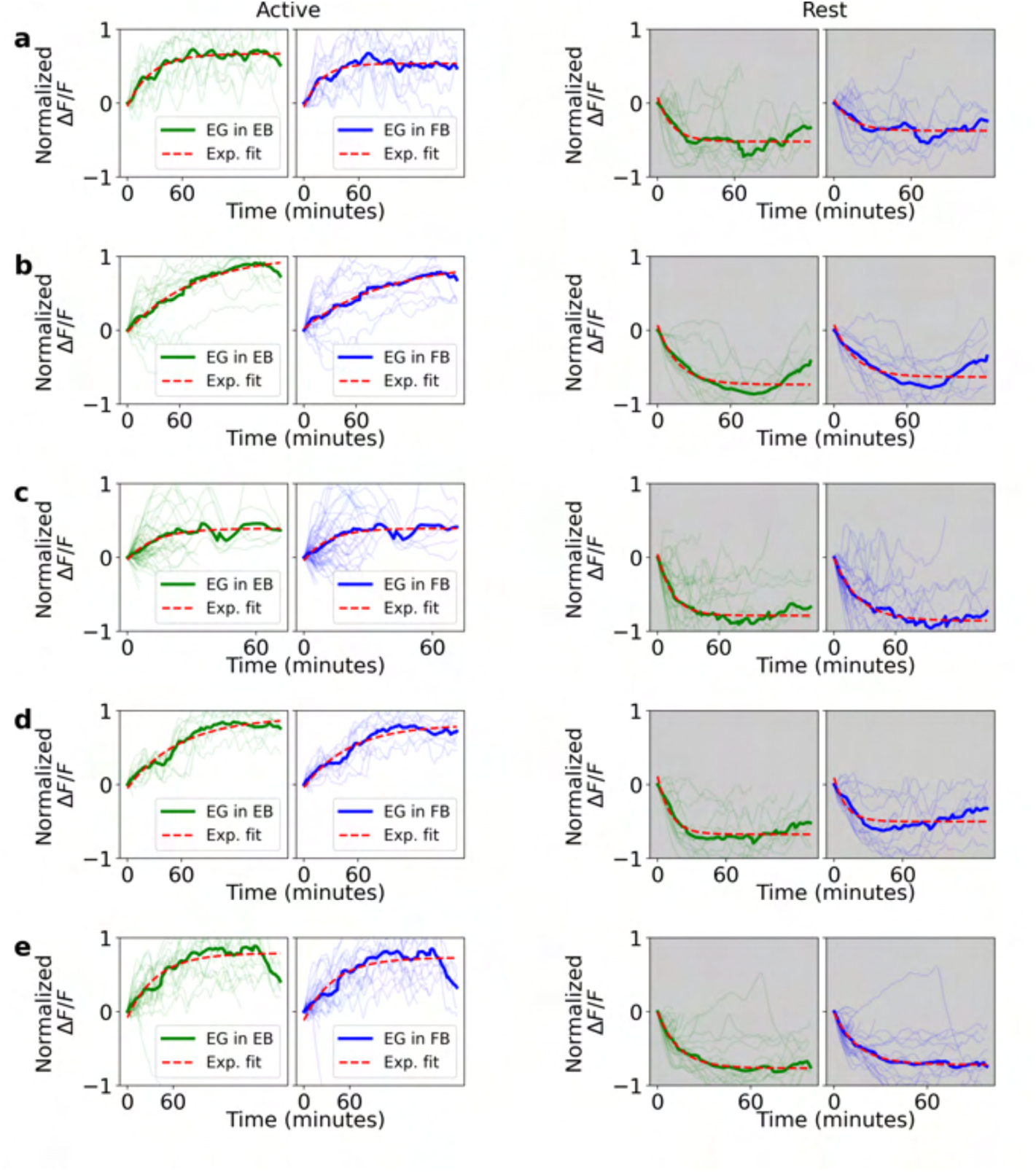
Normalized fluorescence traces during active and rest epochs for 5 flies. **a** Left side: single (thin lines) and average (thick lines) normalized fluorescence traces in the EB (green) and FB (blue) during active epochs. Red lines indicate exponential fit. Right side: same as left side, but during rest epochs. **b-e** Same as a. Each panel is from a different fly.

**Figure S7.**
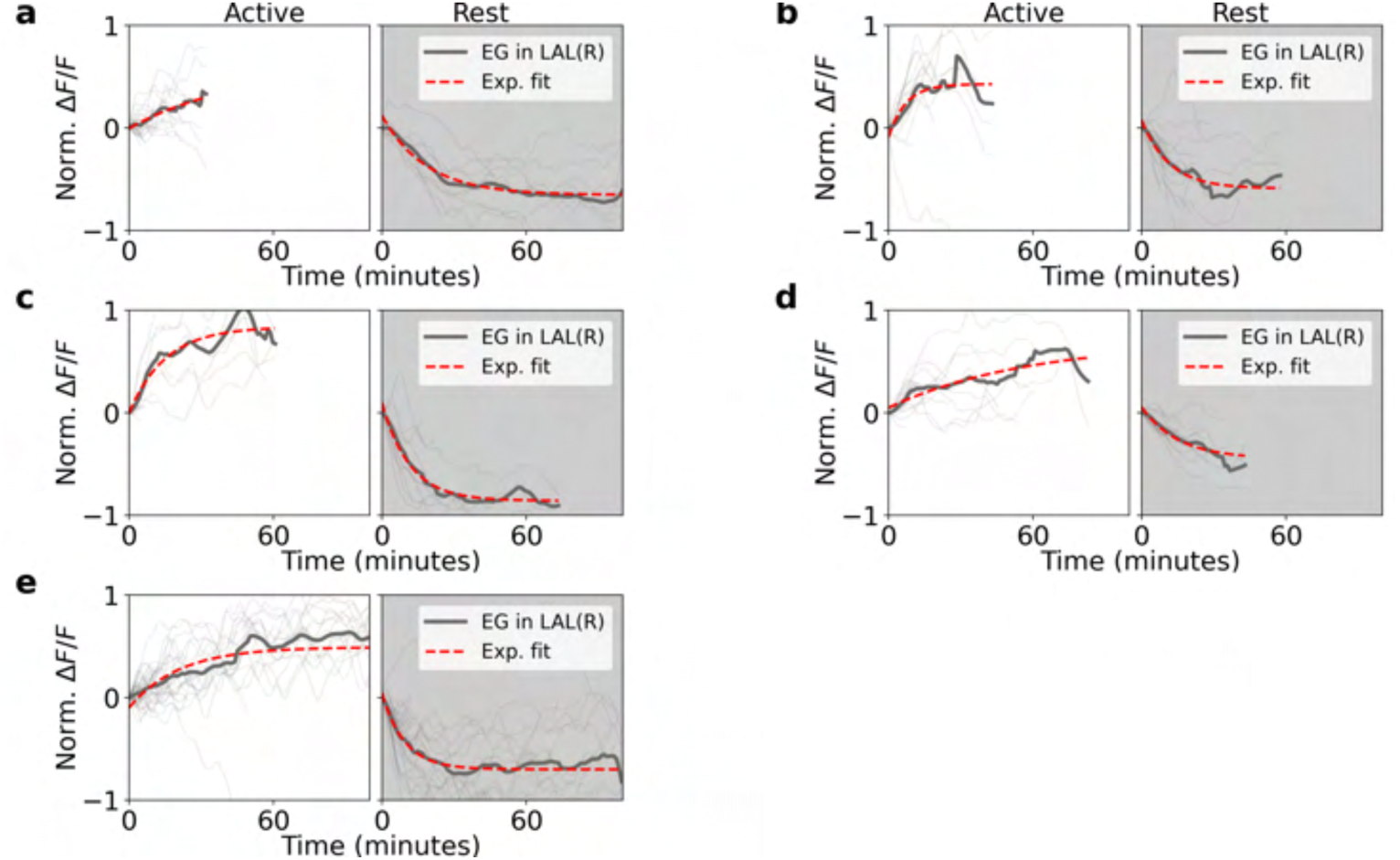
Normalized fluorescence traces during active and rest epochs for 5 flies. **a** Left side: single (thin lines) and average (thick lines) normalized fluorescence traces in the LAL (grey) during active epochs. Red lines indicate exponential fit. Right side: same as left side, but during rest epochs. **b-e** Same as a. Each panel is from a different fly.

**Figure S8.**
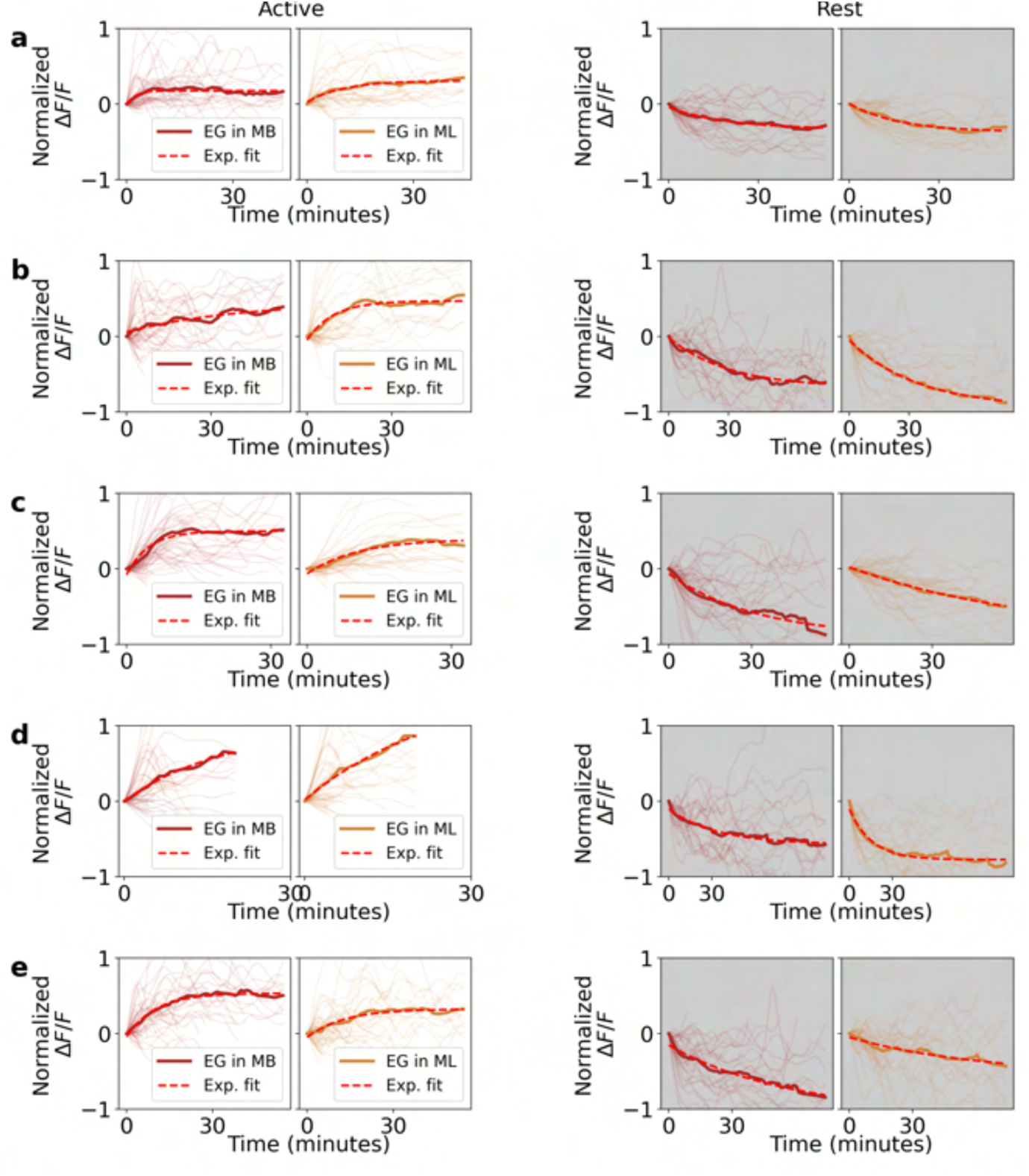
Normalized fluorescence traces during active and rest epochs for 5 flies. **a** Left side: single (thin lines) and average (thick lines) normalized fluorescence traces in the MB (brown) and midline (dark orange) during active epochs. Red lines indicate exponential fit. Right side: same as left side, but during rest epochs. **b-e** Same as a. Each panel is from a different fly.

**Figure S9.**
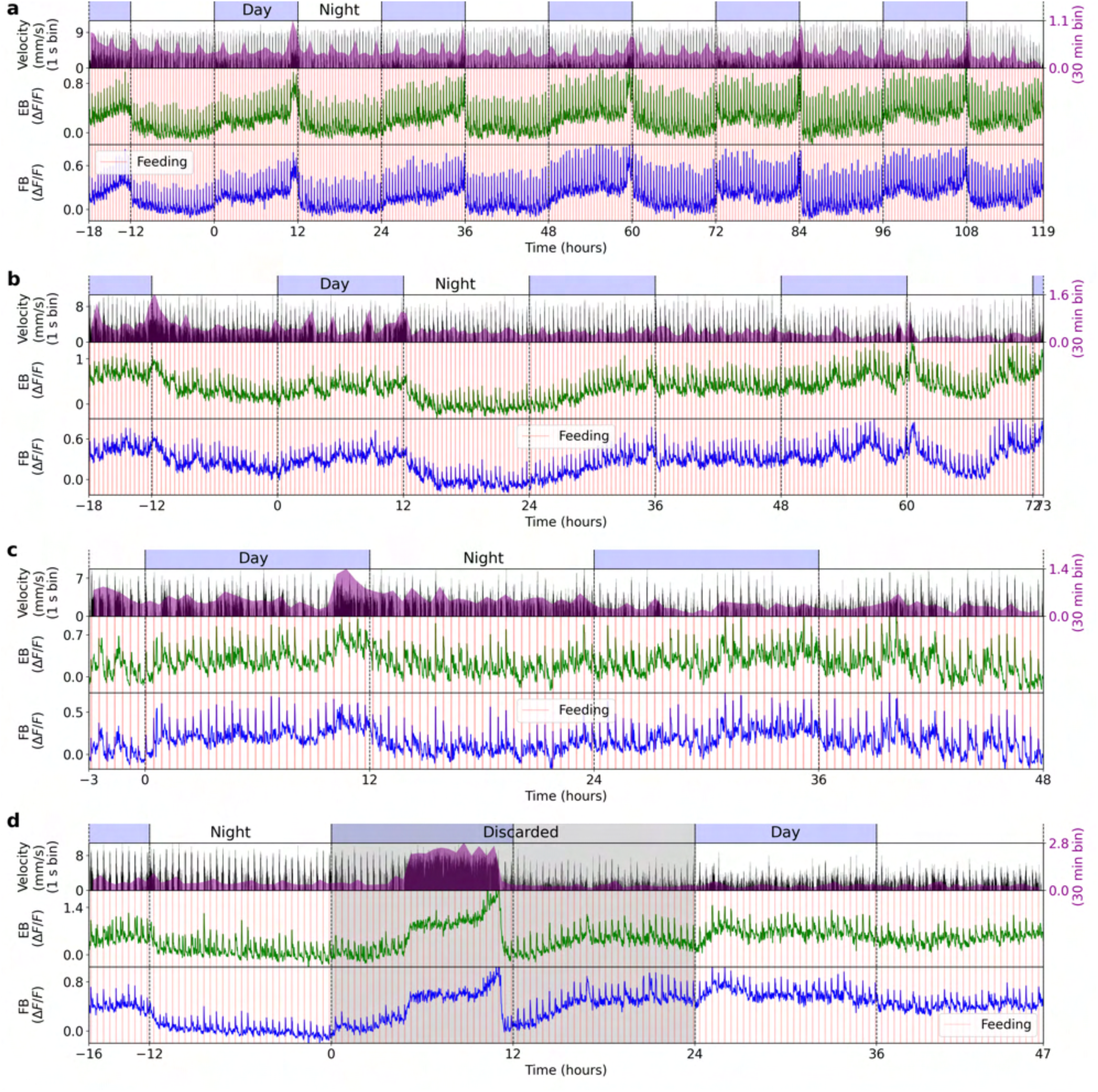
Four different recordings in ensheathing glia where flies are fed every 26 minutes. **a** Top row: day and night cycle in VR. Second row: in black, velocity of the fly in 1 second bins. Purple: velocity of the fly in 30 minutes bins. Third and fourth row: calcium activity of ensheathing glia in the EB (green) and FB (blue). Vertical red lines indicate feeding events. **b-d** Same as a. Each panel shows a different fly. In panel d, the grey area was discarded from the average in Fig. 7j, since there was an epoch of increased walking and calcium activity between hours 5 and 11. The reason for this increase during the recording is unknown but was likely due to an unexpected event during the recording such as a sudden increase in two-photon laser power or temperature. We also discarded the subsequent 12 hours of the experiment.

**Figure S10.**
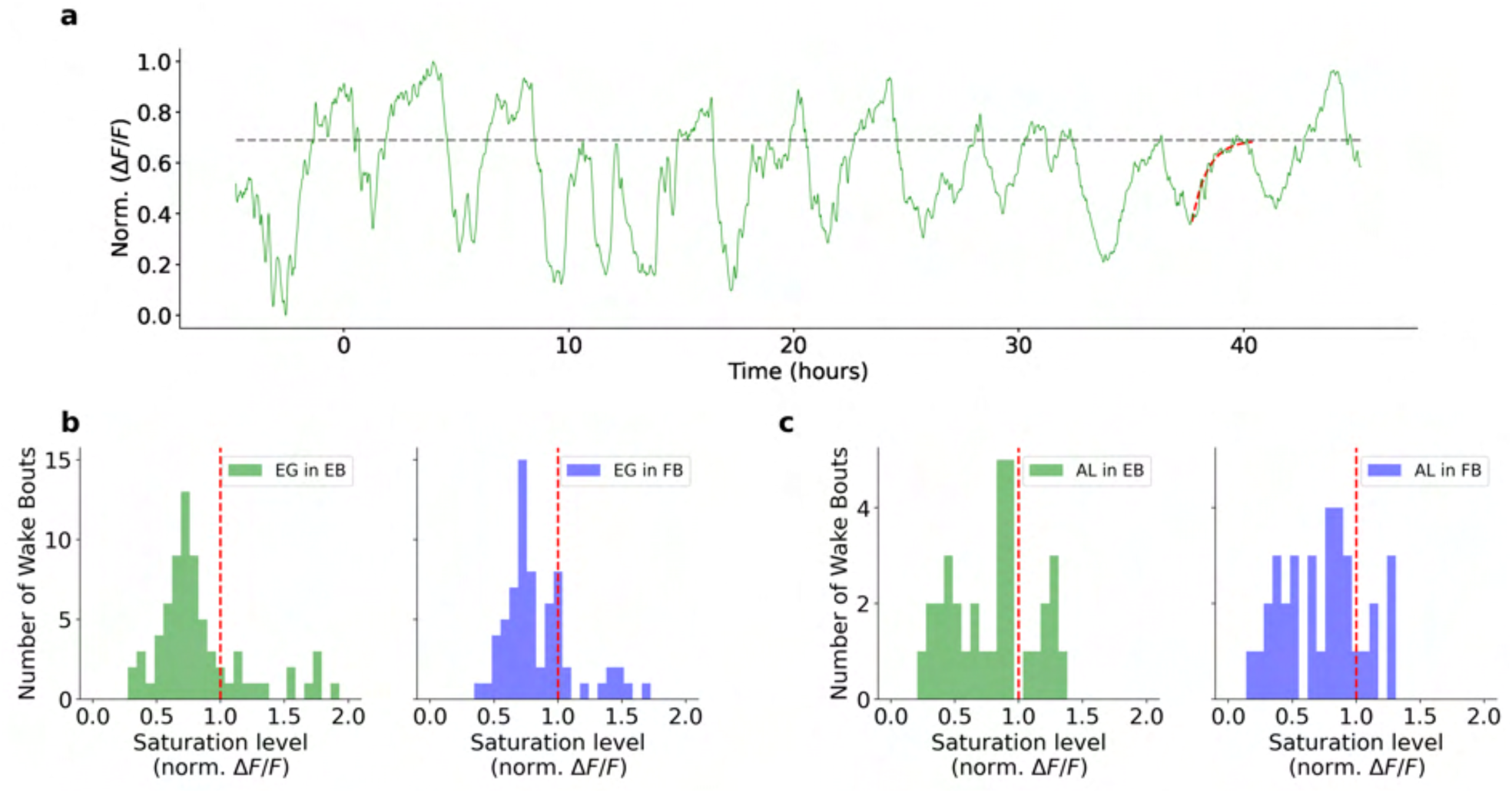
Fluorescence saturates due to calcium dynamics and not due to sensor limitations. **a** Filtered and normalized calcium activity recorded in the EB from ensheathing glia (green) over a long term experiment. This example shows an exponential fit (red) when the fly was active and the saturation level (grey) of the exponential. The saturation level was lower than the maximum value of fluorescence. **a** Distribution of saturation levels during active bouts in the EB (left) and FB (right) for ensheathing glia. The distributions reveal that some saturation levels are below 1 (shown as the red vertical line as the normalized maximum fluorescence), indicating that calcium activity reaches saturation in some active periods before the sensor reaches its maximum during the experiment. **c** Same as b but for astrocytes

**Figure S11.**
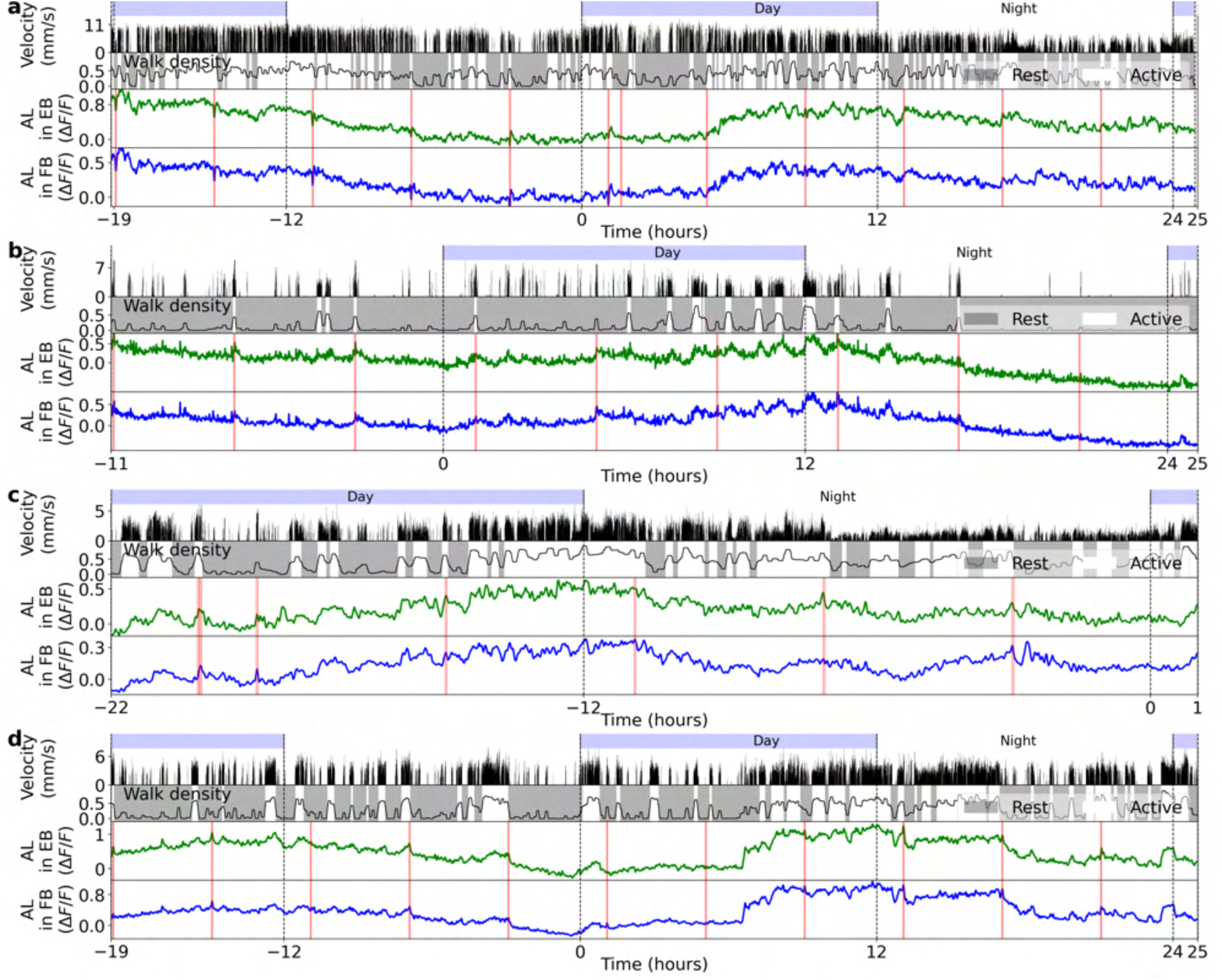
Astrocytes calcium activity in the EB and FB for 4 flies fed every 4 hours. **a** Top row: day and night cycle in VR. Second row: velocity of the fly in 1 second bins. Third row: walk density (see Methods) and rest (grey region) and active (white region) epochs. Fourth and fifth row: calcium activity of astrocytes in the EB (green) and FB (blue). Vertical red lines represent feeding events. **b-d** Same as a. Each panel shows a different fly.

**Figure S12.**
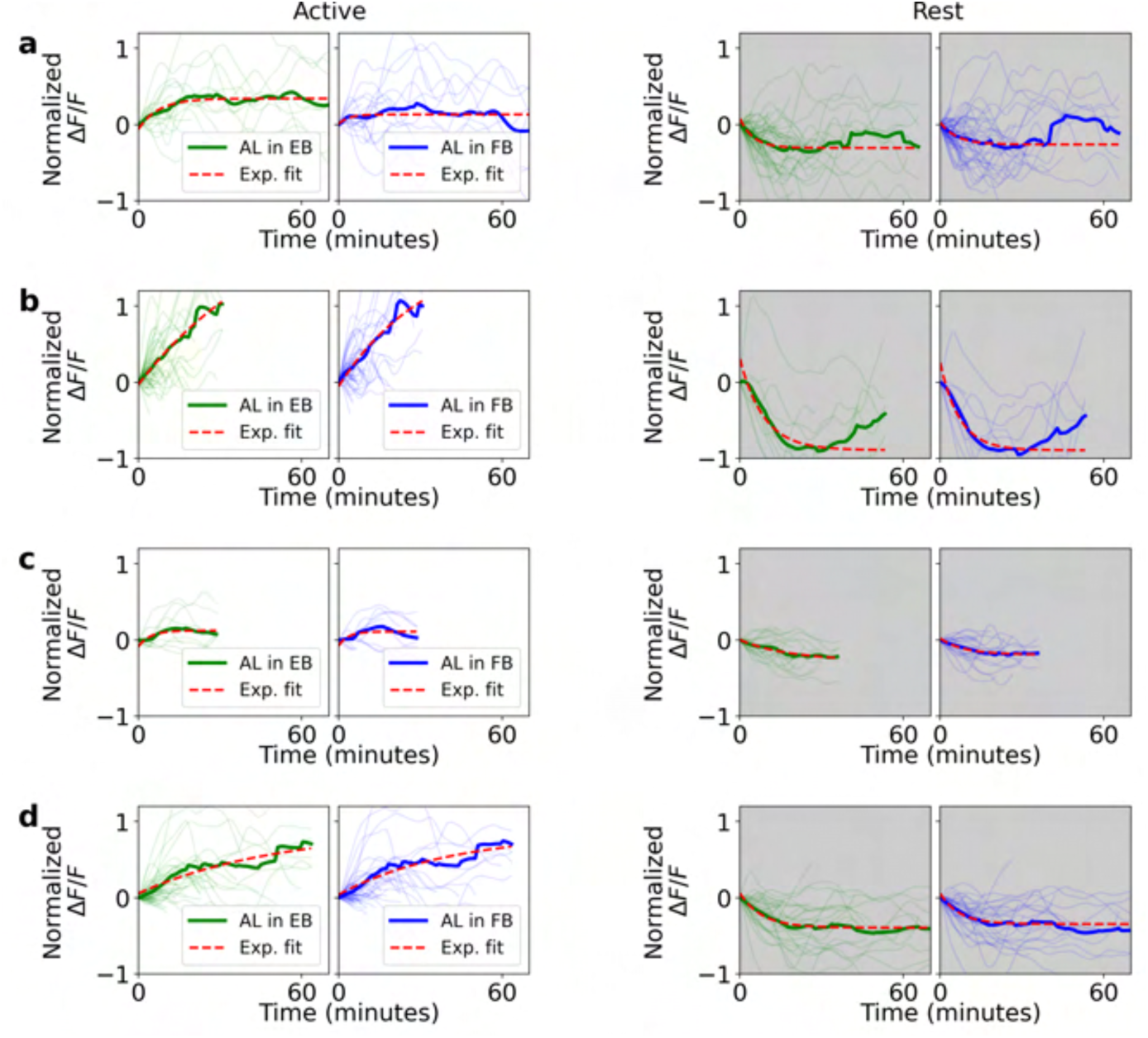
Normalized fluorescence traces in astrocytes during active and rest epochs for 4 flies. **a** Left side: single (thin lines) and average (thick lines) normalized fluorescence traces in the EB (green) and FB (blue) during active epochs. Red lines indicate exponential fit. Right side: same as left side, but during rest epochs. **b-d**Same as a. Each panel is from a different fly.

**Figure S13.**
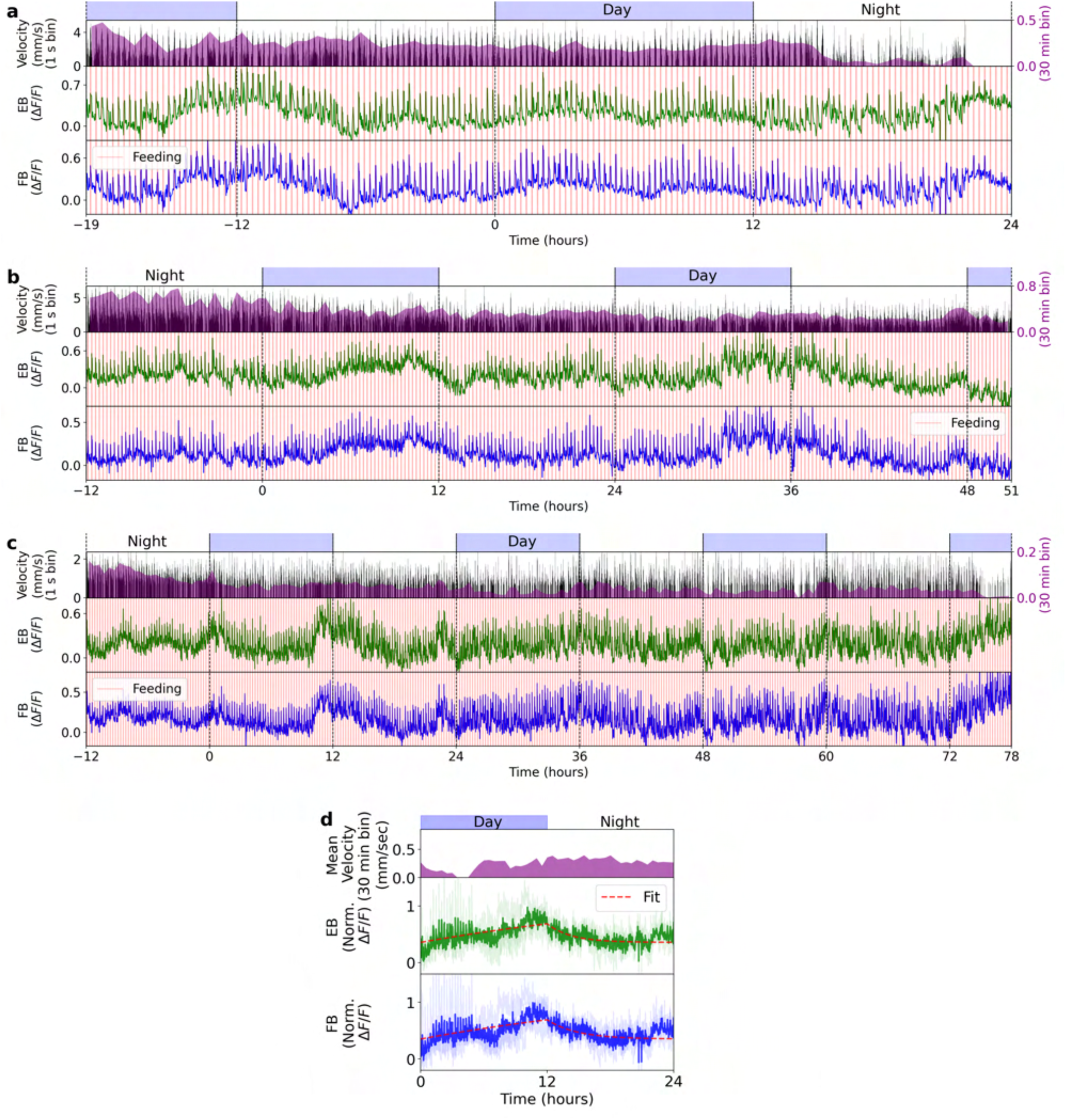
Three different long-term imaging recordings in ensheathing glia where flies are fed every 16 minutes. **a** Top row: day and night cycle in VR. Second row: in black, velocity of the fly in 1 second bins. Purple: velocity of the fly in 30 minutes bins. Third and fourth row: Calcium activity of ensheathing glia in the EB (green) and FB (blue), respectively. Vertical red lines indicate feeding events. **b, c** Same as a, but for different flies. **d** Average of previous recordings (in a, b, and c) with all trials aligned to one 24 hour period. Top row: day and night cycle. Second row: Mean velocity of all flies over 30 minutes bins. Third and fourth row: average glia activity in the EB (green) and FB (blue). Red lines indicate exponential fits during the day and night (see Methods).

**Figure S14.**
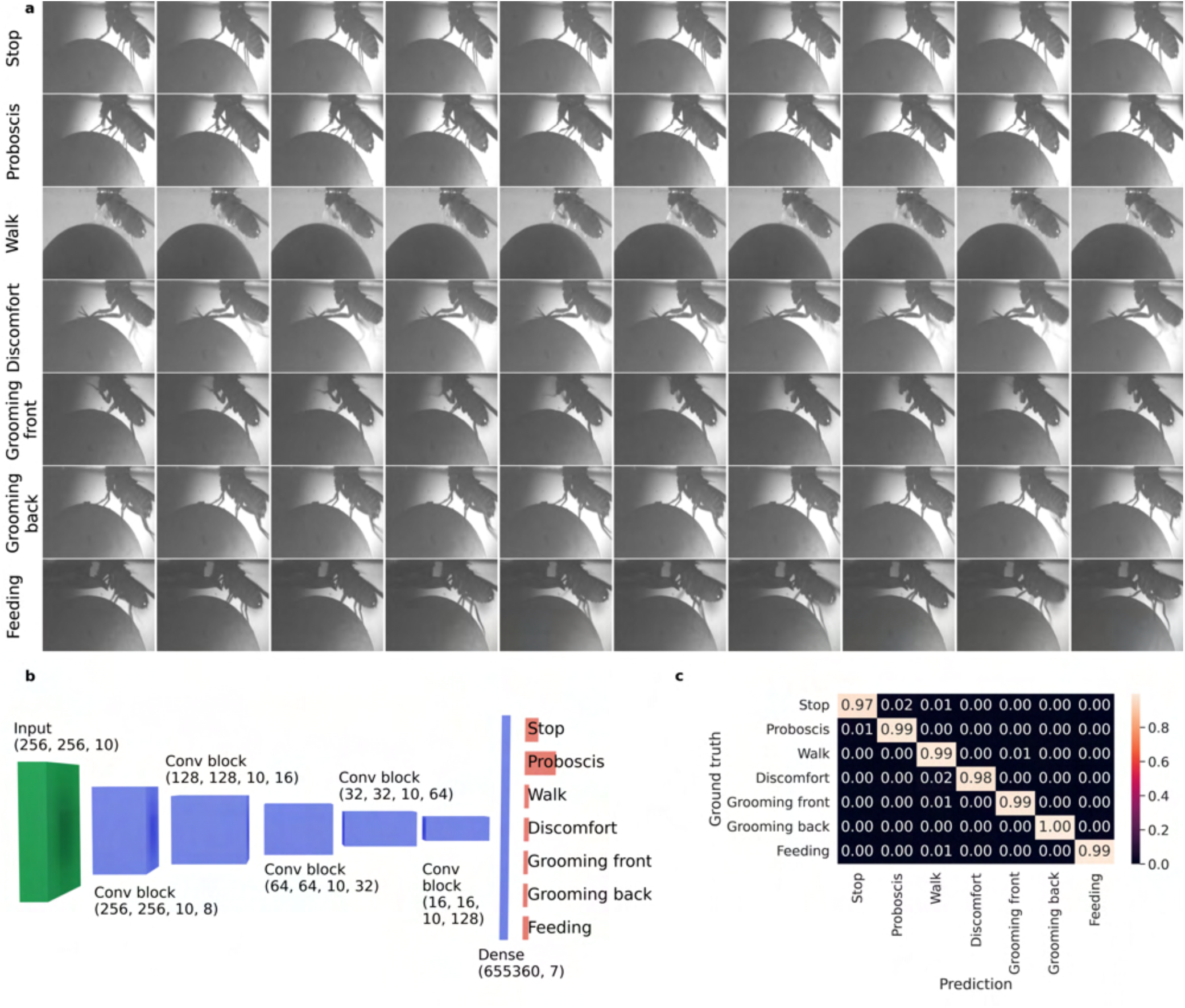
Classification of the behavior of the fly on the ball during long-term imaging. **a** A 3D CNN with 10 consecutive frames with a side view of the fly on the ball is used to classify 7 different behaviors (y-axis). Each row in the plot corresponds to 10 consecutive frames where the fly performs the labeled behavior. **b** Architecture of the 3D CNN. **c** Confusion matrix for each behavior for prediction of the 3D CNN (x-axis) on a test dataset that was manually labeled (ground truth along y-axis).

**Figure S15.**
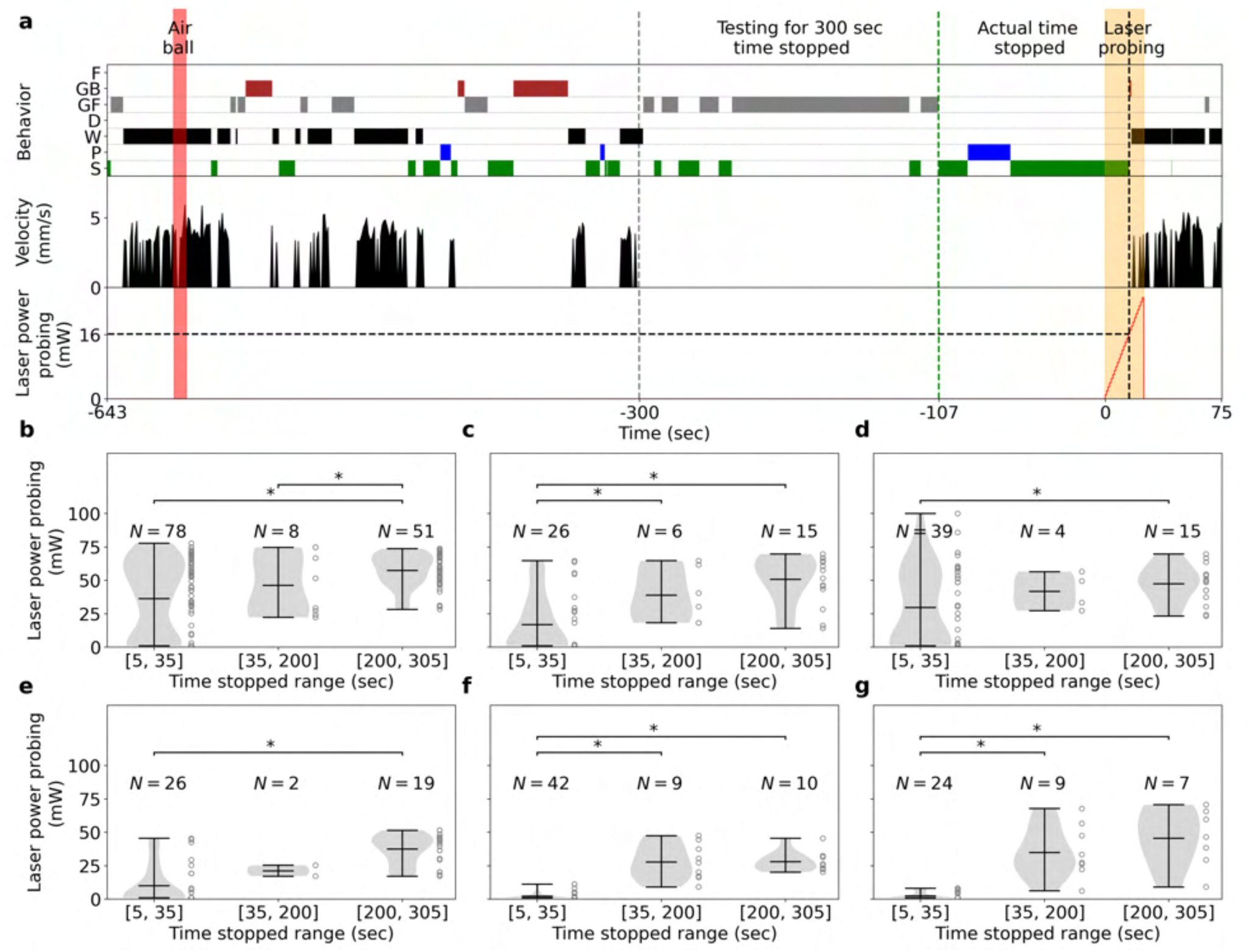
a. Example of trial protocol for testing arousal threshold after 300 seconds of immobility of a fly on the ball. First, the air ball is switched on and off (red area) to ensure the fly is awake. The fly stops walking for 300 seconds (fly velocity is zero in second row) and the laser (third row) starts probing the arousal threshold of the fly by increasing its power every 0.5 seconds (orange area). The laser is switched off when the fly starts walking again (velocity higher than 0). Post processing: behavior classification (first row) is used to compute the actual time fly was immobile, without grooming (107 seconds in this trial), as well as the actual laser power the fly reacts to by grooming, discomfort, or walking during laser probing (16 *mW* in this trial). **b,c,d,e,f,g** Distribution of laser powers at which fly wakes up for all trials for different durations of immobility for 6 flies. Asterisks indicate statistical significance using t-test (p-value lower than 0.05).

**Figure S16.**
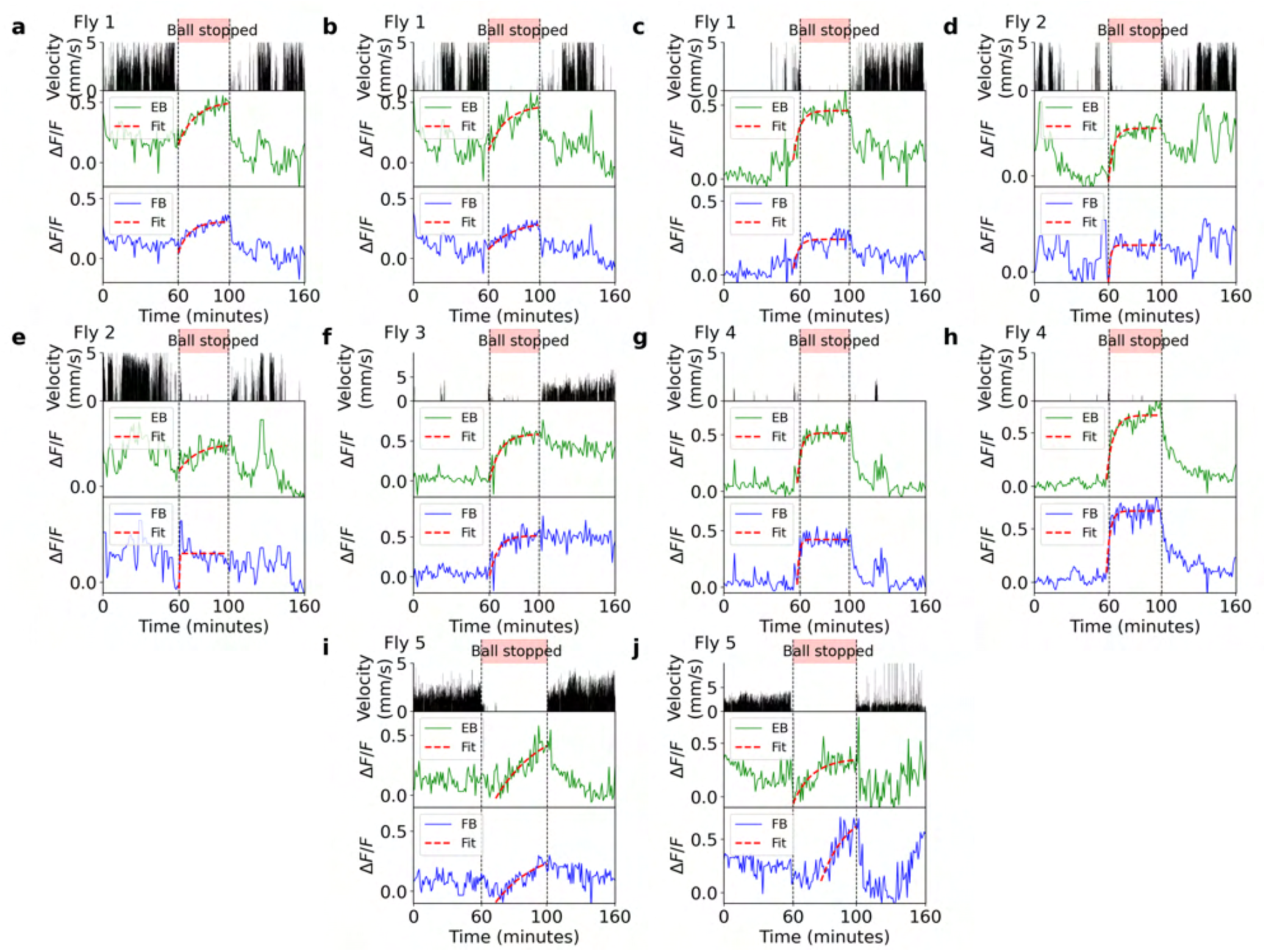
Trials where the ball was blocked during recordings in glia. **a** First row: time where the ball was stopped (red region). Second row: velocity of fly. Third and fourth row: Calcium activity of ensheathing glia in the EB (green) and FB (blue), respectively. Red lines are exponential fits. **b-j** Same as a. Each panel represents a different trial and the fly from which each trial was recorded is shown in the left top corner of each panel.

**Figure S17.**
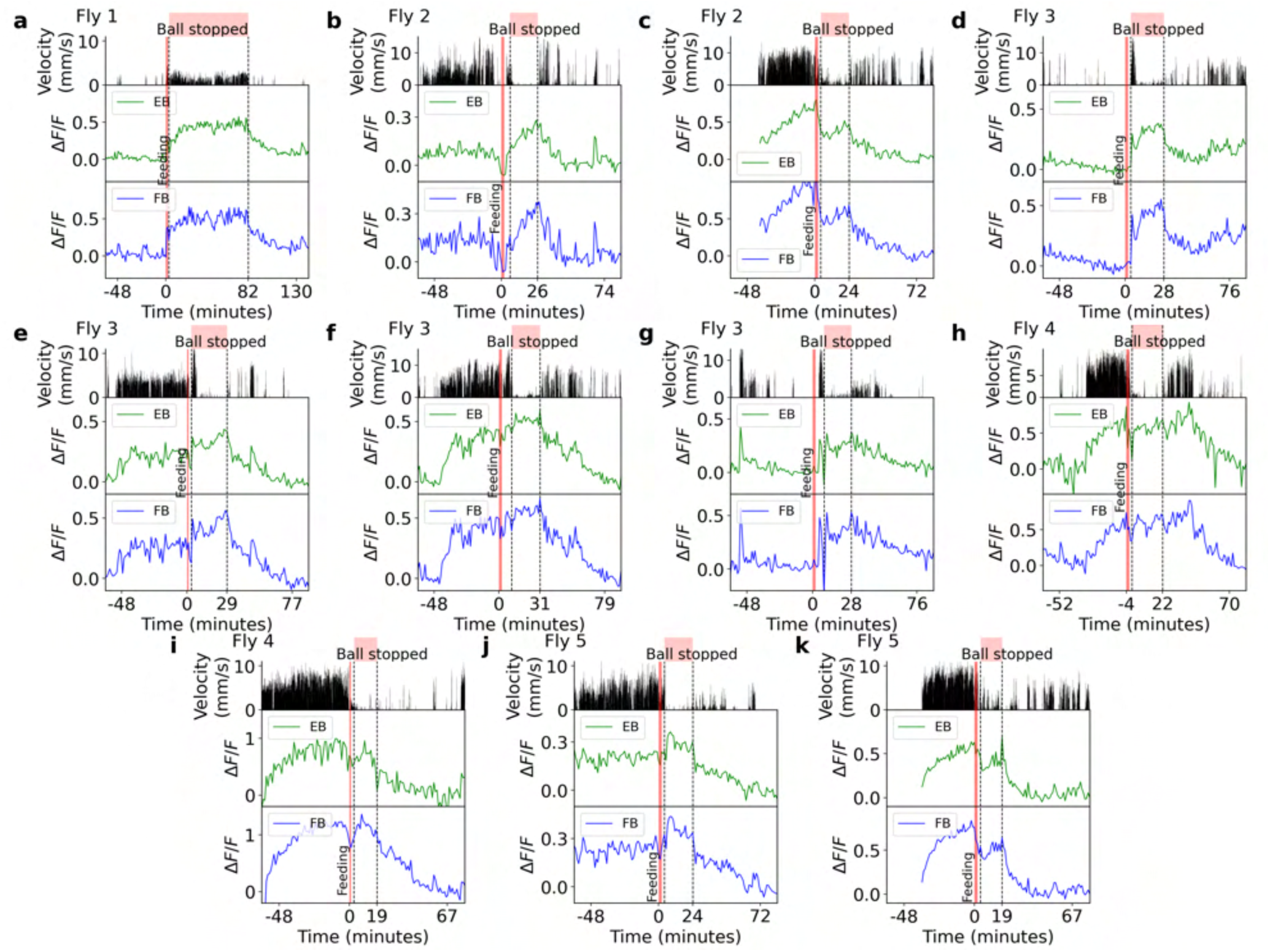
Trials where the ball was blocked after feeding while activity in glia was recorded. **a** First row: time where the ball was stopped (red region). Second row: velocity of the fly. Third and fourth row: Calcium activity of ensheathing glia in the EB (green) and FB (blue). The vertical red line indicates feeding. **b-k** Same as a. Each panel represents a different trial. The fly from which each trial was obtained is shown in the top left corner of each panel.

**Figure S18.**
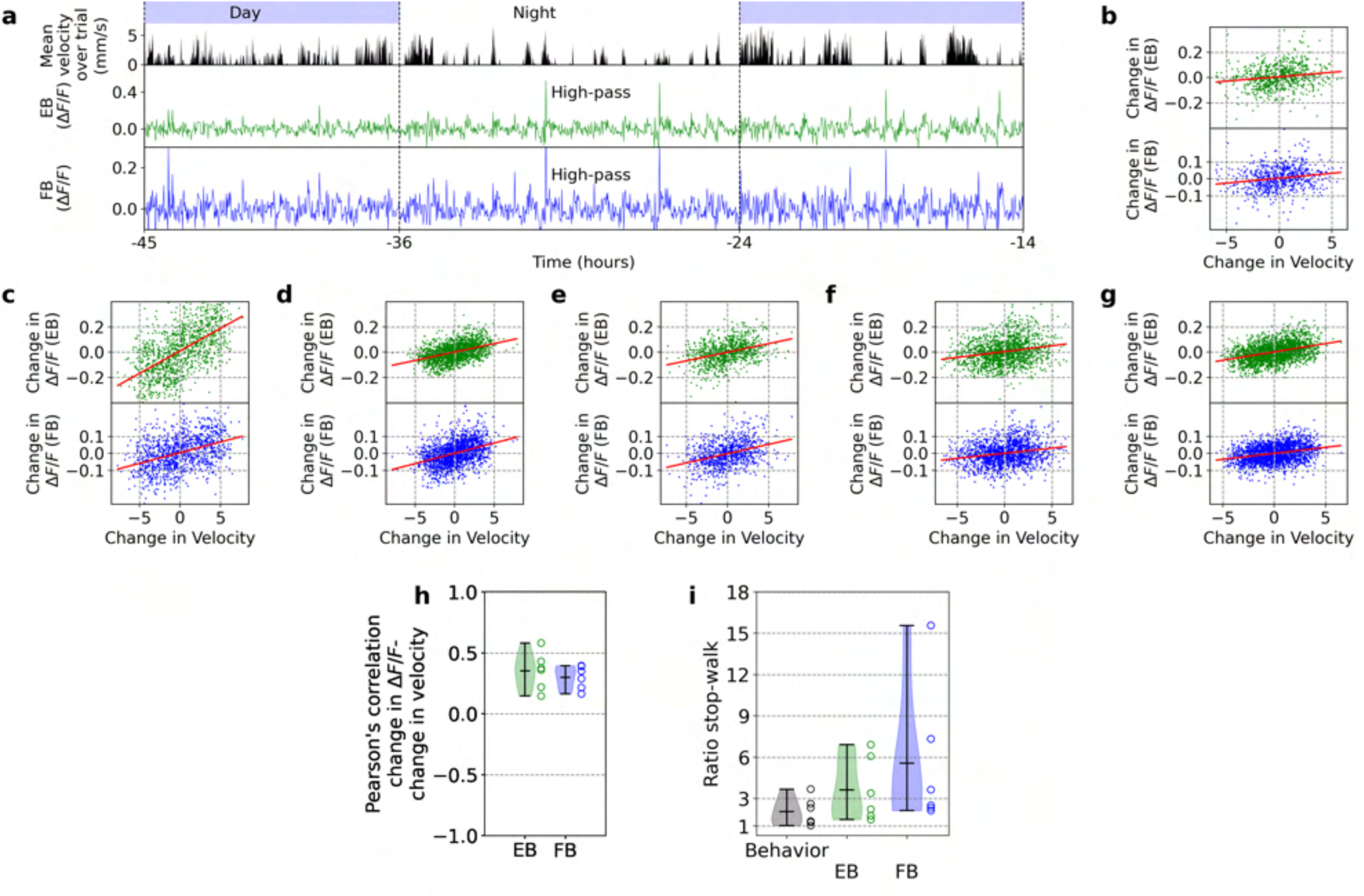
a. Top row shows day and night cycle in VR over time and second row shows mean velocity during recording epoch (1 second). Second and third row show fluorescence signals in the EB and FB, respectively. Fluorescence was filtered with a high-pass filter for periods higher than 0.5 hours. **b** Correlation between mean velocity over trial and high-pass filtered fluorescence. **c-g** Same as b for 5 more flies. **h** Ratios of time in ’stop’ and ’walk’ states (black) and ratio of time constants between ’stop’ and ’walk’ states in EB (green) and FB (blue) for each fly (*N* = 6). Ratios were not significantly different, p-values were above 0.05 using t-test. **i** Correlation coefficients as in e for *N* = 6 flies. All flies had p-values lower than 0.05 (see Table S2).

**Figure S19.**
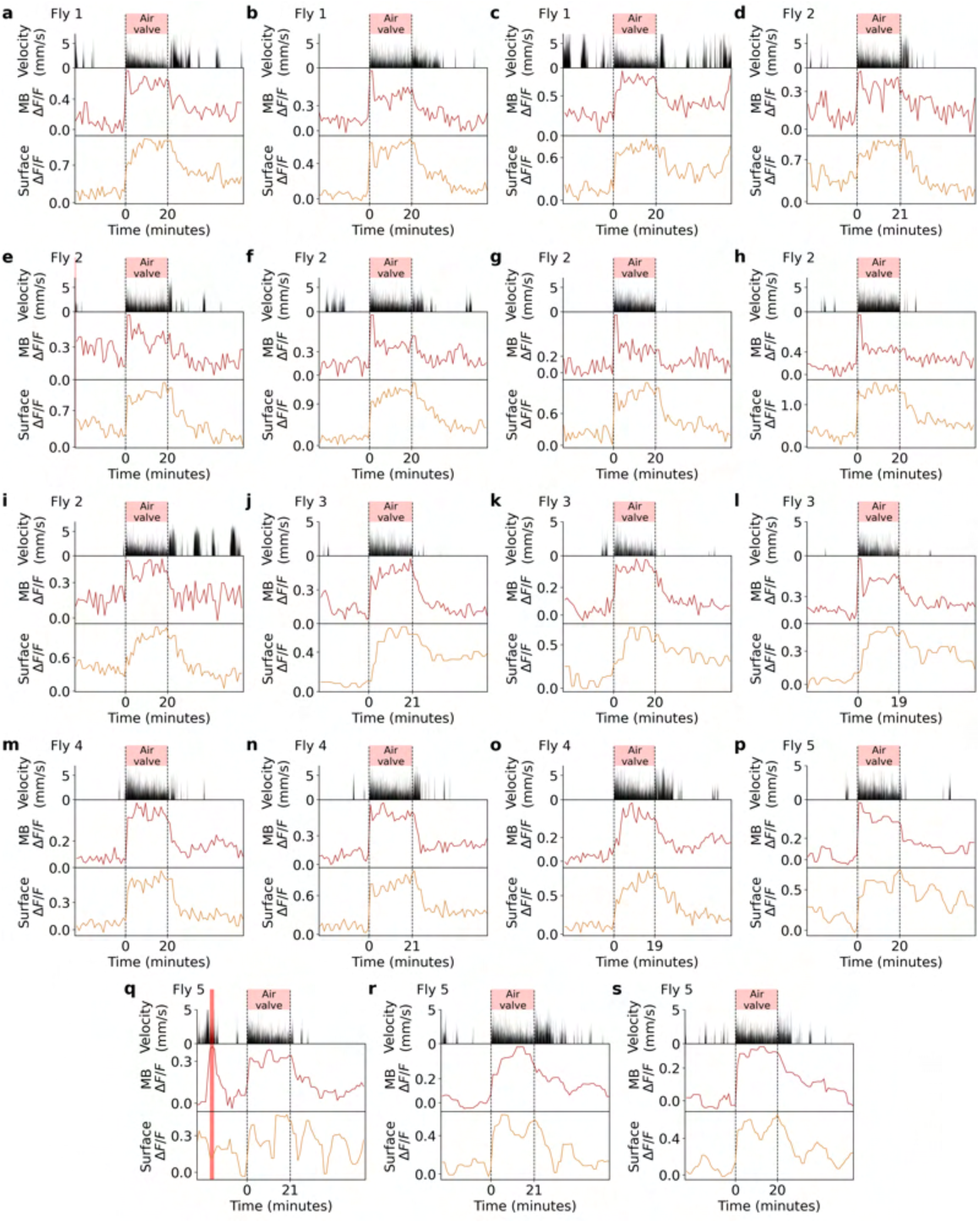
Trials where the air stream supporting the ball was intermittently opened and closed every second to promote walking behavior, while glia in the MB and midline was recorded. **a** First row: time where the air stream of the ball was intermittently interrupted (red region). Second row: velocity of the fly. Third and fourth row: calcium activity of ensheathing glia in the MB (brown) and midline (dark orange). Vertical red lines indicates feeding events. **b-s** Same as a. Each panel represents a different trial from a total of 5 flies. The fly from which each trial was obtained is shown in the top left corner of each panel.

**Figure S20.**
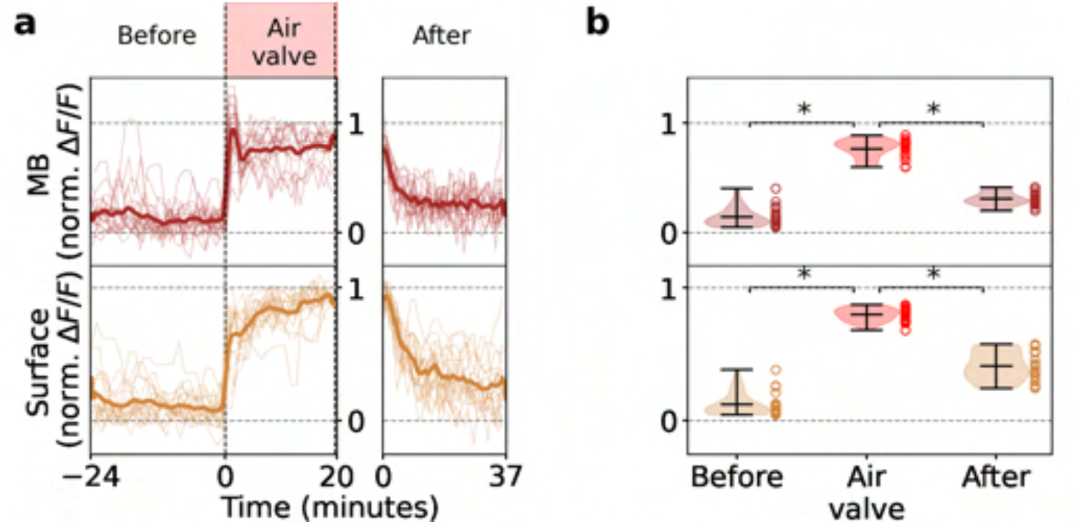
Glia activity in MB and midline increases with walking behavior. **a** Normalized fluorescence traces from the MB (brown) and midline (dark orange) from all the trials (thin lines, see Supplementary Fig. S19) before, during, and after the air valve was used to promote walking. Thick lines indicate the average normalized fluorescence of all trials. **b** Distribution of the mean fluorescence levels of all trials before, during, and after the air valve perturbation. Asterisks indicate statistical significance using t-test (p < 0.05).

**Figure S21.**
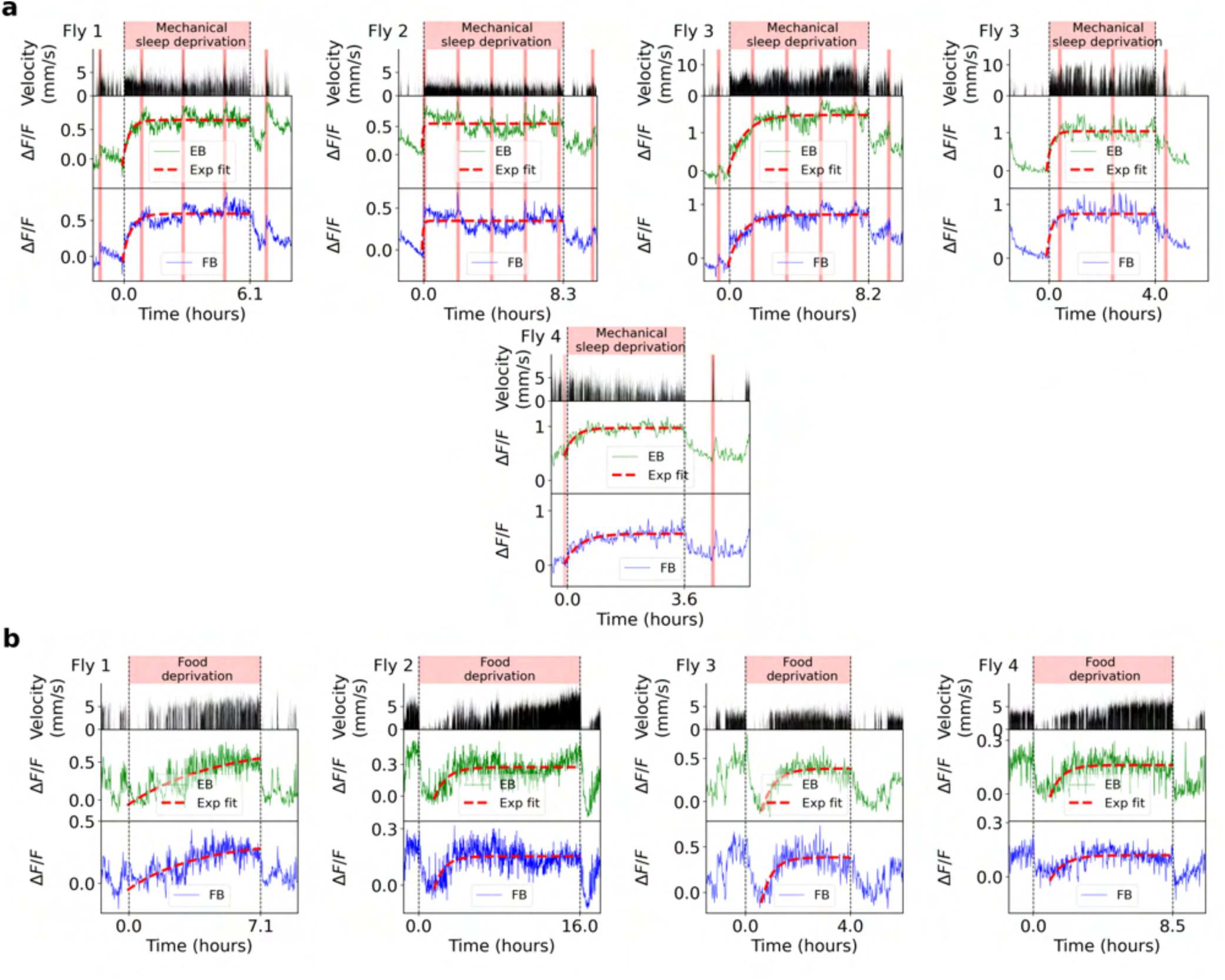
Glia activity during mechanical sleep deprivation (a) and food deprivation (b) in each trial. **a** Each panel shows the velocity over time (second row), and glia activity in EB (third row) and FB (fourth row) during mechanical sleep deprivation (first row) for 5 trials. Vertical red lines indicate feeding events and exponential fits (red) are shown for visualization of saturation levels. The fly from which each trial was obtained is indicated in the top left corner of each panel. **b** Same as a, but during food deprivation, with a different set of flies.

**Figure S22.**
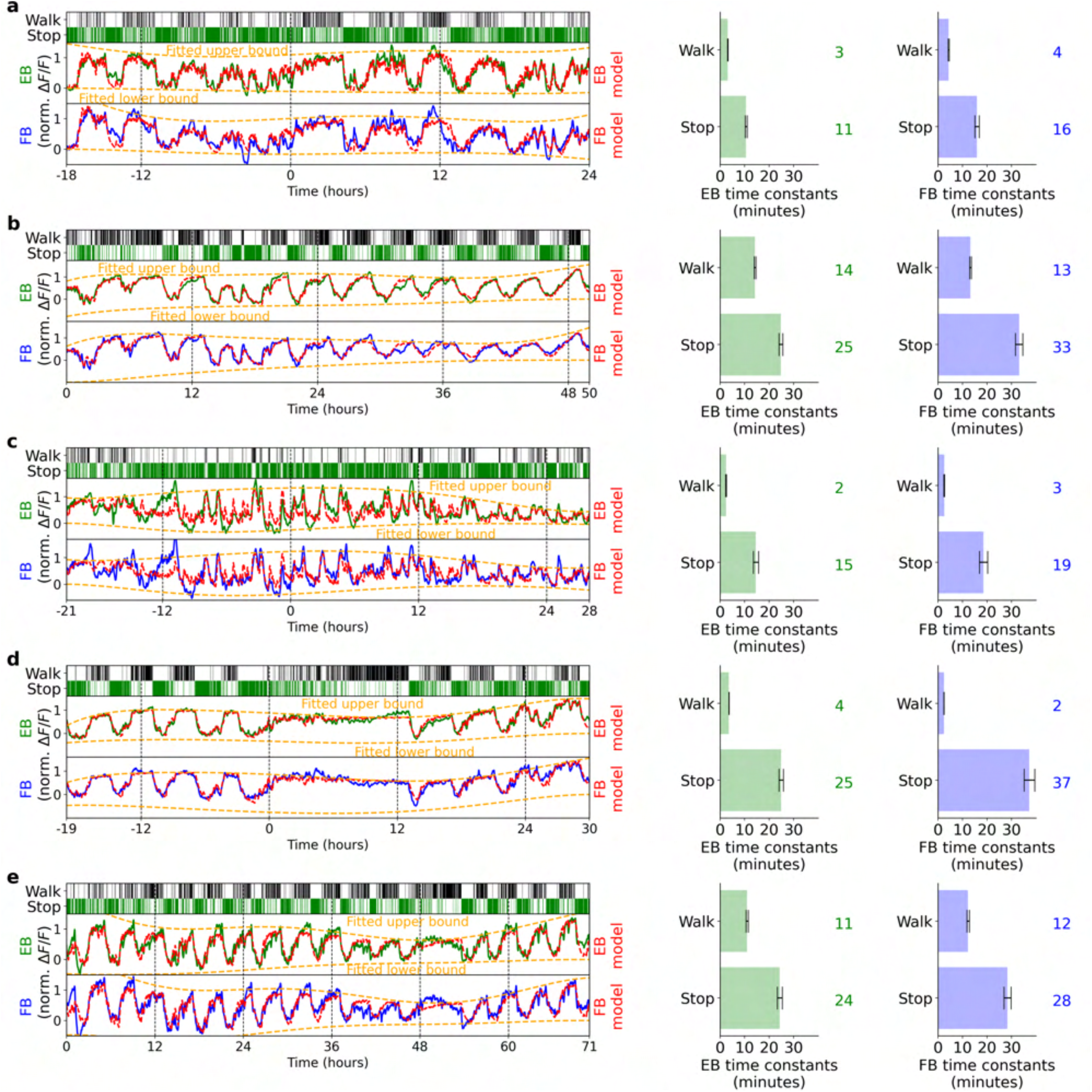
Fitting glia activity in EB and FB with homeostat 2-state model. **a** Left side: top row shows walking and stopping bouts of a fly. Second and third row: Normalized fluorescence in the EB (green) and FB (blue). Red lines show fitted model, while orange lines represent fitted upper and lower bounds of the model. Right side: fitted time constants from EB (green) and FB activity (blue). Grey lines indicate error bars of estimated time constants (see Methods). Green and blue numbers show rounded value of the fitted time constants. **b-e** Same as a. Each panel represents a fitted model for each fly.

**Figure S23.**
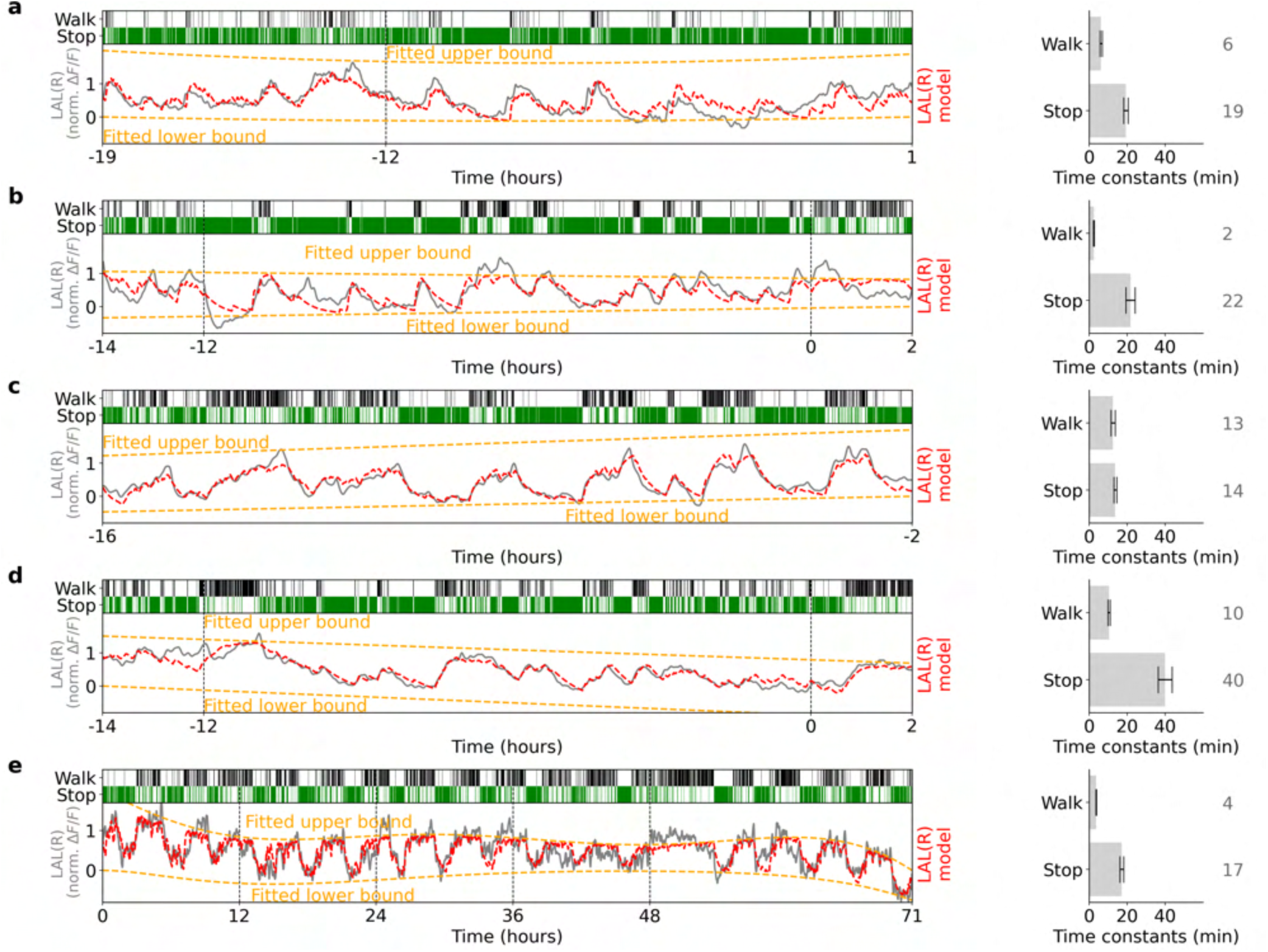
Fitting glia activity in the right LAL with homeostat 2-state model. **a** Left side: top row shows walking and stopping bouts of a fly. Second and third row: Normalized fluorescence in the LAL (grey). Red lines show fitted model, while orange lines represent fitted upper and lower bounds of the model. Right side: fitted time constants in LAL (grey). Grey lines indicate error bars of estimated time constants (see Methods). Grey numbers show rounded value of fitted time constants. **b-e** Same as a. Each panel represents a fitted model for each fly.

**Figure S24.**
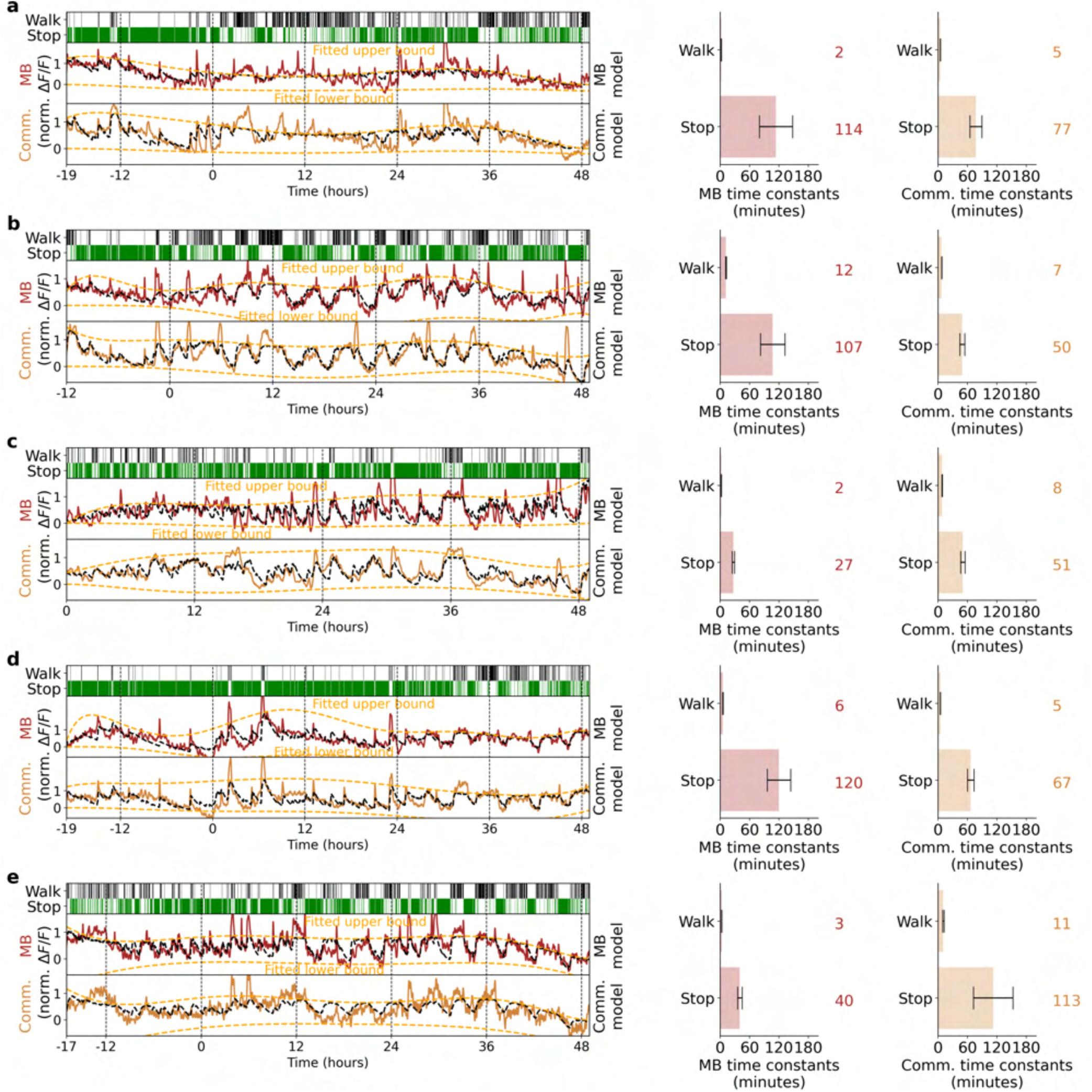
Fitting glia activity in MB and midline with homeostat 2-state model. **a** Left side: top row shows walking and stopping bouts of a fly. Second and third row: Normalized fluorescence in MB (brown) and midline (dark orange). Black lines show fitted model, while orange lines represent fitted upper and lower bounds of the model. Right side: fitted time constants in MB (brown) and midline activity (dark orange). Grey lines indicate error bars of estimated time constants (see Methods). Brown and dark orange numbers show rounded value of the fitted time constants. **b-e** Same as a. Each panel represents a fitted model for each fly.

**Figure S25.**
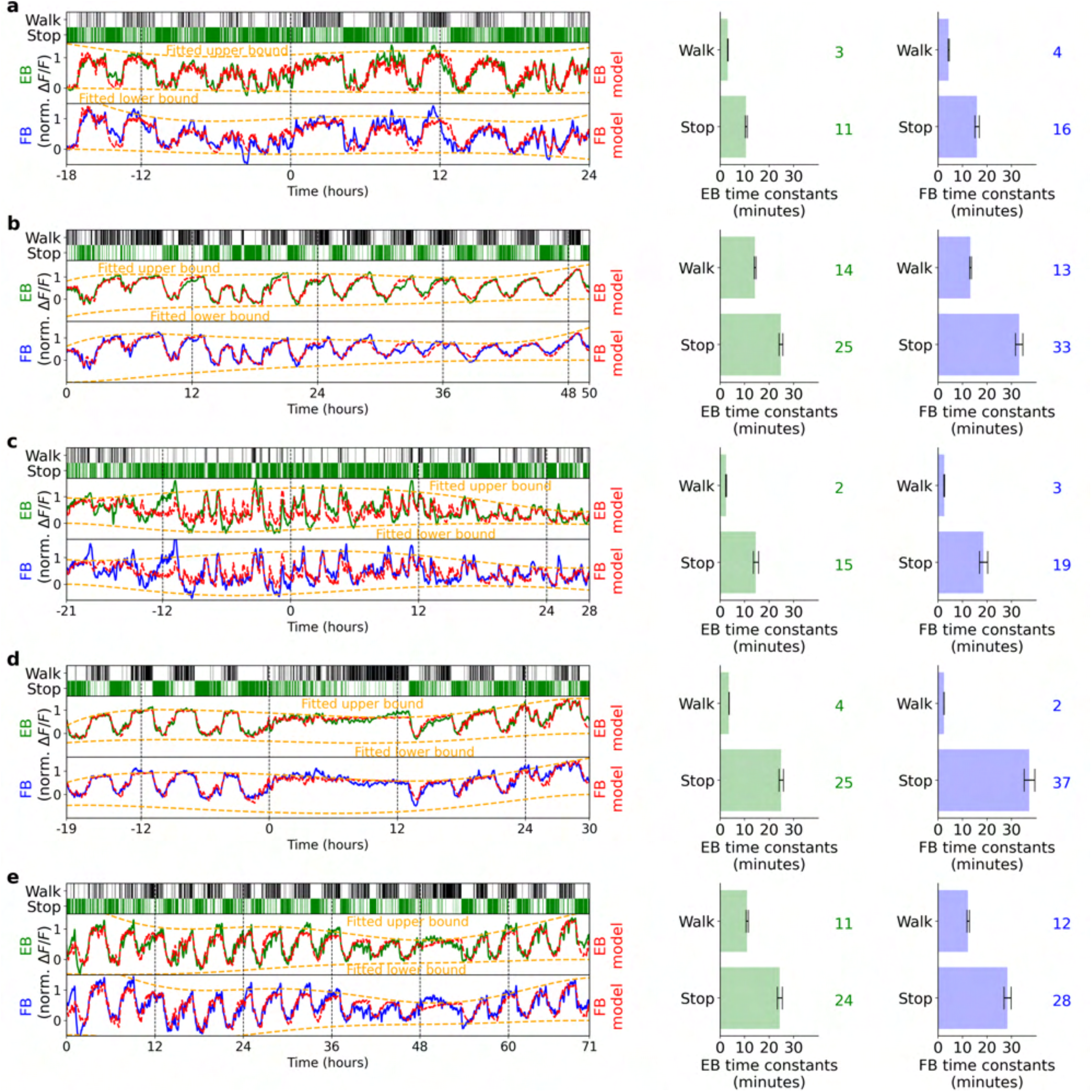
Fitting astrocytes activity in EB and FB with homeostat 2-state model. **a** Left side: top row shows walking and stopping bouts of a fly. Second and third row: Normalized fluorescence in the EB (green) and FB (blue). Red lines show fitted model, while orange lines represent fitted upper and lower bounds of the model. Right side: fitted time constants from EB (green) and FB activity (blue). Grey lines indicate error bars of estimated time constants (see Methods). Green and blue numbers show rounded value of the fitted time constants. **b-d** Same as a. Each panel represents a fitted model for each fly.

**Figure S26.**
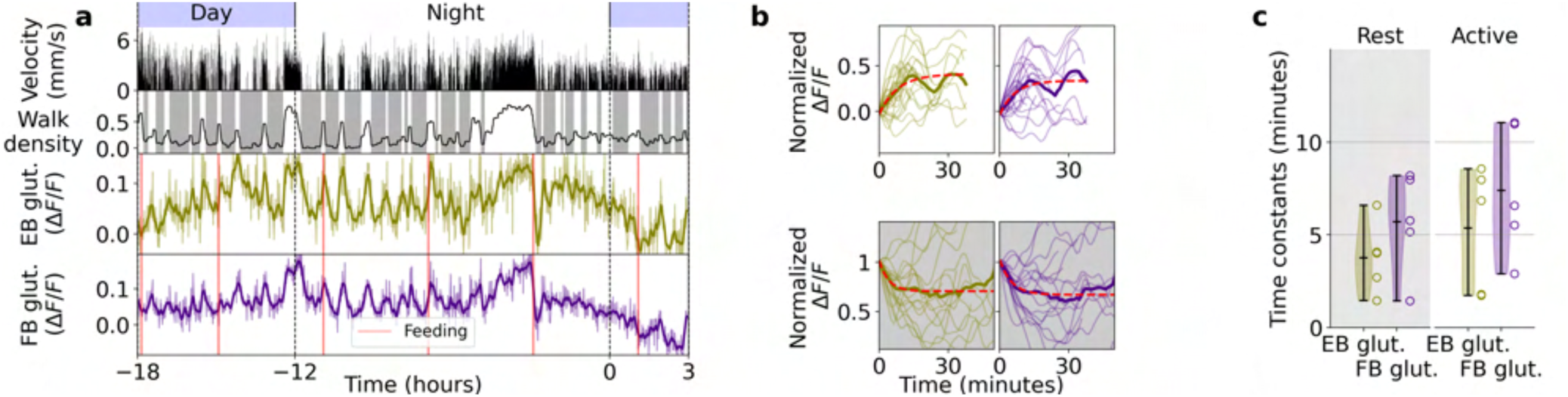
Glia glutamate dynamics. **a** Glutamate long-term imaging in EB and FB. Similar to Fig. 1i for glutamate. Top row: day and night cycle in VR. Second row: fly velocity. Third row: walk density with rest (grey region) and active (white region). Fourth and fifth row: extracellular glutamate activity in EB and FB, respectively. Thick lines indicate band-pass filtered signal (0.1 hours). **b** Single traces of increasing normalized glutamate activity (see Methods) during active state (thin lines), with average (thick line) and exponential fit (red) for EB and FB. **c** Time constants for exponential fit during active (white region) and rest (grey region) for extracellular glutamate in EB and FB in *N* = 5 flies.

**Figure S27.**
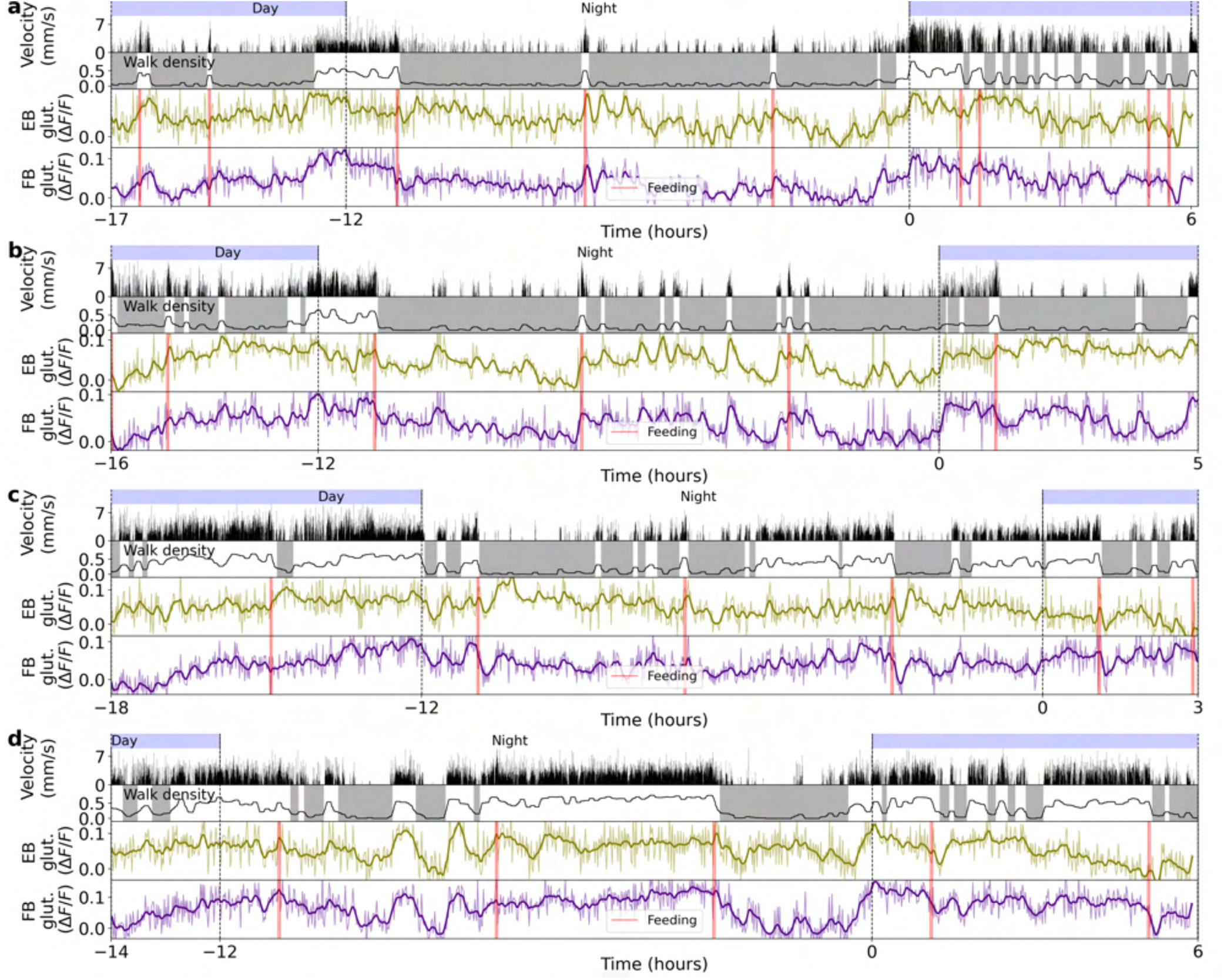
Four different recordings using the glutamate sensor iGluSnFR expressed in ensheathing glia. **a** Top row: day and night cycle in VR. Second row: velocity of the fly in 1 second bins. Third row: walk density (see Methods) and rest (grey region) and active (white region) epochs. Fourth and fifth row: Calcium activity of ensheathing glia in EB (green) and FB (blue). Thick lines indicate a low-pass filter with a 0.1 hours cut-off period. Vertical red lines represent feeding events. **b-d** Same as a. Each panel shows a different fly.

**Figure S28.**
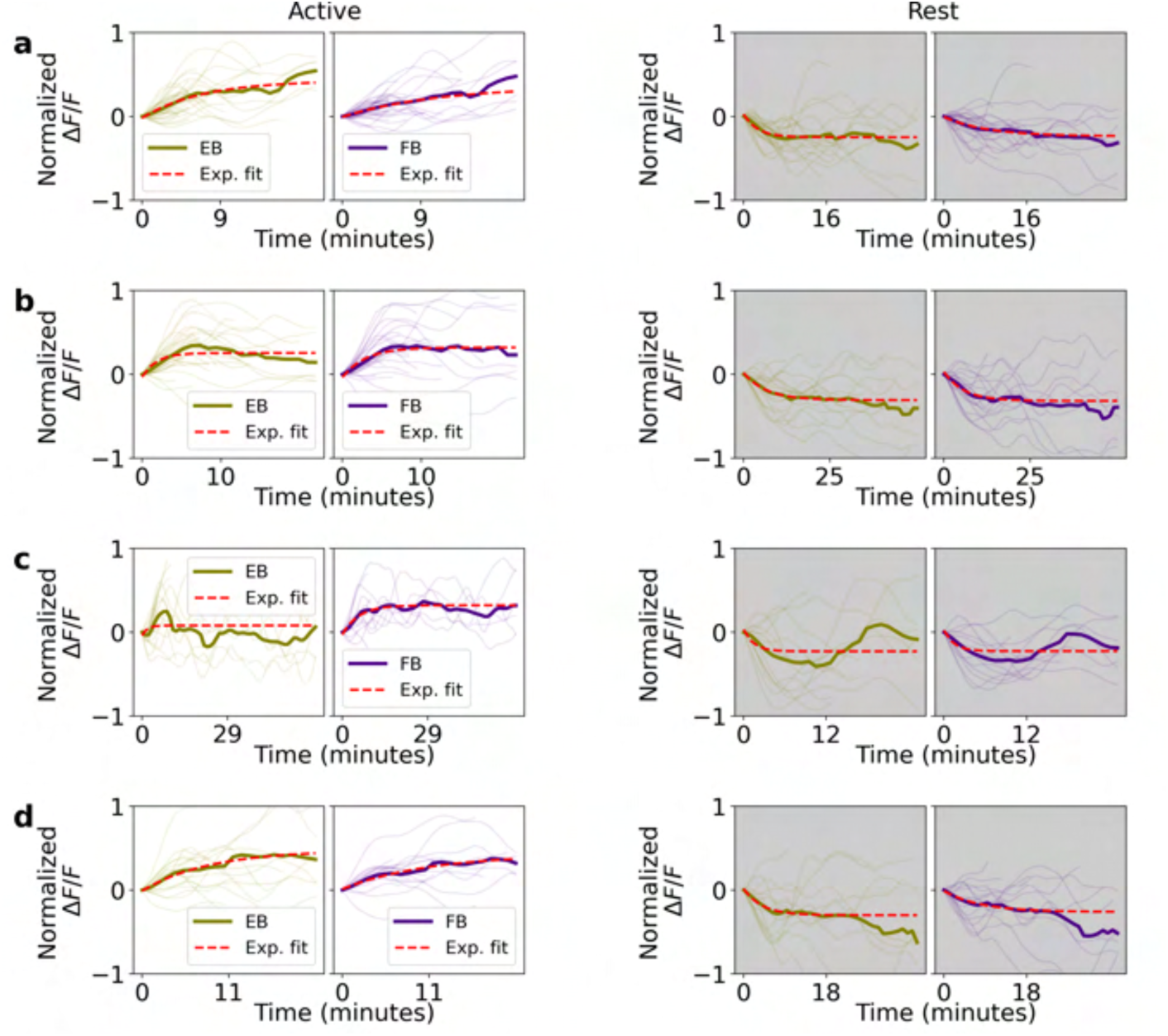
Normalized fluorescence traces recorded with glutamate sensor during active and rest epochs for 4 flies. **a** Left side: single (thin lines) and average (thick lines) normalized fluorescence traces in EB (olive) and FB (indigo) during active epochs. Red lines indicate exponential fit. Right side: same as left side, but during rest epochs. **b-d** Same as a. Each panel is from a different fly.

**Figure S29.**
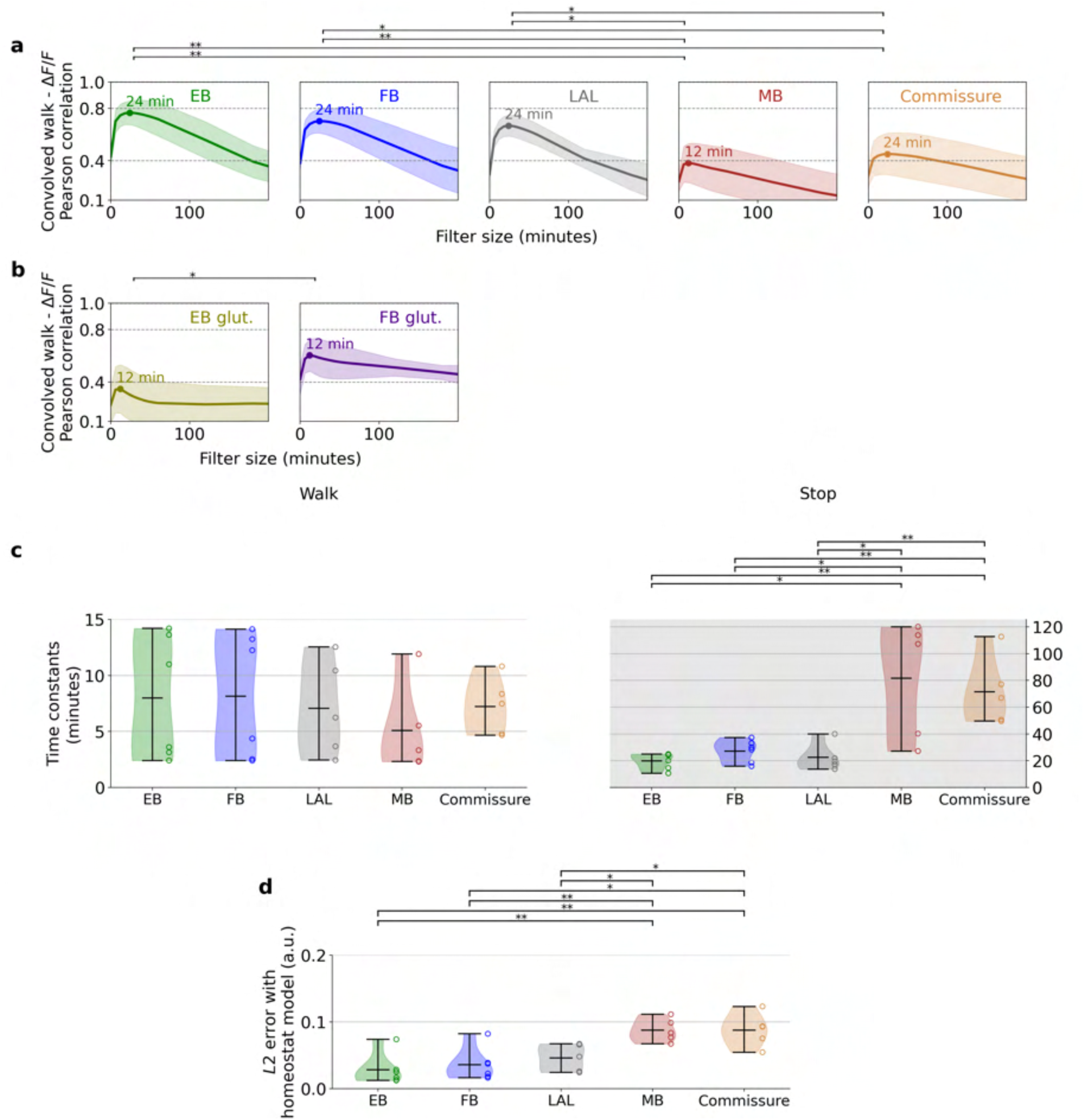
Comparison between glia activity in different neuropils and glutamate activity in the central complex **i** Pearson correlation between convolved ’walk’ and glia calcium activity in different neuropils for different filter sizes (*x*-axis) (see Fig. 6h for concept, and Methods). Thick-colored lines represent average of correlation curves, while light-colored areas represent standard deviation for all flies (*N* = 6 flies for EB and FB, and *N* = 5 flies for LAL, MB, and midline). Colored numbers indicate filter size at maximum correlation value in minutes. **b** Same as a, but for glutamate activity in the EB and FB (*N* = 5 flies). **c** Distributions of time constants for ’stop’ and ’walk’ states from model fitting (*N* = 6 flies in EB and FB, and *N* = 5 flies in LAL, MB, and midline). **d** *L*2 error between normalized fluorescence and the corresponding fitted homeostat 2-state model (*N* = 6 flies in EB and FB, and *N* = 5 flies in LAL, MB, and midline). Asterisks in each panel represent statistical significance using t-test (one asterisk for p-values < 0.05 and two asterisks for p-values < 0.005).

**Figure S30.**
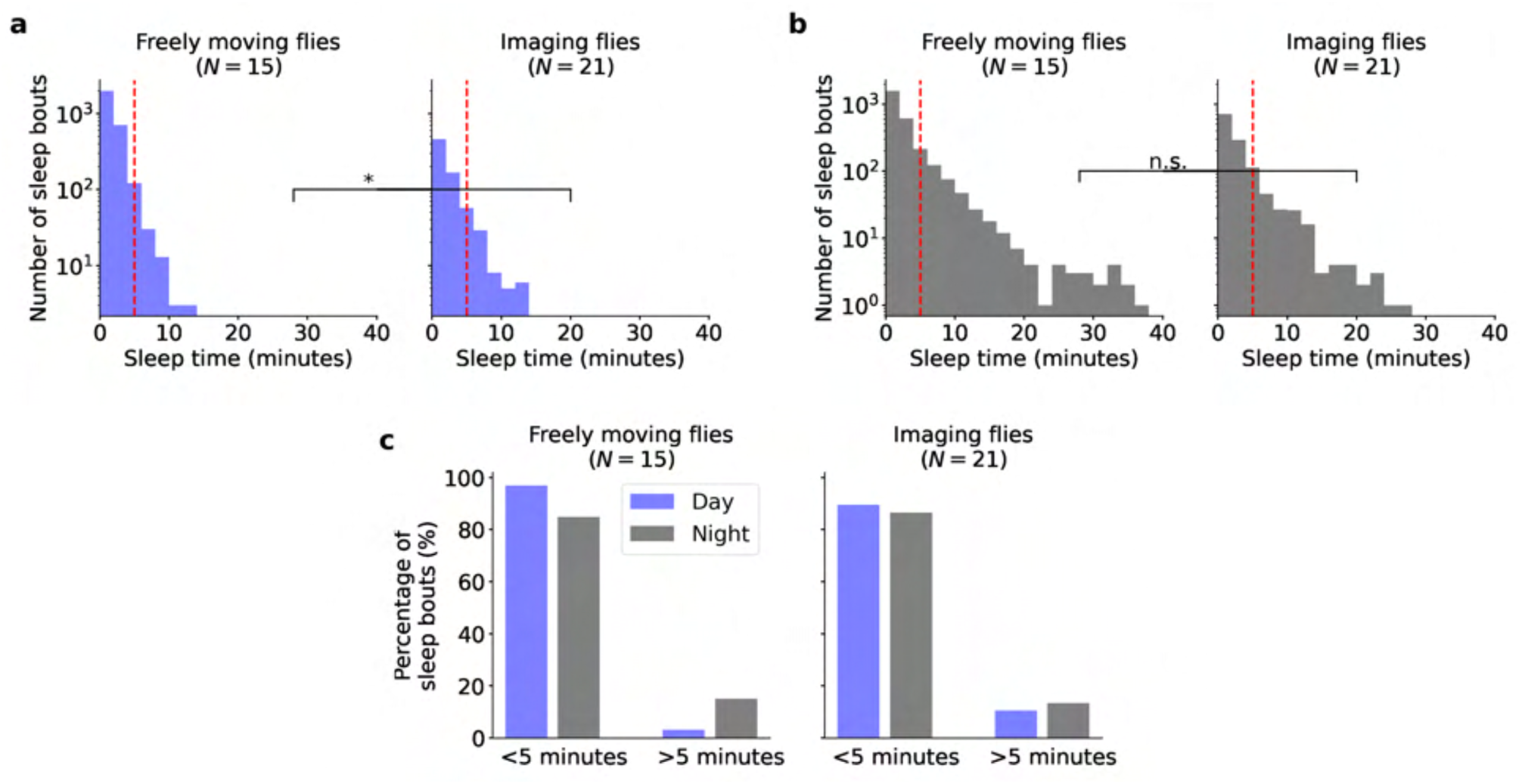
Comparison between sleep distribution in freely moving flies and flies during long-term imaging experiments. **a** Distribution of sleep bouts in freely moving flies (left) and flies during imaging experiments (right) during the day. The asterisk indicates statistical difference between the two distributions using the Kolmogorow-Smirnow test **b** Same as a but during the night. No statistical significance is found between these two distributions. **c** On the left panel, percentage of sleep bouts shorter (left) and larger (right) than 5 minutes during the day (blue) and night (grey) in freely moving flies. On the right panel, the same but for flies during long-term imaging experiments.

**Figure S31.**
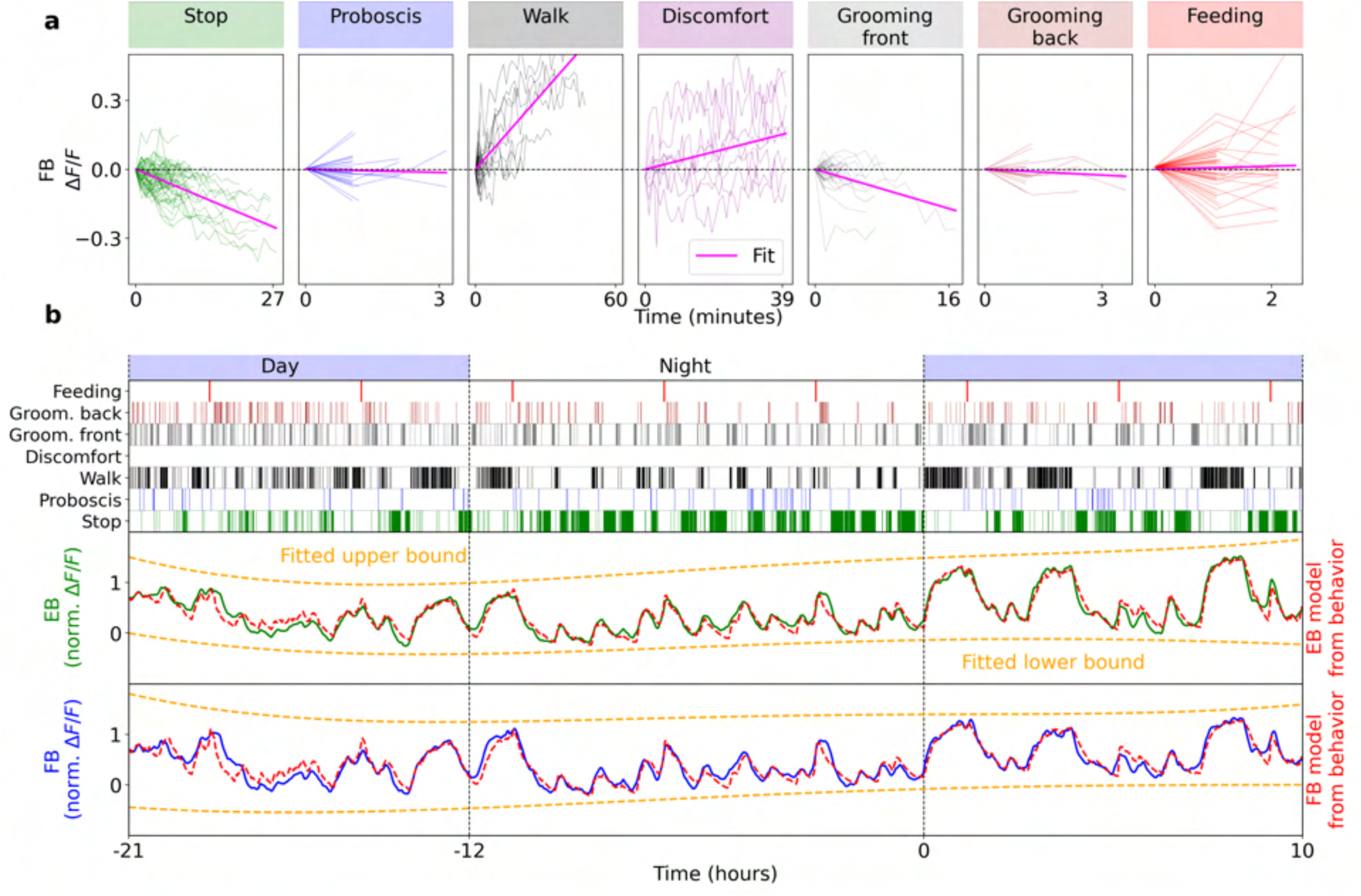
Fitting calcium activity with 7 behavioral states (seven-state model). **a** Determination of sign of weights of each behavior for model fitting. Fluorescence FB traces while fly is performing the respective behavior for at least to consecutive fluorescence recordings (2 minutes, combined across flies, see Methods). Purple shows linear fit, negative slope results in negative weight, and *vice versa*. Flies were often walking fast during feeding, likely resulting in increasing fluorescence. Note that the scale on y-axes differs between plots. **b** Model fitting with 7 states. Top: day-night cycle. Second row: classified fly behavior over time. Third row: band-pass filtered activity of glia in the EB (green line); red line shows model fit, orange lines show fitted correction of fluorescence levels. Fourth row: same as third row for FB.

**Figure S32.**
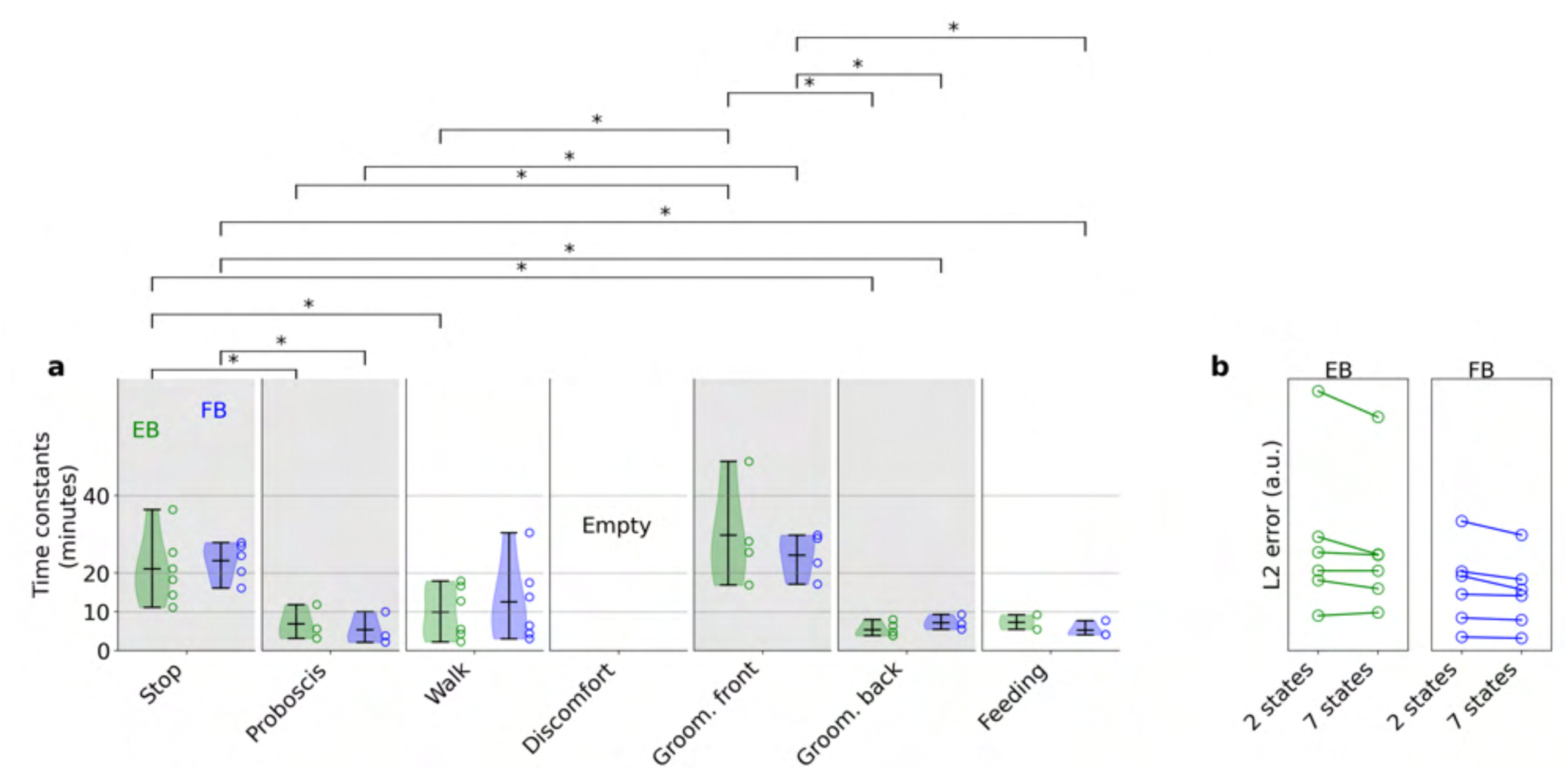
Significance levels for behaviors in 7-state model. Time constants of the 7 classified behaviors fitted in 7-state model for the EB (green) and FB (blue, Fig. S31a). Only time constants with an error of less than 20% were included in the histograms (see Methods). Asterisks include statistical significance between all possible pairs of fitted time constants, based on p-values lower than 0.05 using t-test. **b** Comparison between the 2-state and 7-state models in the EB (left) and FB (right). L2-error between fit and data for *N* = 6 flies.

**Figure S33.**
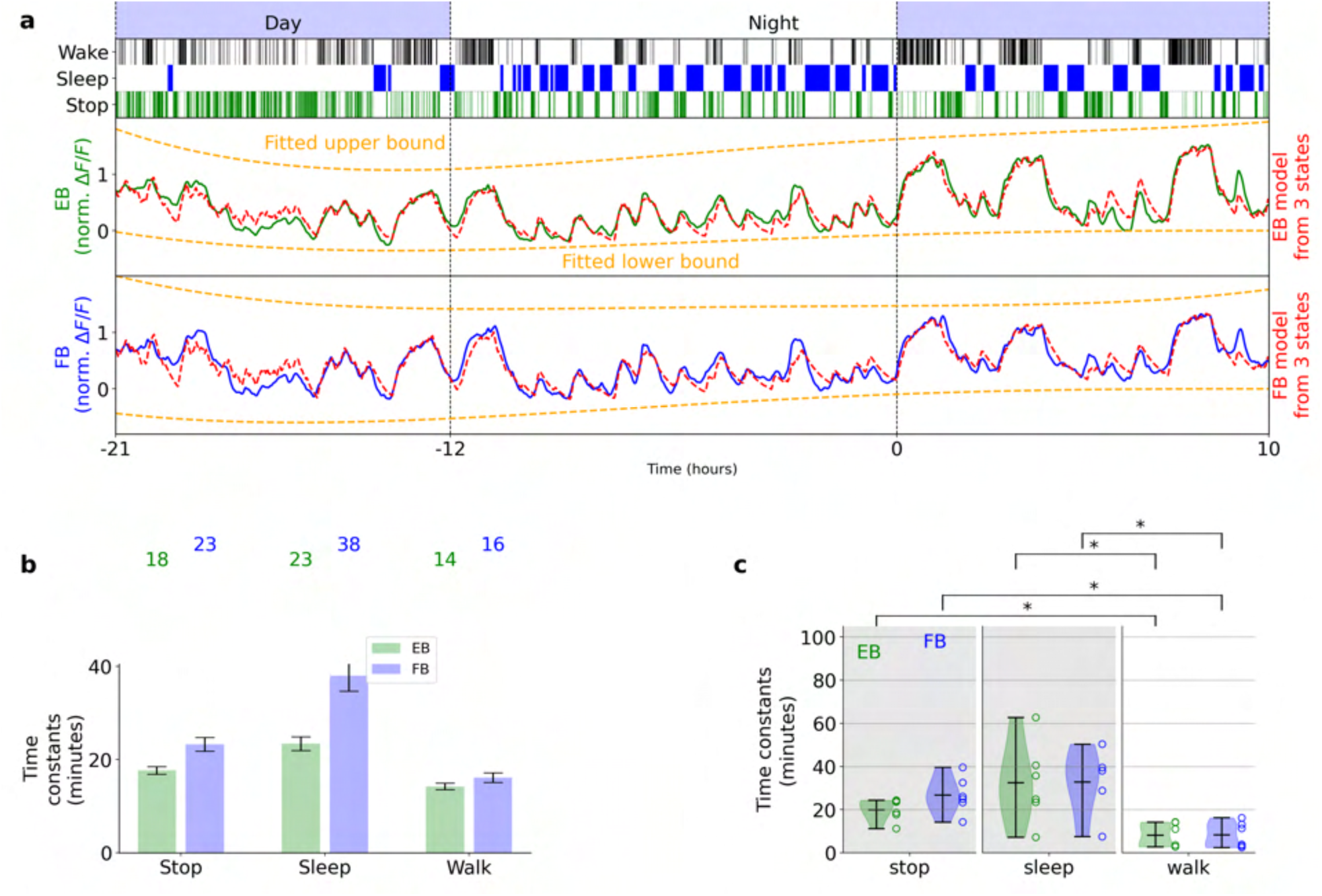
Fitting glia activity with homeostat 3-state model defining a ’sleep’ state as epochs where the fly was stopped for more than 5 minutes. **a** First row shows day and night cycle in VR, second row shows behavior of fly 1 extracted from ball velocity. Sleep is defined as epochs where the fly stops walking for at least 5 minutes. The third and fourth rows show fitting of the 3-state model (red line) and corresponding bounds (orange lines) in EB and FB, respectively. **b** Time constants of 3 states resulting from fitting in a for EB (green) and FB (blue). **c** Distribution of time constants for the 3 states for *N* = 6 flies. Asterisks represent statistical significance between states, with p-values lower than 0.05 using t-test.

**Figure S34.**
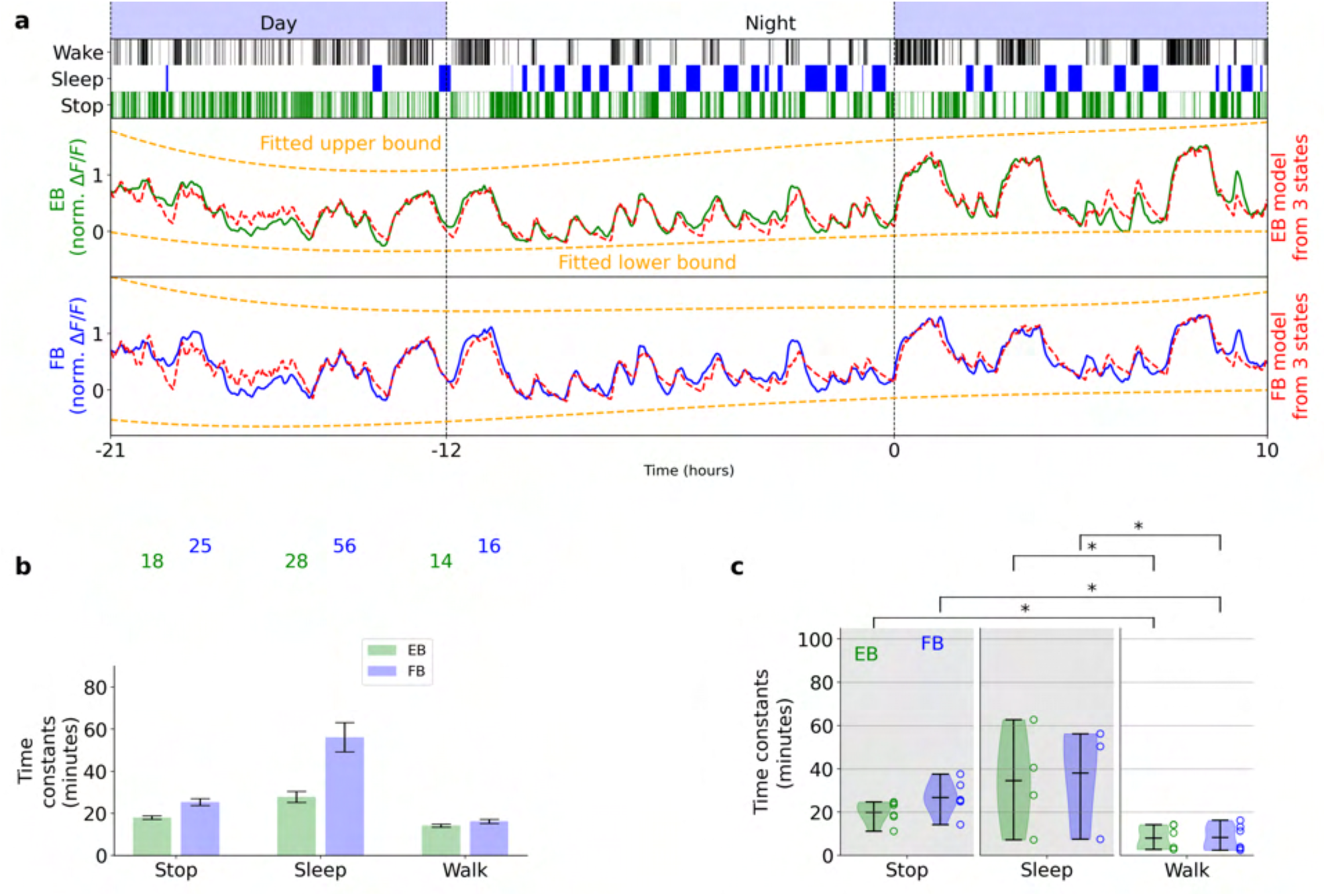
Fitting glia activity with homeostat 3-state model with a ’sleep’ state define as only after 5 minutes of immobility. Same as Supplementary Fig Fig. S33.

**Figure S35.**
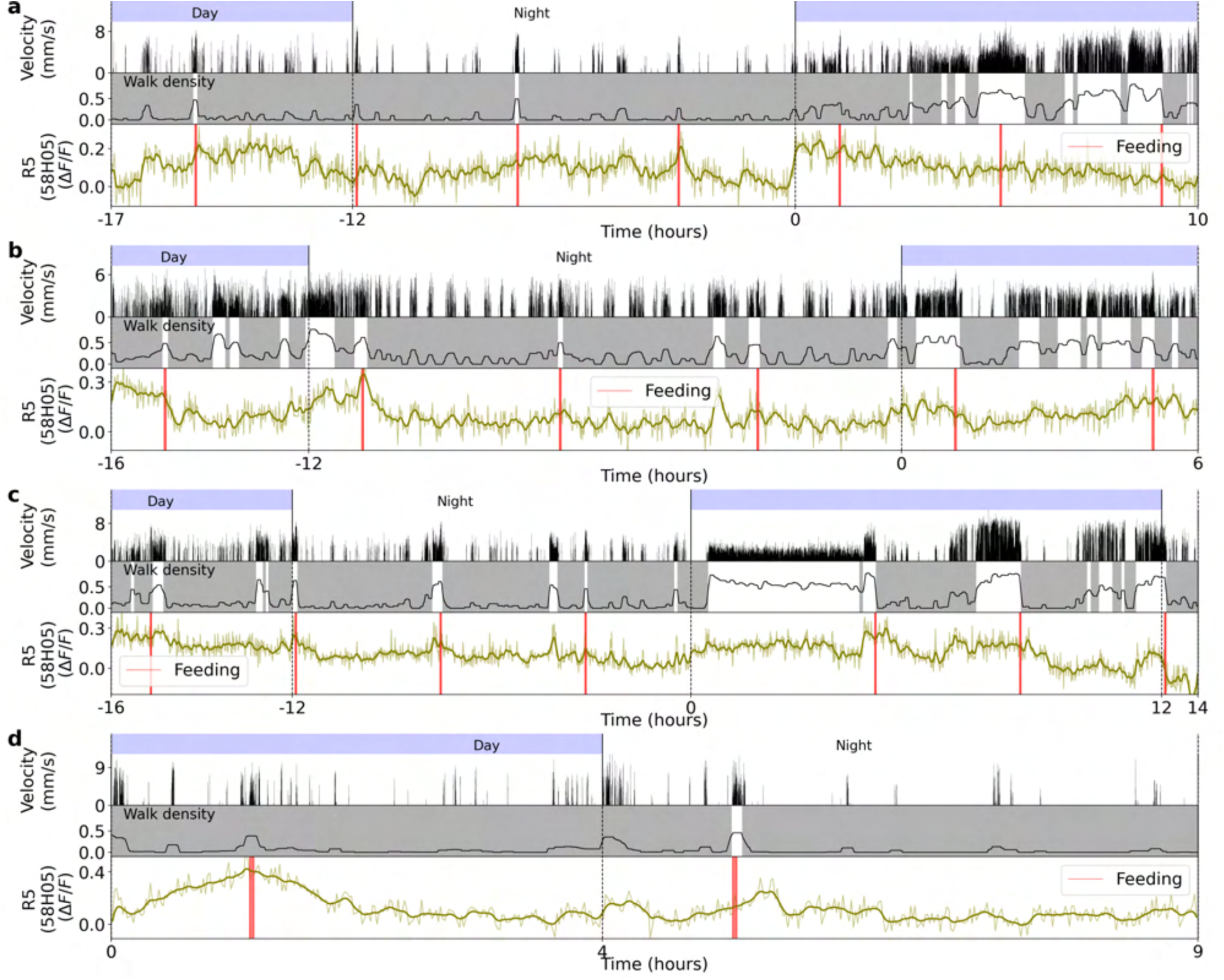
Four different long-term imaging recordings of calcium activity in R5 neurons labeled by 58H05-GAL4. **a** Top row: day and night cycle in VR. Second row: velocity of the fly in 1 second bins. Third row: walk density (see Methods) and rest (grey region) and active (white region) epochs. Fourth row: Calcium activity of R5 neurons. The thick line indicates a low-pass filter with a 0.1 hours cut-off period. Vertical red lines represent feeding events. **b-d** Same as a. Each panel shows a different fly.

**Figure S36.**
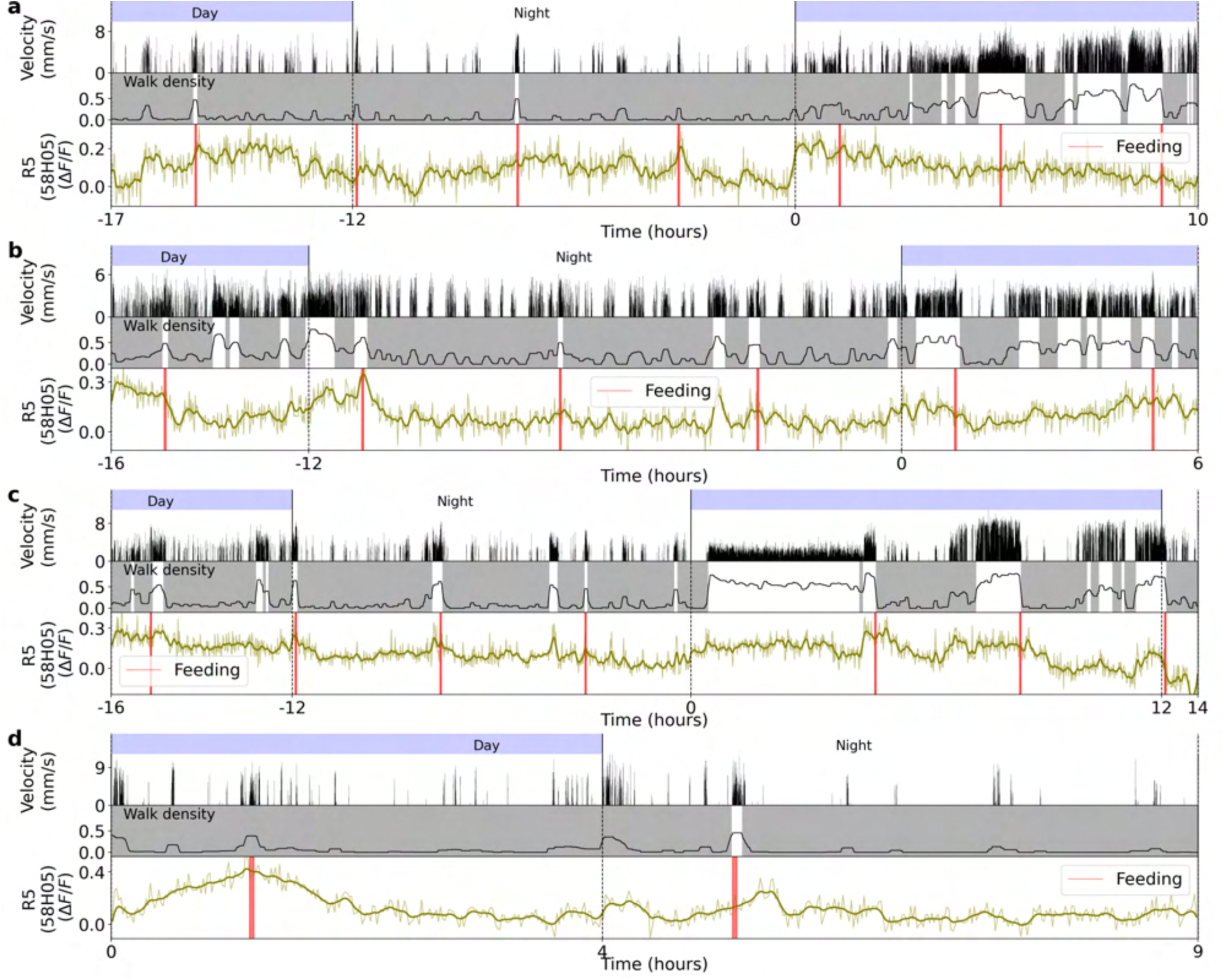
Normalized fluorescence traces during active and rest epochs for four flies in R5 neurons (labeled by GAL4-58H05). **a** Left side: single (thin lines) and average (thick lines) normalized fluorescence traces from activity in R5 neurons during active epochs. Red lines indicate exponential fit. Right side: the same as the left side, but during rest epochs. **b-d** Same as a. Each panel is from a different fly.

**Figure S37.**
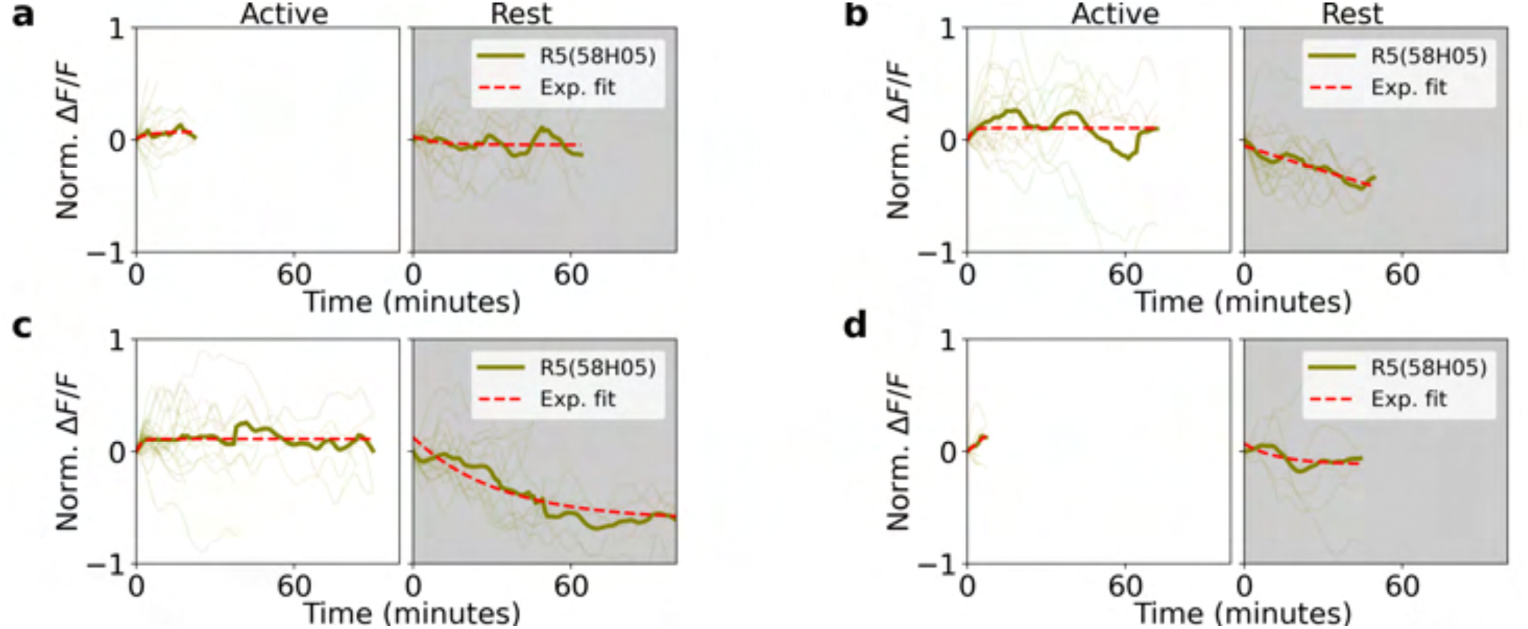
Fitting calcium activity of R5 neurons (labeled by GAL4-58H05) with the homeostat 2-state model. **a** Left side: the top row shows walking and stopping bouts of a fly. Second row: Normalized fluorescence from R5 neurons. Red lines show the fitted model, while orange lines represent the fitted upper and lower bounds of the model. Right side: fitted time constants from the model. Grey lines indicate error bars of the estimated time constants (see Methods), while colored numbers show the rounded value of the fitted time constants. **b-d** Same as a. Each panel represents a fitted model for each fly.

**Figure S38.**
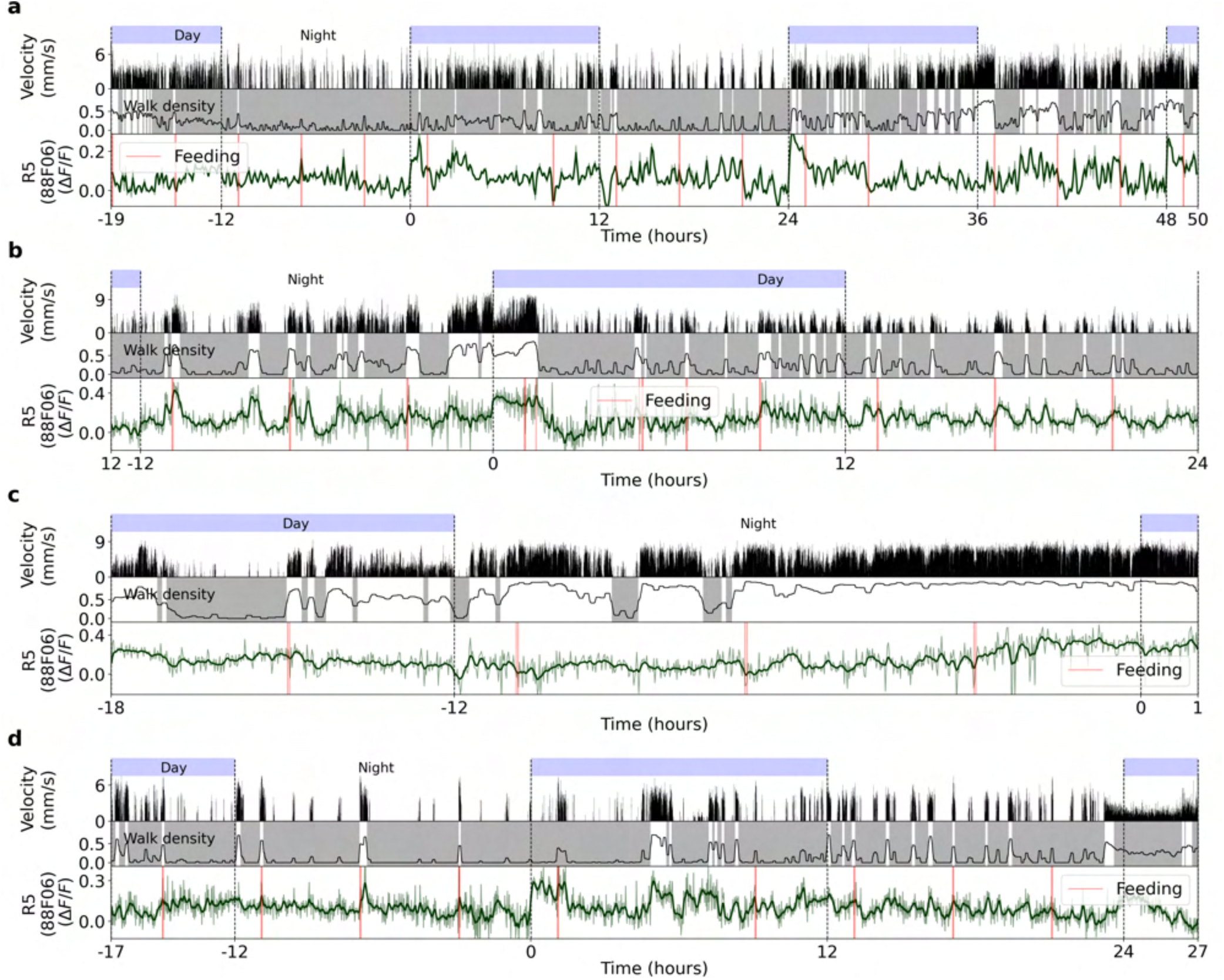
Four different recordings of calcium activity in R5 neurons labeled by GAL4-88F06. **a** Top row: day and night cycle in VR. Second row: velocity of fly in 1 second bins. Third row: walk density (see Methods), rest (grey region) and active (white region) epochs. Fourth row: Calcium activity of R5 neurons. Thick line indicates low-pass filtering with a 0.1 hours cut-off period. Vertical red lines represent feeding events. **b-d** Same as a. Each panel shows a different fly.

**Figure S39.**
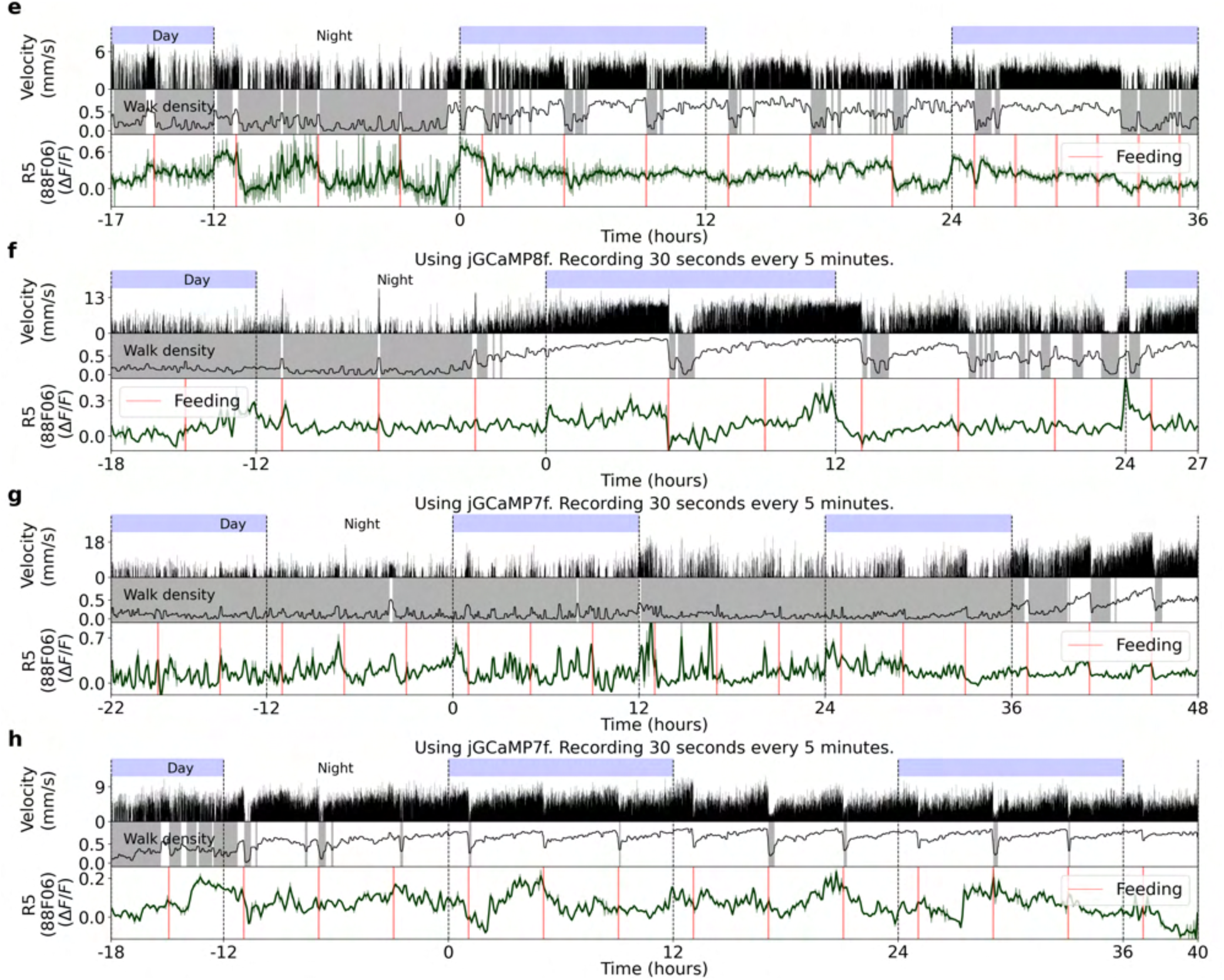
Four more recordings of calcium activity in R5 neurons labeled by GAL4-88F06. Same as Supplementary Fig. S38

**Figure S40.**
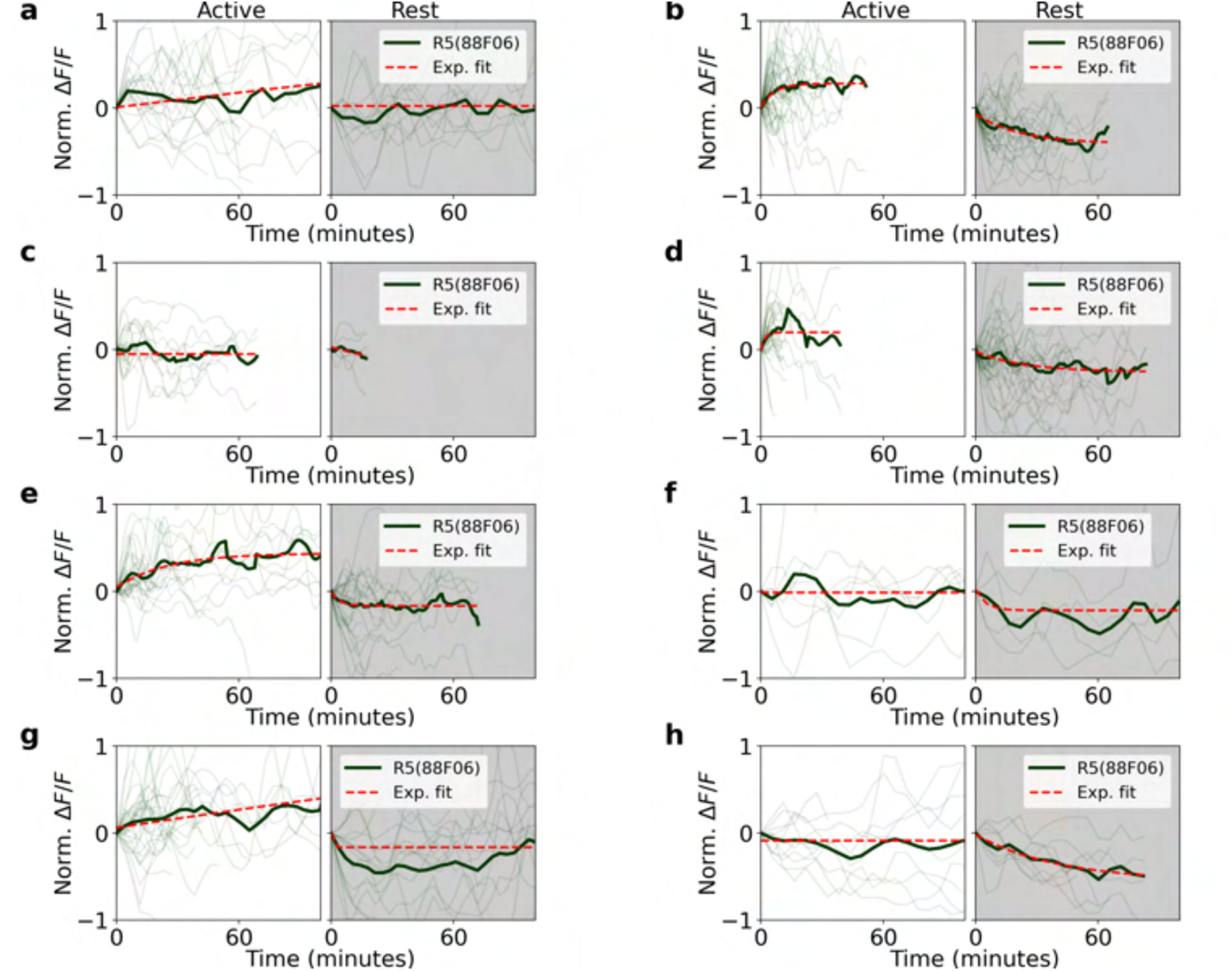
Normalized fluorescence traces during active and rest epochs for eight flies in R5 neurons (labeled by GAL4-88F06). **a** Left side: single (thin lines) and average (thick lines) normalized fluorescence traces of activity in R5 neurons during active epochs. Red lines indicate exponential fit. Right side: the same as left side, but during rest epochs. **b-d** Same as a. Each panel is a different fly.

**Figure S41.**
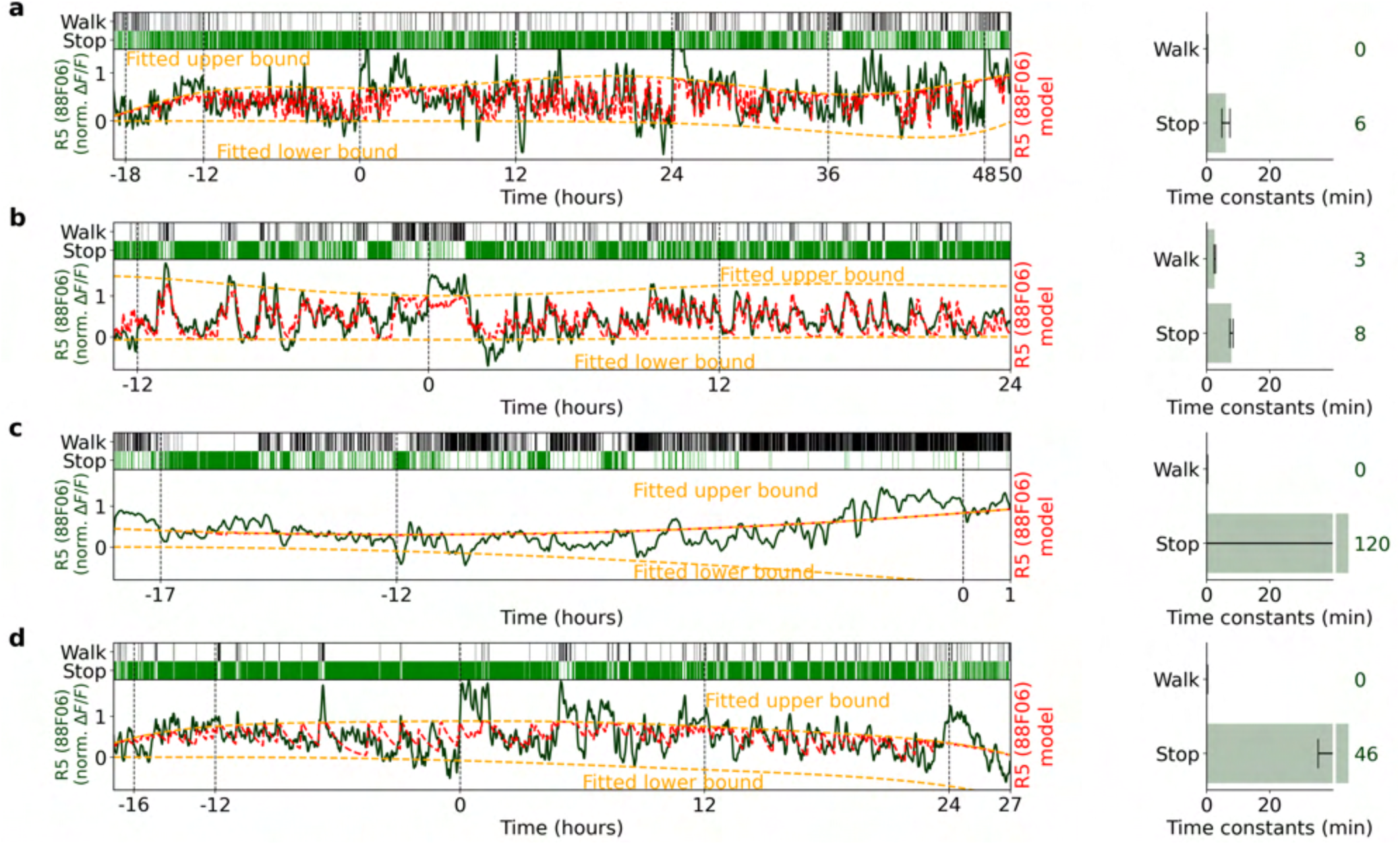
Fitting calcium activity of R5 neurons (labeled by GAL4-88F06) with homeostat 2-state model. **a** Left side: top row shows walking and stopping bouts of a fly. Second row: Normalized fluorescence of R5 neurons. Red lines show the fitted model, while orange lines represent fitted upper and lower bounds of model. Right side: time constants obtained from model fitting. Grey lines indicate error bars of estimated time constants (see Methods), while colored numbers show rounded values of fitted time constants. **b-d** Same as a. Each panel represents a fitted model for each fly.

**Figure S42.**
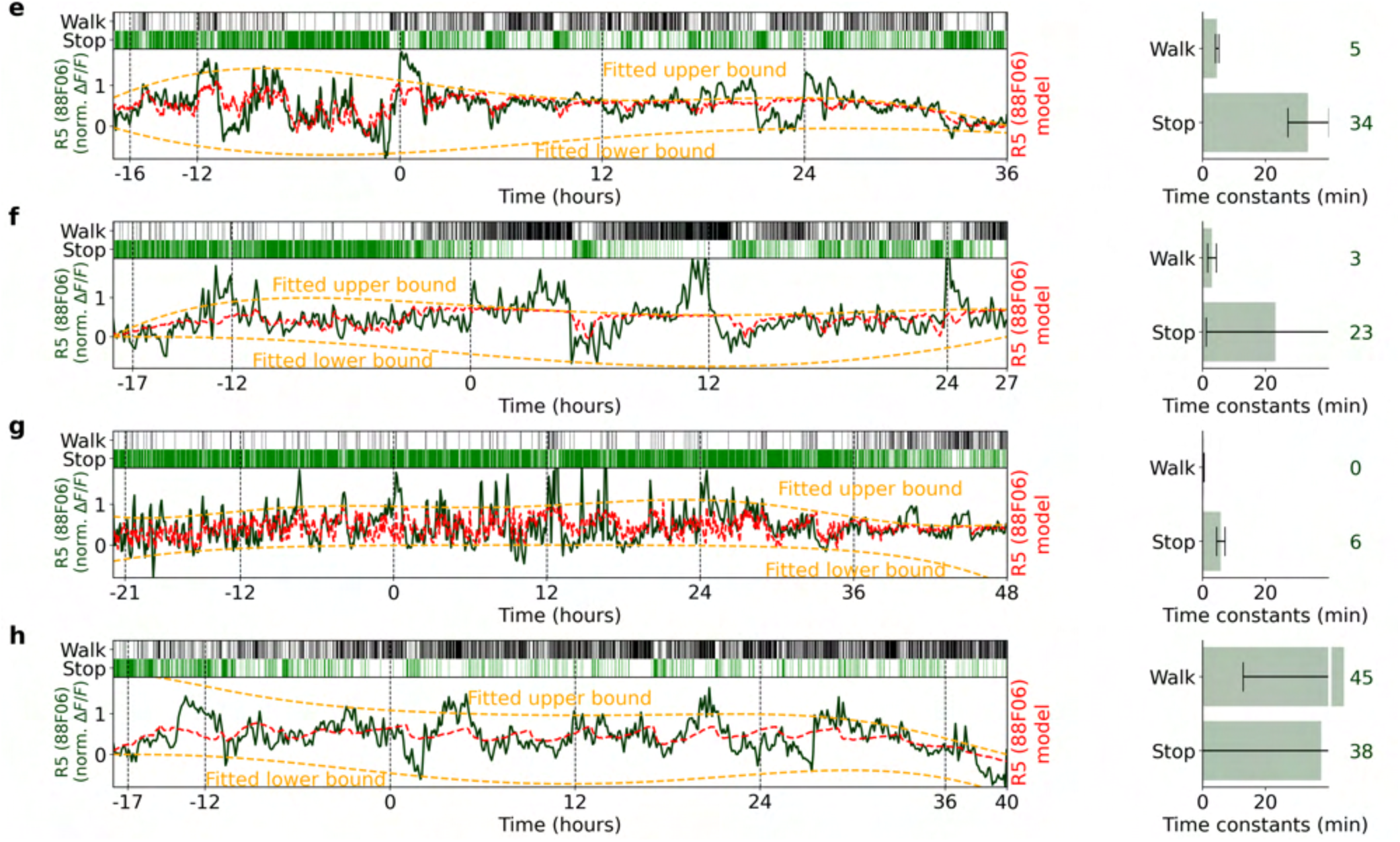
Four more recordings and fits of homeostat 2-state model with calcium activity of R5 neurons (labeled by GAL4-88F06). Same as Supplementary Fig. S41.

**Figure S43.**
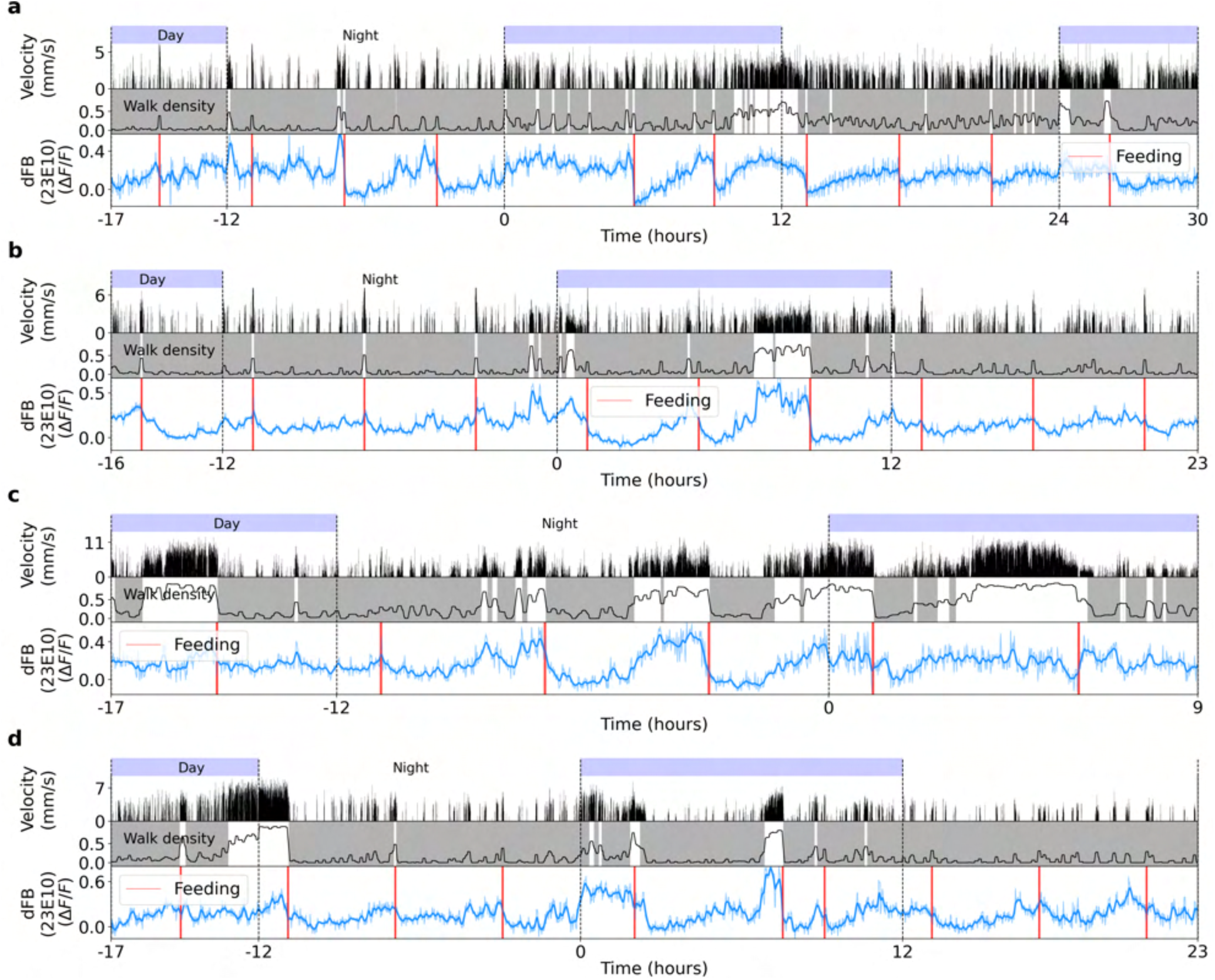
Four different recordings of calcium activity in dFB neurons labeled by GAL4-23E10. **a** Top row: day and night cycle in VR. Second row: velocity of fly in 1 second bins. Third row: walk density (see Methods), rest (grey region), and active (white region) epochs. Fourth row: Calcium activity of dFB neurons. The thick line indicates a low-pass filter with a 0.1 hours cut-off period. Vertical red lines represent feeding events. **b-d** Same as a. Each panel shows a different fly.

**Figure S44.**
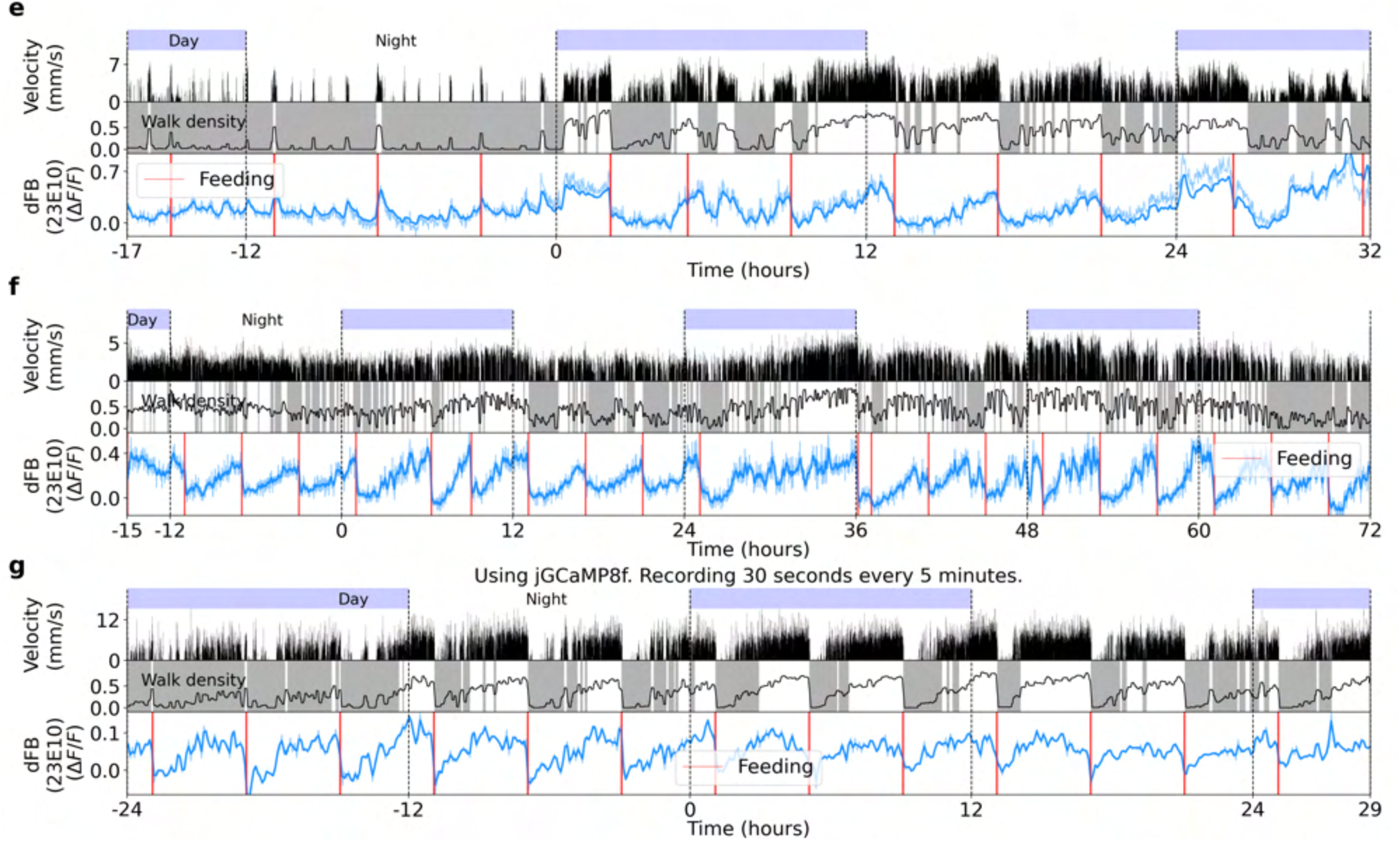
Three recordings of calcium activity in dFB neurons labeled by GAL4-23E10. Same as Supplementary Fig. S43

**Figure S45.**
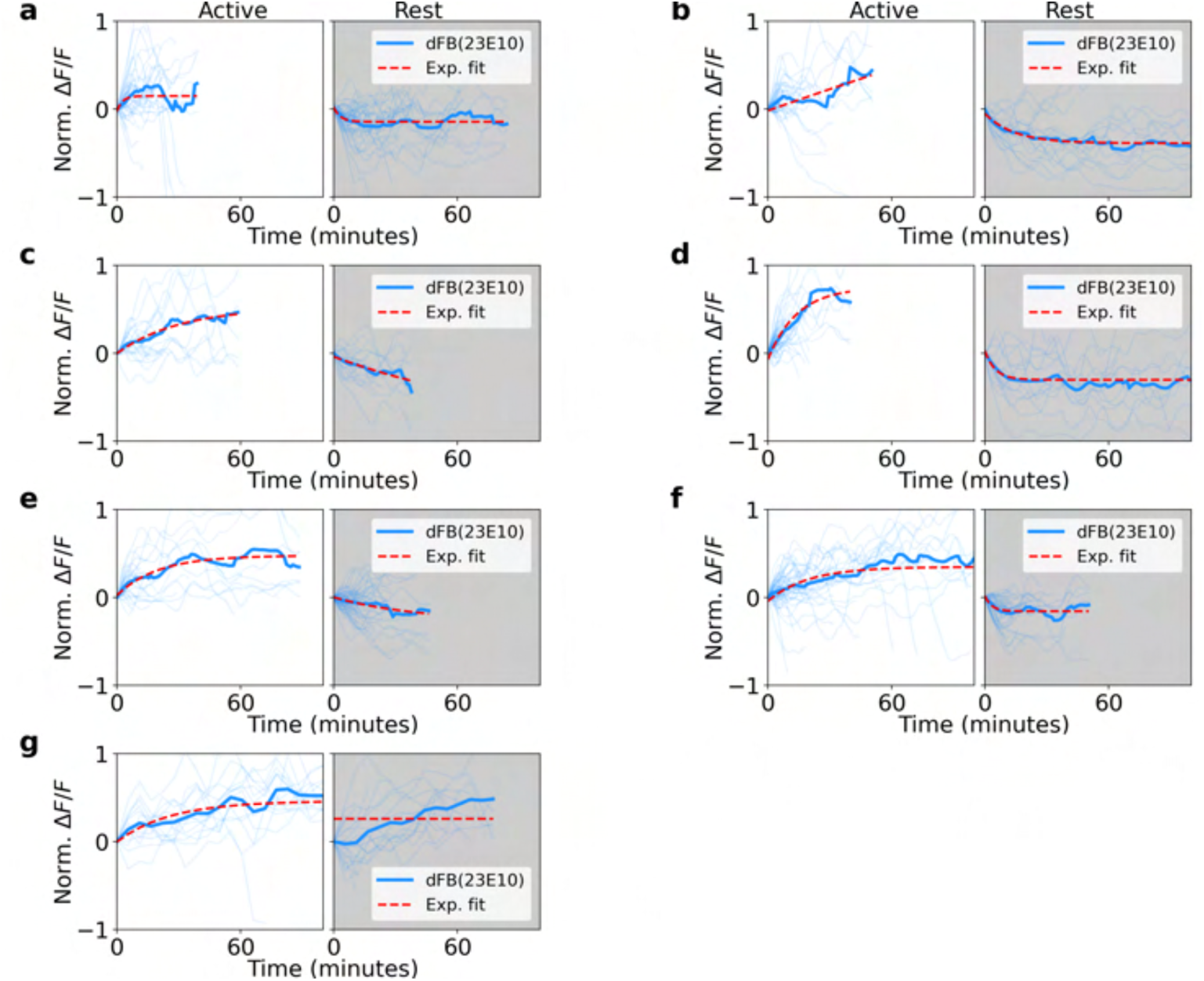
Normalized fluorescence traces during active and rest epochs for seven flies in dFB neurons (labeled by GAL4-23E10). **a** Left side: single (thin lines) and average (thick lines) normalized fluorescence traces of activity in dFB neurons during active epochs. Red lines indicate exponential fit. Right side: same as the left side, but during rest epochs. **b-d** Same as a. Each panel is from a different fly.

**Figure S46.**
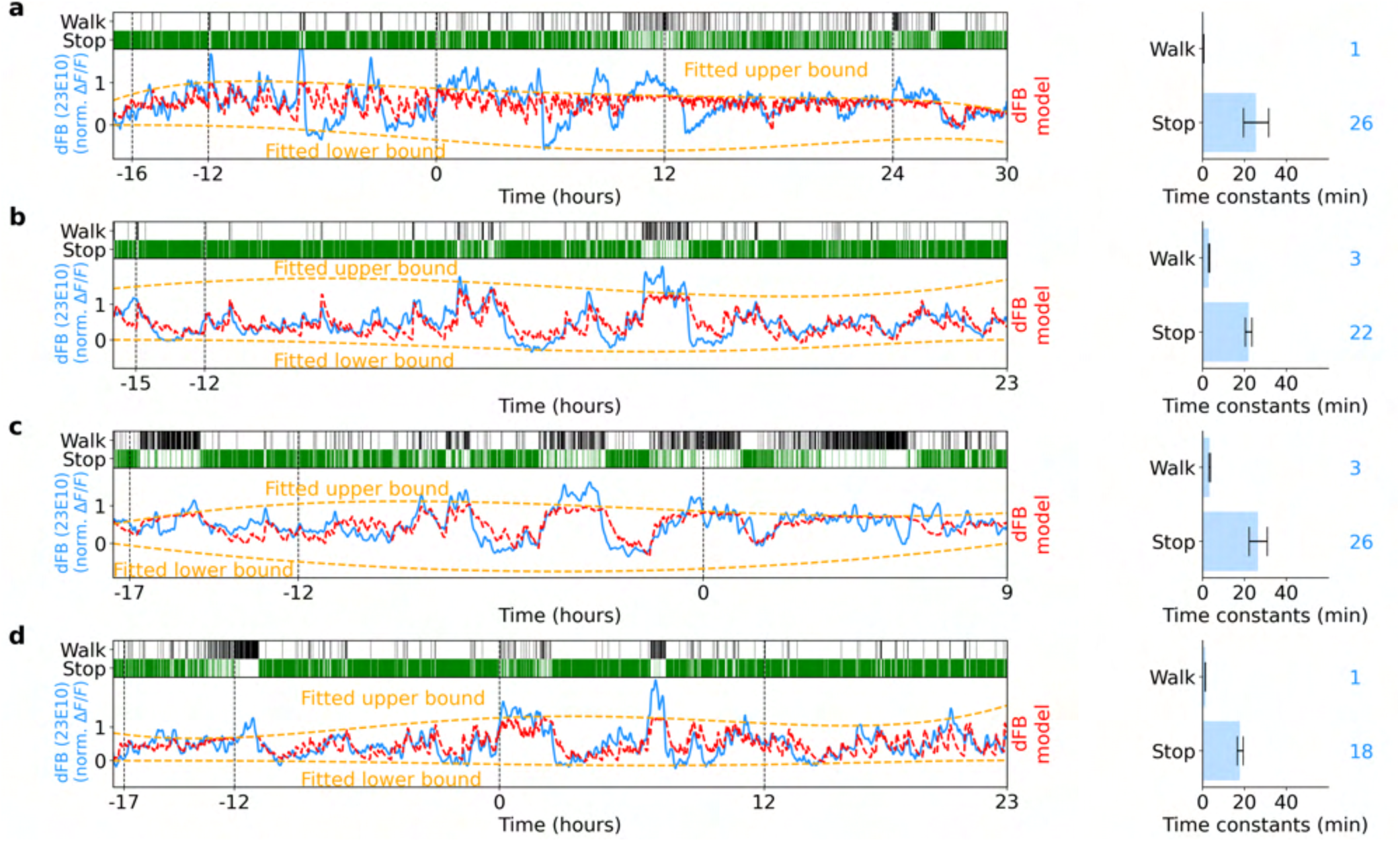
Fitting calcium activity of dFB neurons (labeled by GAL4-23E10) with homeostat 2-state model. **a** Left side: top row shows walk and stop bouts of a fly. Second row: Normalized fluorescence of dFB neurons. Red lines show fitted model, while orange lines represent fitted upper and lower bounds of the model. Right side: fitted model time constants. Grey lines indicate error bars of estimated time constants (see Methods), while colored numbers show the rounded value of the fitted time constants. **b-d** Same as a. Each panel represents a fitted model for each fly.

**Figure S47.**
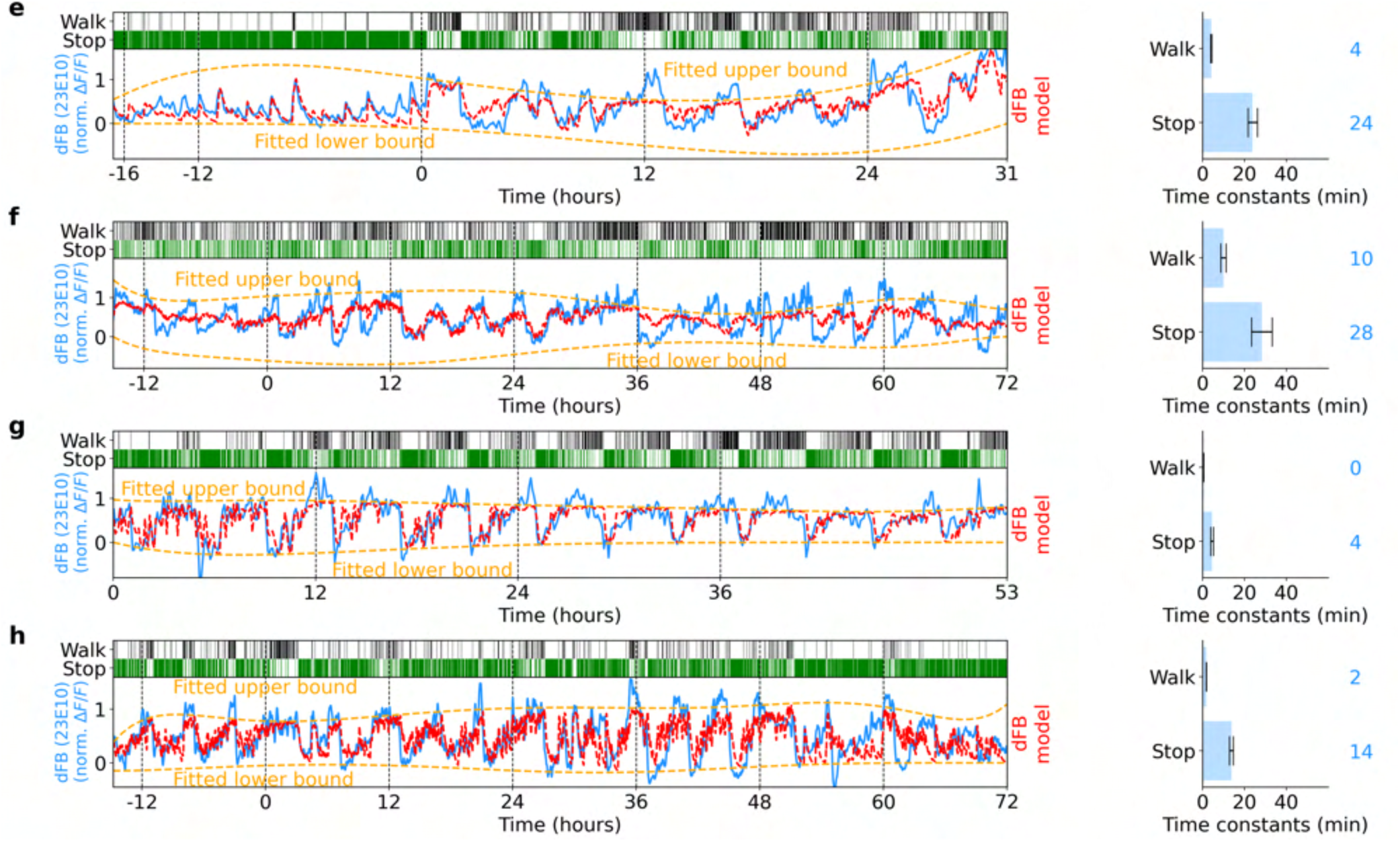
Four more experiments and fits of the homeostat 2-state model with calcium activity of dFB neurons (labeled by GAL4-23E10). Same as Supplementary Fig. S46.

**Figure S48.**
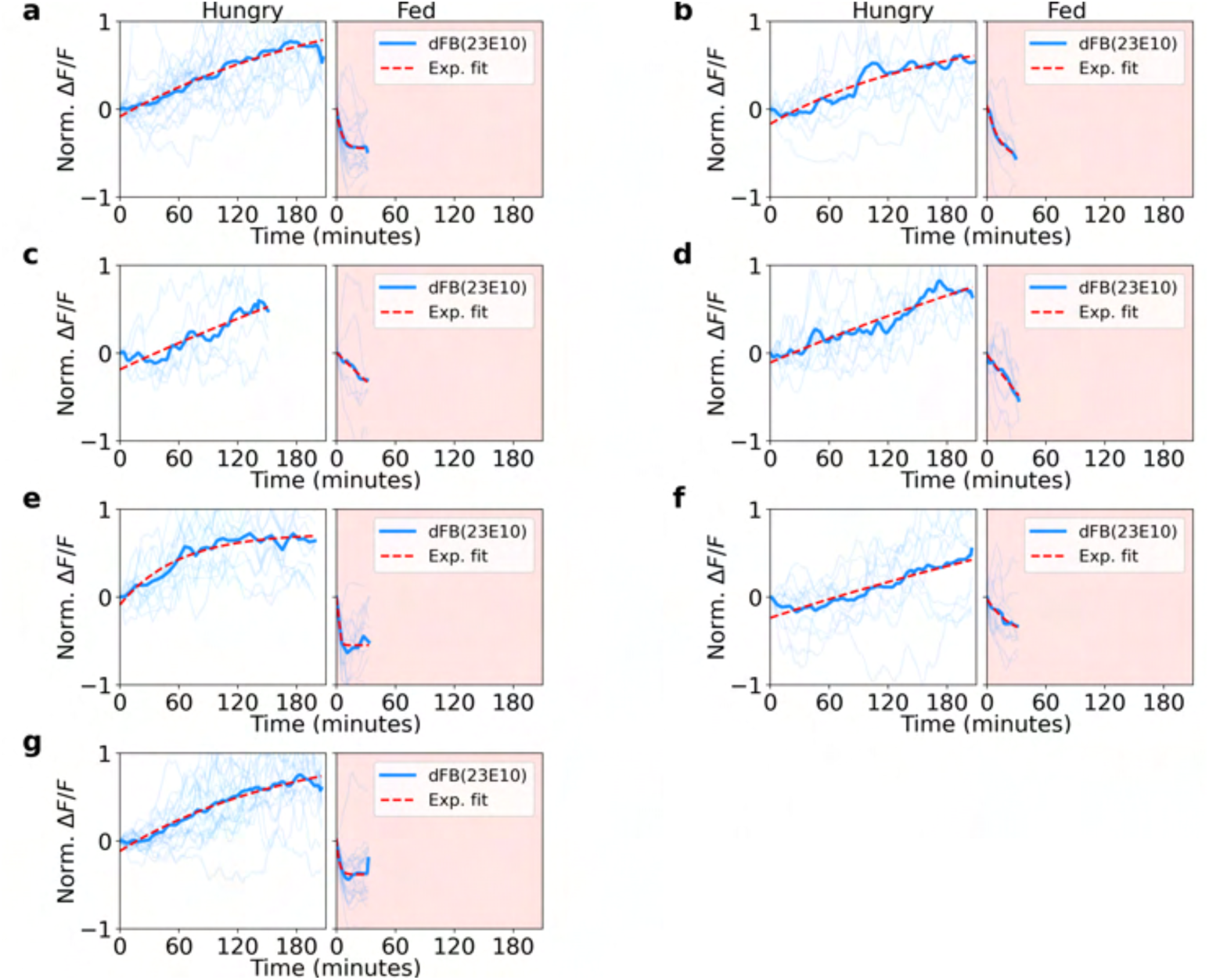
Normalized fluorescence traces during hungry and fed epochs for seven flies in dFB neurons (labeled by GAL4-23E10). **a** Left side: single (thin lines) and average (thick lines) normalized fluorescence traces of activity in dFB neurons during hungry epochs. Red lines indicate exponential fit. Right side: same as the left side, but during fed epochs. **b-d** Same as a. Each panel is from a different fly.

**Figure S49.**
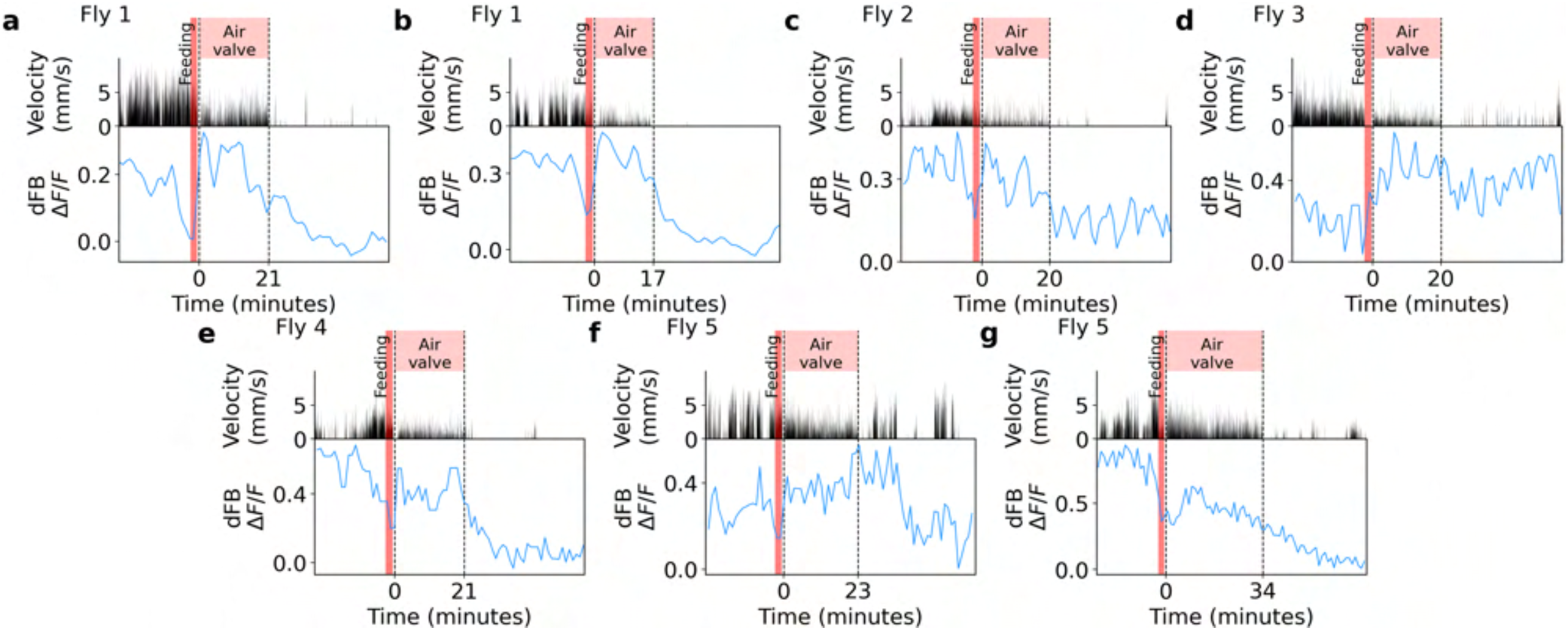
Trials where air supply of the ball was intermittently interrupted to promote walking after feeding, while activity in dFB neurons was recorded. **a** First row: times where the ball was perturbed by the air valve (red region). Second row: velocity of the fly. Third row: Calcium activity in dFB neurons. The vertical red line indicates feeding event before perturbing the ball. **b-g** Same as a. Each panel represents a different trial. The fly from which each trial was recorded is shown in the top left corner of each panel.

**Figure S50.**
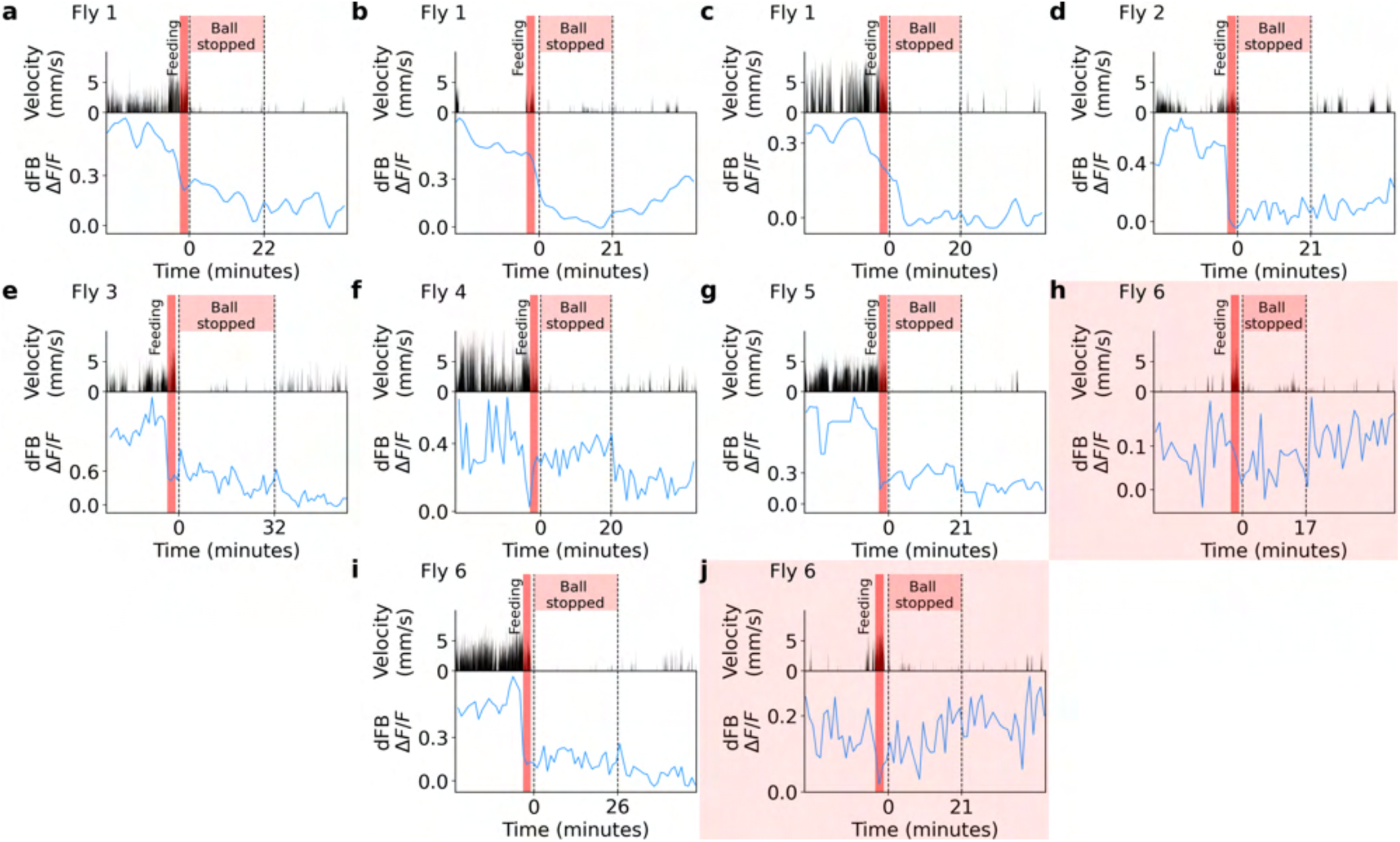
Trials where the ball was blocked after feeding while activity in dFB neurons was recorded. **a** First row: time where the ball was stopped (red region). Second row: velocity of the fly. Third row: Calcium activity dFB neurons. The vertical red line indicates the feeding event before blocking the ball. **b-j** Same as a. Each panel represents a different trial. Trials h and j were not considered for the analysis (highlighted in red) because dFB neurons had low activity levels before feeding. The fly from which each trial was recorded is shown in the top left corner of each panel.

**Figure S51.**
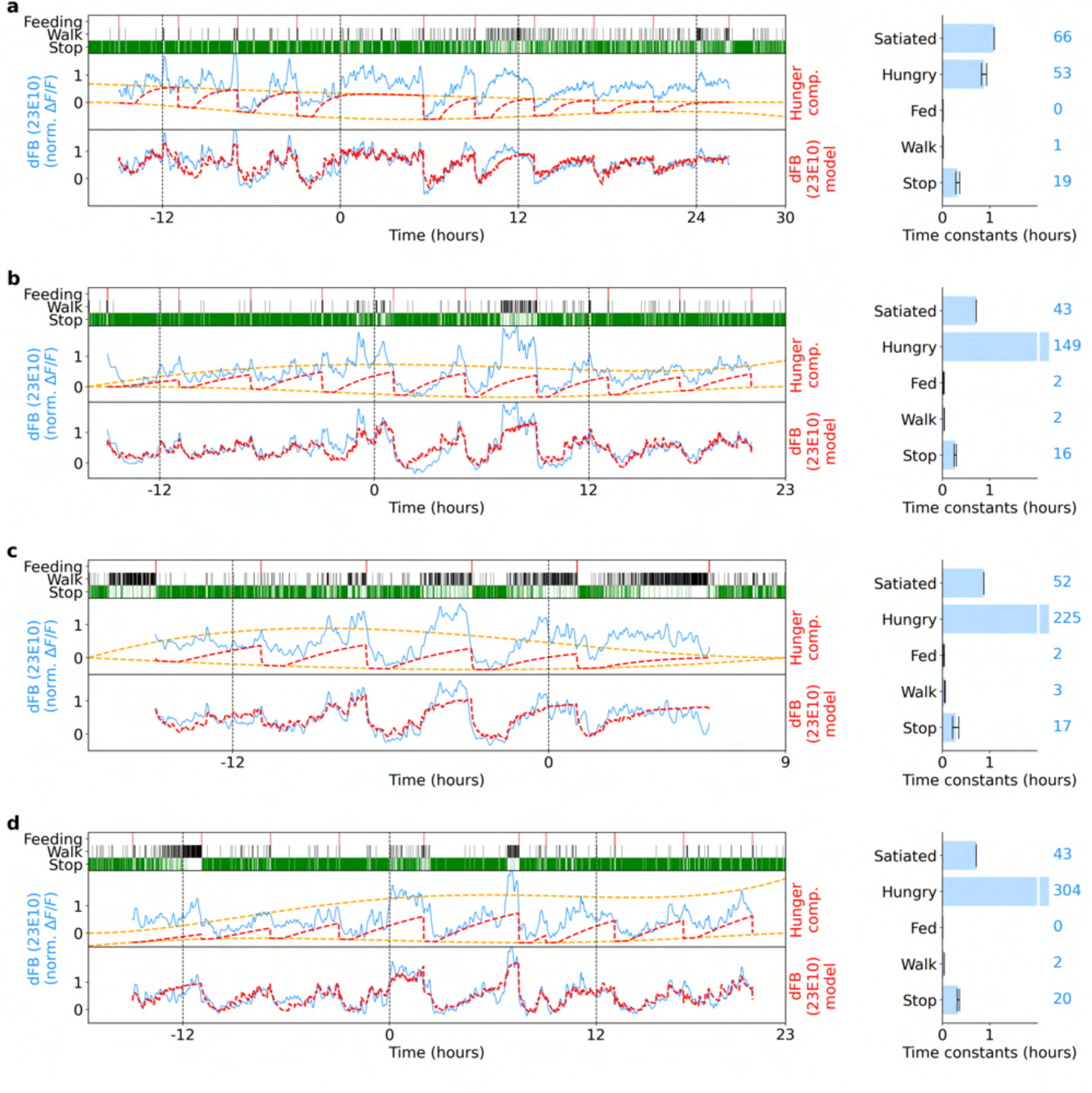
Fitting calcium activity of dFB neurons (labeled by GAL4-23E10) with hunger-walk model. **a** Left side: top row shows feeding events as well as walk and stop bouts of a fly. Second row: normalized fluorescence of dFB neurons in blue. Red line shows hunger component fitted by the model, while orange lines represent fitted upper and lower bounds of the hunger component. Third row: normalized fluorescence of dFB neurons in blue and fitted hunger-walk model in red. Right side: fitted time constants from the model. Grey lines indicate error bars of estimated time constants (see Methods), while colored numbers show their rounded fitted value. **b-d** Same as a. Each panel represents a fitted model for a different fly.

**Figure S52.**
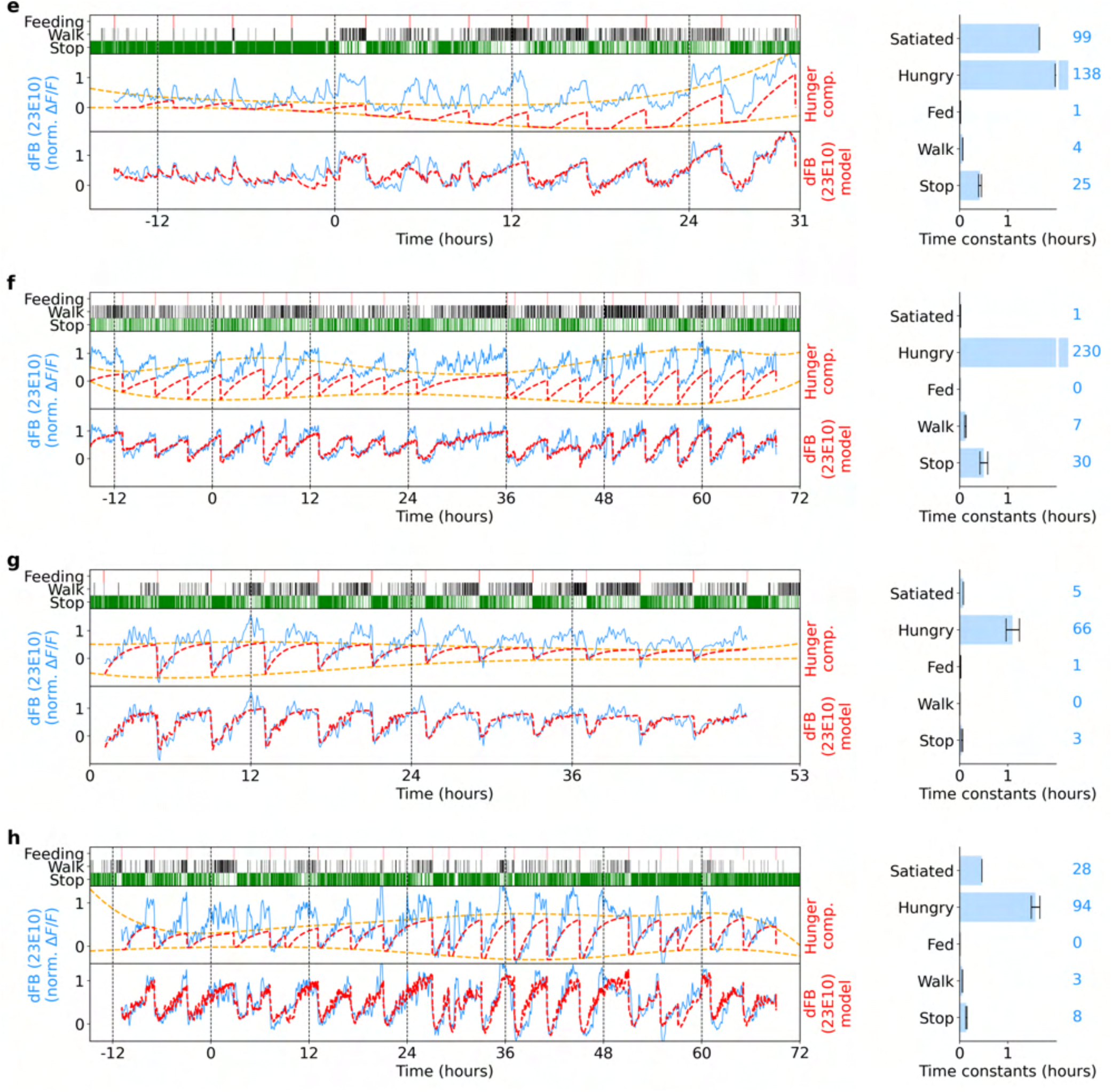
Another four experiments with dFB neurons and fits of hunger-walk model to calcium activity (labeled by 23E10-GAL4). Same as Supplementary Fig. S51.

**Figure S53.**
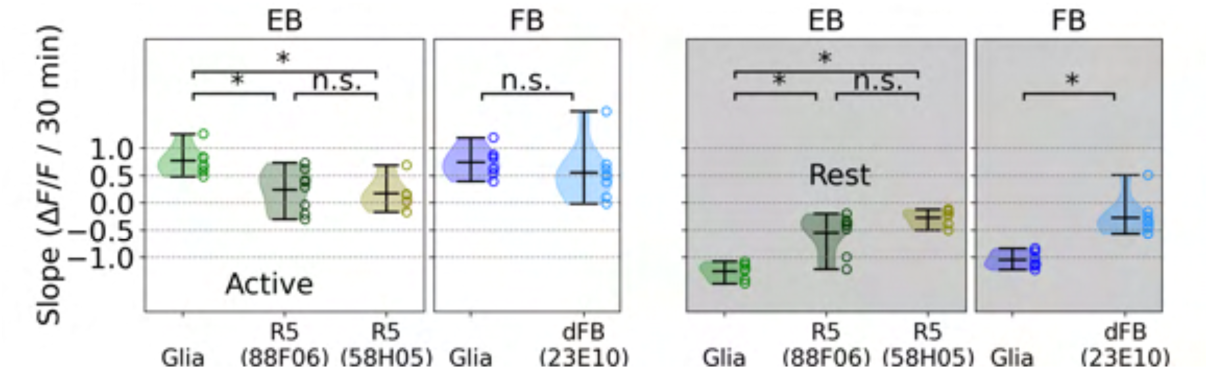
Left side: slope of a linear fit between time that flies spent in active epochs and average fluorescence traces from glia and neurons in EB and FB. Right side: same as left side but during resting epochs.

**Figure S54.**
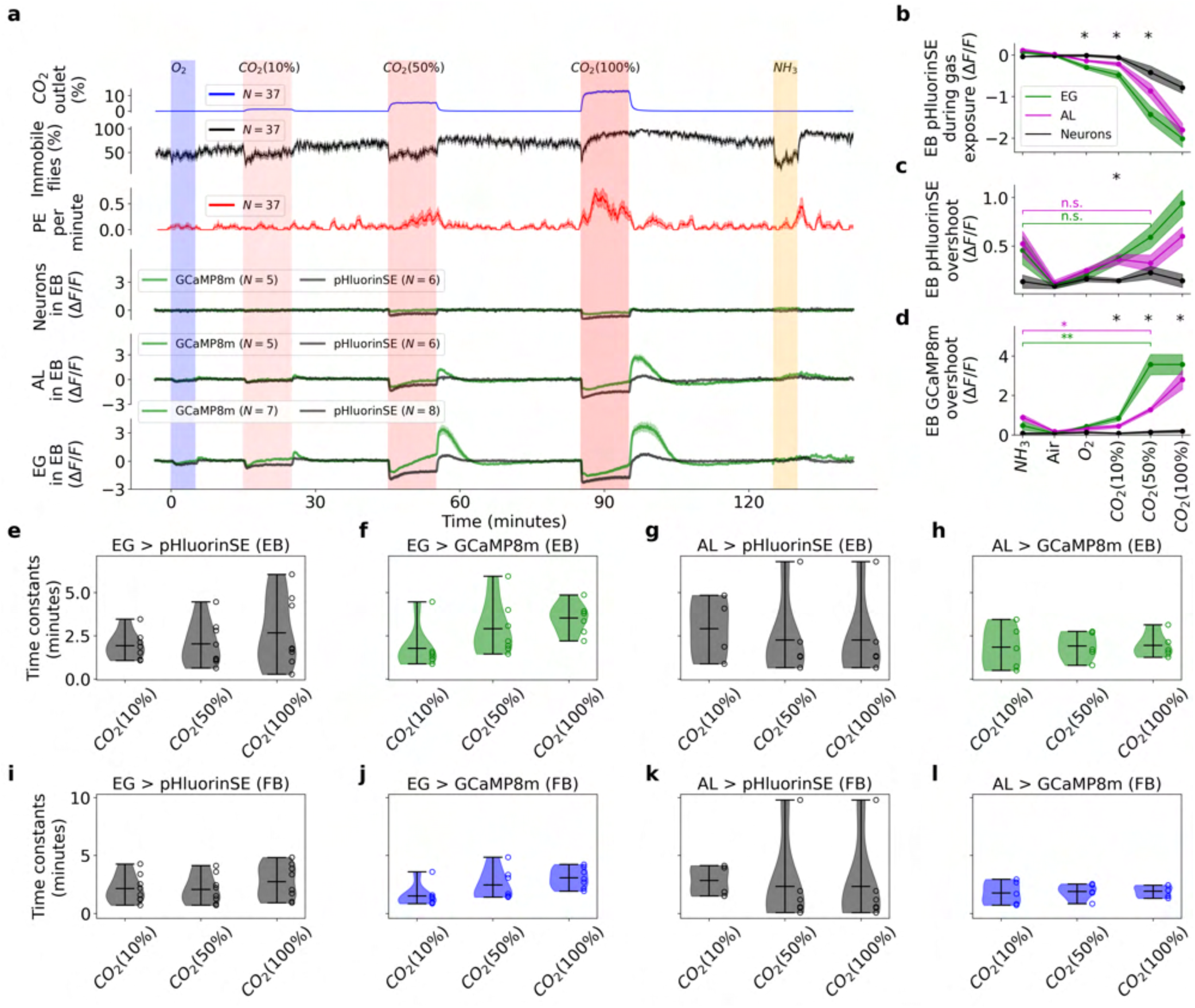
Analysis of CO_2_ / pH response in glia. **a** Same as Fig. 8f but for the EB. **b-d** Same as 8j-l but for the EB. **e-h** Fitted time constants for the decay after CO_2_ exposure in the EB for ensheathing glia (EG) expressing pHluorinSE and jGCaMP8m, and astrocytes (AL) expressing pHluorinSE and jGCaMP8m, respectively. **i-l** Same as e-h but in the FB.

**Figure S55.**
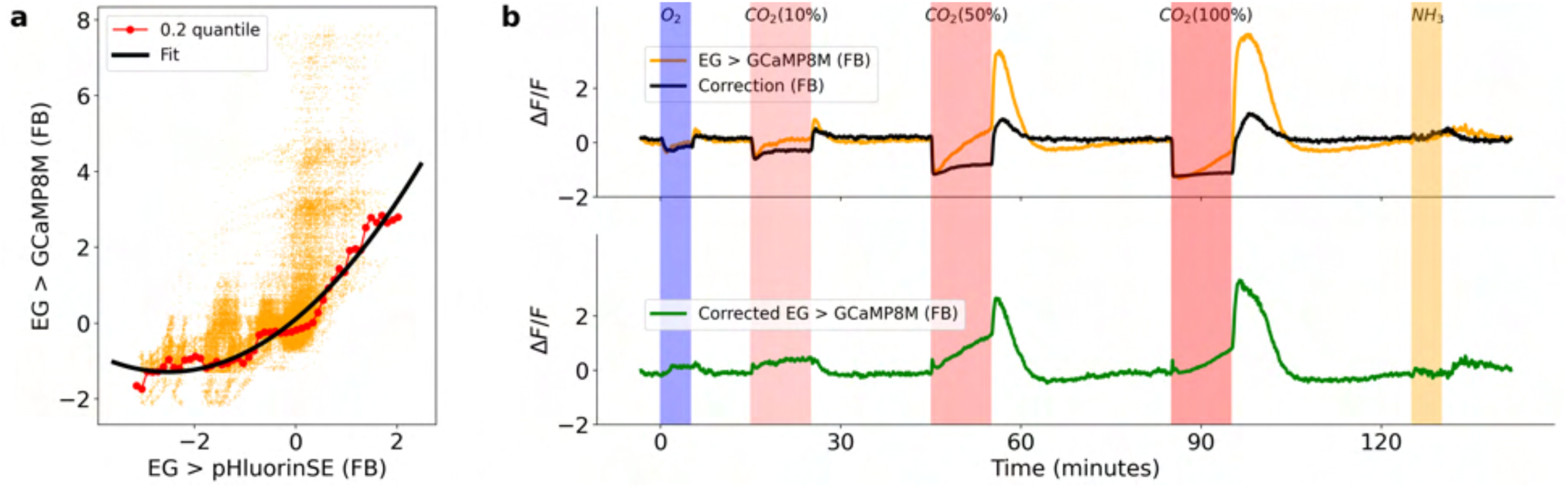
jGCaMP8m correction during of CO_2_ / pH response in glia. **a** pHluorinSE versus jGCaMP8m expressed in ensheathing glia in the FB (orange) during the CO_2_ exposure experiment (Fig. 8f). The red points indicate the 0.2 quantiles of jGCaMP8m values for each pHluorinSE bin (see Methods), while the black line is a polynomial fit of the red points. **b** On the first row, average jGCaMP8m in the FB of EG (orange) and jGCaMP8m correction (black) obtained from a. The second row shows the corrected jGCaMP8m signal (green), obtained from the difference between the recorded jGCaMP8m (green, first row) and the correction (black, first row).

**Figure S56.**
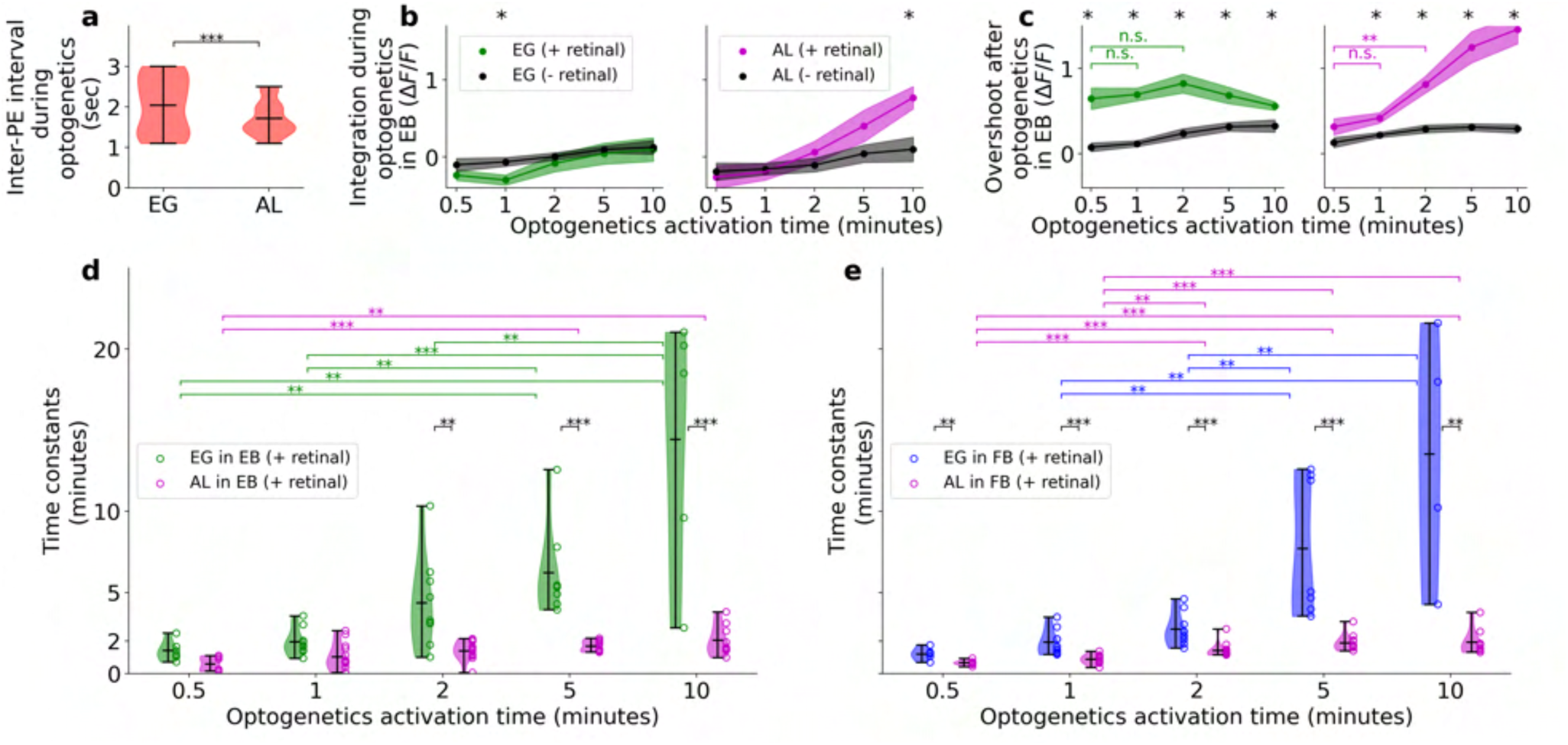
Analysis of glia dynamics during optogenetics activation. **a** Time between consecutive PEs (period) during optogenetics activation of EG and AL. Statistical significance was extracted using the Kolmogorov–Smirnov test (p < 0.0005)**b** and **c** same as 9f and g, but for the EB. **d** Fitted time constants from the calcium decay in EG (green) and AL (magenta) after each optogenetic activation experiment in the EB. **e** Same as c but for the FB where EG is now in blue. Asterisks indicate statistical significance using the t-test between EG and AL (black), and between different optogenetics activation times in EG (green) and AL (magenta). One, two, or three asterisks indicate statistical significance for p < 0.05, p < 0.005, or p < 0.0005, respectively.

**Figure S57.**
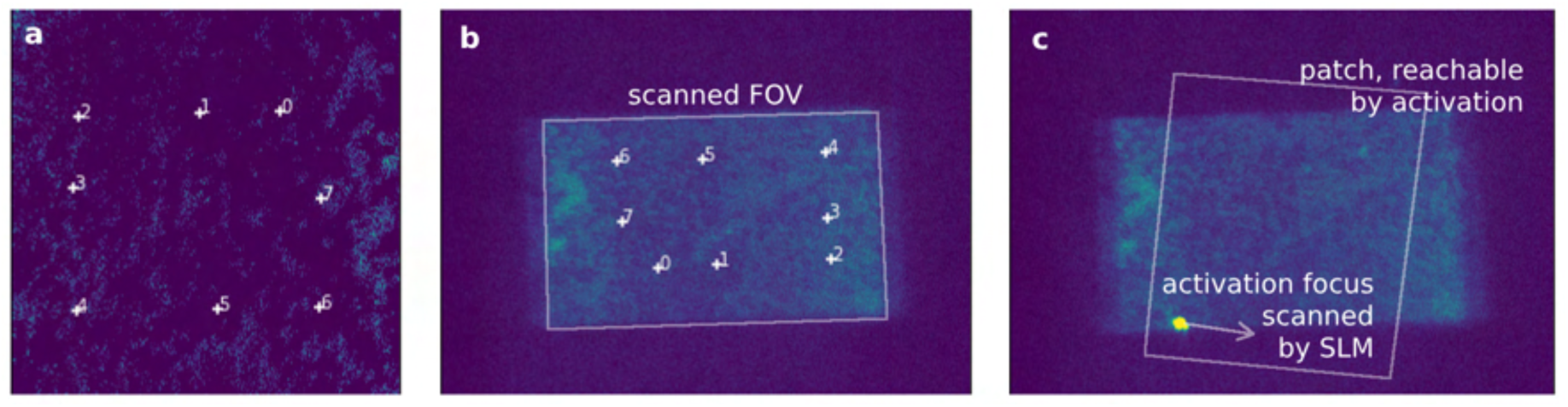
Images of fluorescent beads used for optogenetic activation calibration: **a** 2-photon image of the beads, **b** area camera view of the sample, illuminated by scanning beam (the exposure of the camera is longer than scanning period), **c** area camera view of the scanned sample, with optogenetic laser focused in one spot. Calibration procedure: 2-photon images (a) are taken simultaneously with the area scan camera image (b), and several prominent features are manually corresponend in both images. Transform matrix is found between pixel coordinates of the 2-photon and area camera image. Next, k-space is sampled with SLM by displaying phase ramps with varying spatial frequencies, sampling the k-space in x- and y-directions from -0.5 to 0.5 inverse pixels. Corresponding foci locations are found in area camera images (c), and the second transform matrix is calculated, linking area camera pixels to the SLM coordinates. The bounds of the patch, illuminated by the SLM are found. The final transform matrix is obtained by chaining both found transforms, linking 2-photon image coordinates to the coordinates of the optogenetic activation illumination patch. This transform is applied to any activation pattern drawn on 2-photon image, to obtain activation pattern in coordinates linked to the SLM, thus allowing to run Gerchberg-Saxton algorithm on it and yield the corresponding phase hologram.

